# High Resolution Biological and Physical Sampling Reveals Expression of Domoic Acid Biosynthetic Genes at Frontal Zones

**DOI:** 10.1101/2023.10.19.562961

**Authors:** Monica Thukral, Allegra T. Aron, Ariel J. Rabines, Daniel Petras, Christina M. Preston, Hong Zheng, Zoltan Fussy, Chase James, William Ussler, Andrew J. Lucas, Clarissa R. Anderson, Christopher A. Scholin, Pieter C. Dorrestein, John P. Ryan, Andrew E. Allen

## Abstract

Ocean microbes are the foundation of marine food webs, regulating carbon cycling and ecosystem dynamics. How they proliferate, die, move, and interact is regulated by physical, chemical, and biological factors that are dynamic and challenging to quantify in the natural environment. A significant limitation in many marine field studies is the inability to continuously sample the ever-changing ocean environment over space and time. In this study, we integrated spatiotemporal and multi-omic sample collection in an intensive sampling effort of phytoplankton ecology in Monterey Bay, California during the spring of 2021. Sampling methods coupled: (1) manual shipboard CTD sampling, (2) autonomous sampling using a Long-Range Autonomous Underwater Vehicle (LRAUV) equipped with an Environmental Sampling Processor (ESP), and (3) high-resolution physical measurements by an autonomous vertical profiler (Wirewalker). Sampling occurred as upwelling waned alongside declining domoic acid (DA) and low abundances of toxigenic *Pseudo-nitzschia*. Conditions needed to spark a widespread and toxic *Pseudo-nitzschia* bloom were absent, yet low-level DA was driven by similar mechanisms to those causing elevated DA. Three DA biosynthetic intermediate molecules were reported in the environment for the first time. Both shipboard and ESP sampling approaches identified DA biosynthetic gene expression at frontal zones. DA and expression of *dabA*, the gene encoding the first committed step of DA biosynthesis, were higher in association with recently upwelled water that supplied nutrients for growth and DA biosynthesis. Detection of subtle variations in *dab* gene expression in response to environmental variation provide a window into the ecological dynamics underpinning major toxic events.

**Graphical Abstract:** 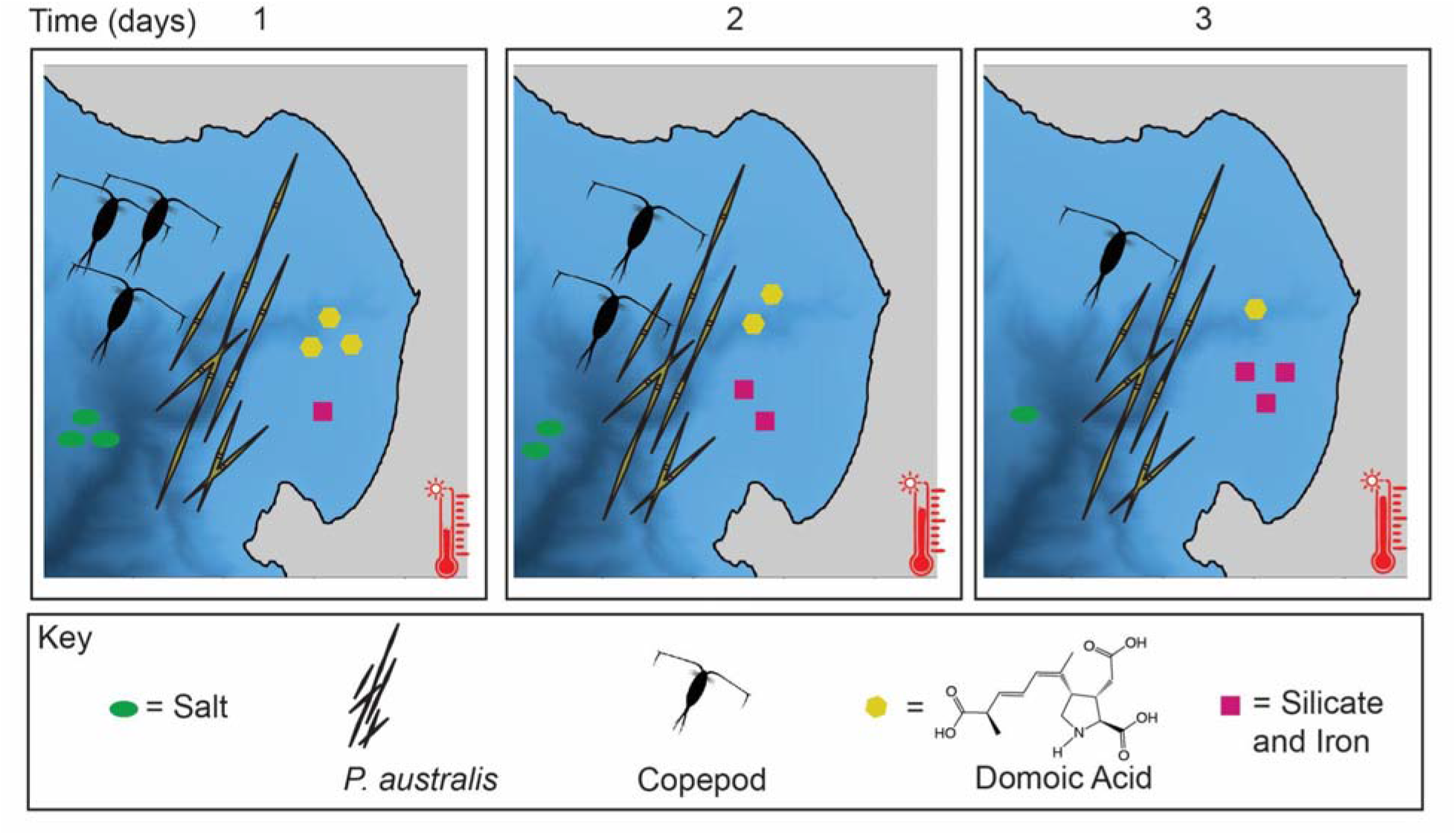

## Introduction

Microbes are essential in the regulation of marine food webs and elemental cycling, and are therefore important ecologically and economically [1]. Microalgae in coastal regions contribute significantly to marine primary productivity, modulate biogeochemical cycling, and are critical in climate models [2]. Wind-driven upwelling delivers nutrients to nearshore waters in spring and summer months in the California Current System. These coastal waters are often advected offshore during strong or prolonged episodes of upwelling, leading to spatial and temporal heterogeneity in conditions favorable to phytoplankton growth [3, 4]. This dynamic population of microbes generates and cycles short-lived organic molecules dissolved in seawater that make up a quarter of photosynthesis-derived carbon on the planet [5]. Due to the highly bioavailable nature of most dissolved molecules and the limited computational ability to annotate spectra as molecules in non-targeted metabolomics, much remains to be explored to understand the “microbe-metabolite network” in the ocean [5].

During these dynamic oceanographic conditions, harmful algal blooms (HABs) can thrive, especially in coastal regions. HABs threaten human health and coastal economies by producing toxic molecules that can harm and cause detrimental fisheries closures. The *Pseudo-nitzschia* diatom genus is a well-known HAB-former worldwide and often forms blooms in response to elevated nutrient levels in coastal upwelling zones and at the sea-ice interface [6]. Some species of *Pseudo-nitzschia* biosynthesize an irreversible neurotoxin domoic acid (DA) that can act as an ionotropic glutamate receptor agonist, causing cellular excitotoxicity and apoptosis [7]. DA bioaccumulates in the food web and can cause mass mortality to marine mammals and birds, and to humans if they consume DA-containing shellfish or crustaceans [8]. Exposure to DA can cause amnesic shellfish poisoning (ASP) which causes seizures and vomiting among other adverse health outcomes [9, 10]. The DA biosynthetic gene cluster (*dab*) was recently discovered with *in vitro* characterization of three DA biosynthetic genes and their biosynthetic intermediate molecules [11]. Among these, *dabA* encodes a terpene cyclase that combines glutamate and geranyl pyrophosphate to form N-geranyl-L-glutamic acid (L-NGG). The product of *dabD*, a Cytochrome P450 enzyme, oxidizes L-NGG to 7’-carboxy-L-NGG. Finally, a dioxygenase encoded by *dabC* cyclizes the compound into isodomoic acid which is hypothesized to isomerize to DA.

While blooms of *Pseudo-nitzschia* occur annually along the North American west coast due to seasonal upwelling [12, 13], they do not always result in DA production or negative ecosystem impacts. *Pseudo-nitzschia australis,* a particularly toxic species, was primarily responsible for an unprecedented toxic bloom off the west coast of North America in the spring of 2015 [14, 15]. The vast geographic expanse of the bloom has been attributed to the 2014-2016 marine heatwave that allowed for *Pseudo-nitzschia* range expansion. Despite anomalously warm temperatures, strong upwelling persisted in a nearshore coastal band in central California, thereby continually fueling the *Pseudo-nitzschia* bloom with nutrient-replete source waters, which caused cellular silicic acid (Si)-limitation, a potential driver for the elevated increased DA production [8, 14]. Si-limitation can also be mediated by iron-limitation in, which would likely influence the magnitude of DA production during upwelling-relaxation transitions [16, 17]. Top-down controls are also gaining momentum in studies of DA production as a potential strong driver that may complement or act as a multiplier to bottom-up processes. Although there are conflicting results on how copepods affect DA production, recent work has shown that *Pseudo-nitzschia* exposed to copepods produced more DA, suggesting that copepods affect cellular toxicity [18–21]. Furthermore, the copepodamide suite of molecules were characterized as chemical cues that cause increased DA production [22].

Historically, Monterey Bay experiences *Pseudo-nitzschia* blooms annually with inter-annual variability of intensity and toxicity, driven by discrepancies in seasonal upwelling originating at Pt. Año Nuevo and Pt. Sur [15, 23–25]. Increased upwelling has been associated with toxic *Pseudo-nitzschia* blooms along the California coast. To monitor such harmful events, routine sampling is conducted at coastal piers in Monterey Bay and elsewhere, in addition to comprehensive rapid response sampling of large-scale toxic bloom events [26–28]. However, few studies have examined factors that influence *Pseudo-nitzschia* ecology during low compared to high abundance [29, 30].

In this work, we explored the ecological dynamics of *Pseudo-nitzschia* during a non-DA-outbreak state as an important end member to extreme conditions. We present one of the first synchronous studies of Monterey Bay using multiple sampling methods (Eularian, Lagrangian, and Quasi-Lagrangian) and assets (Wirewalker, shipboard CTD/Niskin, and ESP/LRAUV) to support a multi-‘omic study of the microbial ecology of HABs. A Lagrangian sampling method follows the same mass of water as it moves spatially over time, while a Eulerian sampling method samples a fixed location over time. We show that the principal environmental inducers DA production, such as silica limitation and grazing, are similar during non-DA-outbreak and DA-outbreak events. *Dab* gene expression was identified at frontal zones, and expression of *dabA* was elevated on the high salinity side of fronts, concurrent with recently upwelled water. In addition, this study associates *dab* gene transcription with DA production, providing insights into how microbial metabolism is linked at low abundance and toxicity levels. The results of these studies provide fundamental science to inform environmental monitoring of HABs.

## Materials and Methods

Full methods and raw data can be accessed in the Supplementary Information. Briefly, ship-CTD sampling was conducted over three days: 20-22 of April 2021 in Monterey Bay, California using a Conductivity Temperature Depth (CTD) rosette with Niskin bottles to collect water at the surface, subsurface chlorophyll maximum (SCM), and deep depths (**Fig. 1a**). Water samples (filtered and whole) were used for metabarcoding (16S, 18SV4, 18SV9, ITS2), metatranscriptomics, metabolomics, macronutrients, and chlorophyll. Stations were selected to provide a coupled quasi-Lagrangian and Eulerian time series. Shipboard samples at three fixed locations daily where moored Wirewalkers operated continuously during April 17-24 – stations “N1”, “E1”, and “S1”--created the Eulerian time series. The quasi-Lagrangian time series locations were determined by drifting with a robotic fleet consisting of a Waveglider and a 3G-ESP/LRAUV. The ship collected samples at four stations, 2-km North, South, East and West of the drifting vehicle (**Fig. 1b**). Between April 20-25 the LRAUV/ESP drifted within a water mass with high chlorophyll biomass as it moved offshore, and the ESP collected and preserved samples regularly to create the quasi-Lagrangian time series (**Fig. 1c**).

**Fig. 1.**
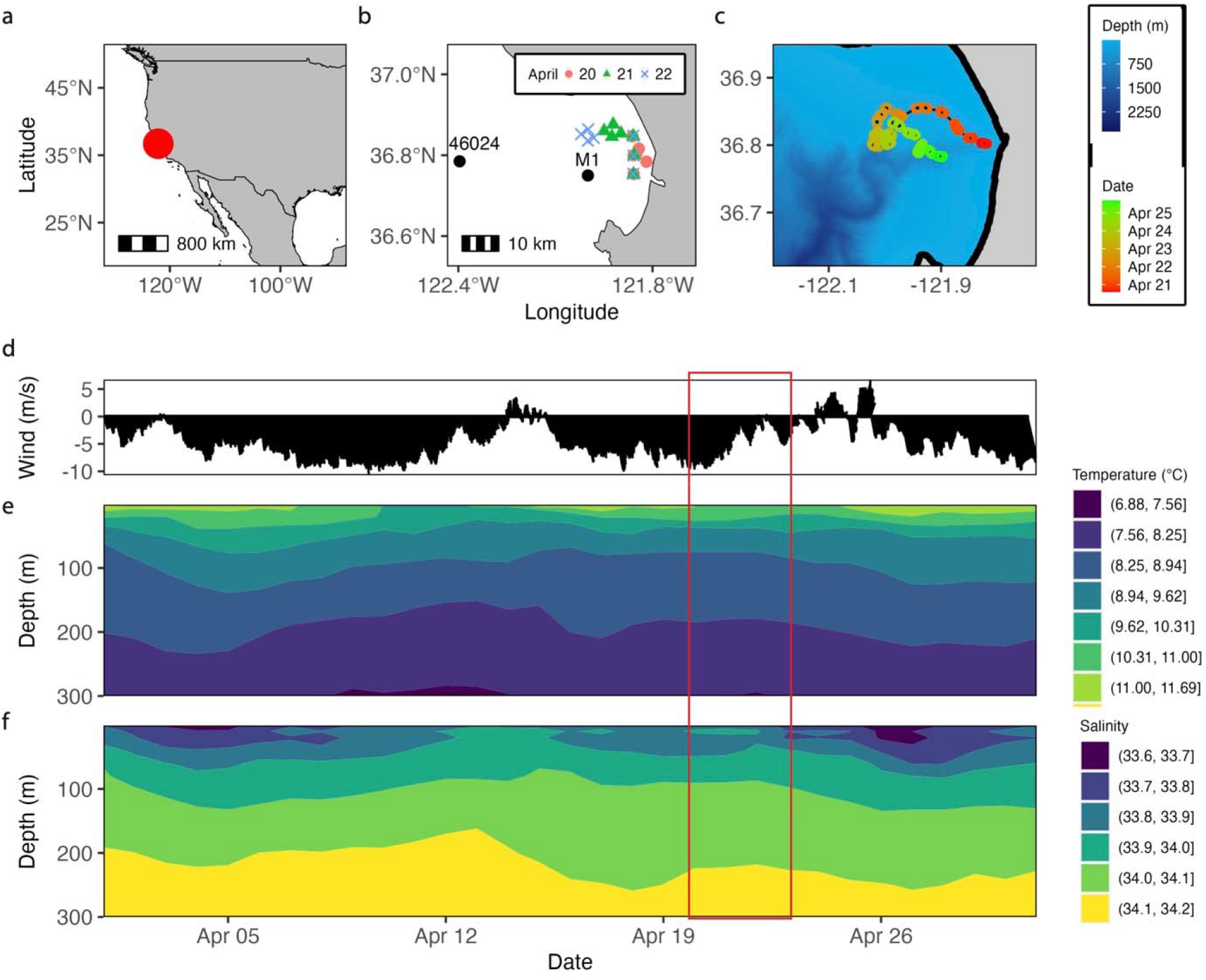
Environmental Context and Regional Upwelling Conditions of Monterey Bay, CA, USA during April of 2021. (a) Map with red dot indicating Monterey Bay, CA, USA, (b) Ship CTD sampling map and observing locations, and the (c) 3G-ESP/LRAUV path and bathymetry bottom depth (m). Colored and shaped symbols in (b) indicate each of the three days of shipboard sampling taken on April 20, 21, and 22. CTD casts were taken at three depths (surface, SCM, and deep) at all stations indicated. Colored symbols in (c) indicate the day and time the LRAUV reached the location indicated. (d) wind speed from National Data Buoy Center (NDBC) Station 46042; negative values are upwelling favorable. (e) Daily mean temperature (°C) and (f) daily mean salinity from MBARI Mooring M1. The ship sampling period is indicated with a red box (April 20-22, 2021).

Metabarcoding data from ship samples were generated by preparing amplicon libraries of 18SV9, 18SV4, 16S, and ITS2 from DNA extracted from filtered CTD seawater. Metatranscriptomes of cDNA were generated from ribosomal RNA-depleted total RNA extracted from filters collected by the 3G-ESP. Amplicon libraries of 18SV9, 16S, and ITS2 from the 3G-ESP were prepared by amplifying this cDNA.

DA was measured with three methods: (1) real-time detection of particulate DA using the Environmental Sample Processor (ESP) with an Surface Plasmon Resonance (SPR) instrument on the LRAUV that sampled within the tracked water mass (Ussler, unpublished); (2) whole water sampling from a CTD-Niskin rosette for solid phase extraction and liquid chromatography coupled to tandem mass spectrometry (LC-MS/MS) untargeted metabolomics analysis to measure total DA (tDA); and (3) extractions of filtered sea water from CTD Niskin-rosette to measure particulate DA (pDA) using LC-MS/MS targeted metabolomics [31, 32].

## Results

Due to the dynamic nature of regional wind-driven circulation, different water types are continuously stirred and mixed in Monterey Bay [24]. The endmember water types sampled in this study are upwelled waters that are relatively rich in nutrients and, and resident bay waters that are relatively rich in plankton (**Fig. S1a, S1b, S1c, S1d**) [14, 33]. At the two northern stations (N1 and E1) on the day of sampling, nutrients were relatively high at the surface at >25 µM silicic acid, >20 µM nitrate, >1.8 µM phosphate. These nutrients supported the phytoplankton biomass (range chlorophyll-*a* 0.108-9.316 µg/L, mean 2.036 µg/L) (**Fig. S1e, S1f**). Despite the California central coast region’s characteristic iron stress, most silicic acid: nitrate (Si:N) ratios measured were >1.0 (range Si:N = 0.66-37.96, mean Si:N = 3.76), and were not associated with silicic acid drawdown resulting from diatom iron limitation [4, 34, 35] (**Table S1**). In addition, the Si_ex_ proxy for diatom iron limitation [36, 37] averaged greater than 1.0 (mean Si_ex_ = 4.53), generally suggesting minimal impact of diatom iron limitation. Analysis by Spearman’s rank correlation between environmental variables reflected significant positive correlations between temperature and chlorophyll, and salinity and dissolved nutrients (**Fig. S1g**).

### Environmental Context and Regional Upwelling Conditions

Regional upwelling was strongly developed by the April study period, as measured by the Biological Effective Upwelling Transport Index (BEUTI) and measurements of surface winds and temperature and salinity in the water column (**Fig. 1d, Fig. S1h**). In the week before sampling, upwelling-favorable winds caused a steady increase in upwelling at Pt. Año Nuevo, which caused outcropping of salinity isohalines and temperature isotherms at M1 (**Fig. 1e, 1f**). By 14 April, the persistent upwelling favorable winds relaxed rapidly, and mixing of recently upwelled water with resident bay water was evident in Wirewalker and LRAUV observations. The Bay experienced a counterclockwise flow of water, as indicated by high frequency radar measurements of surface currents (cordc.ucsd.edu), which is a typical flow pattern [38]. During the ship sampling period (April 20-22), upwelling had weakened, but the biological community likely still experienced upwelled water introduced to the Bay during the preceding period of strong upwelling. The quasi-Lagrangian path of the drifting LRAUV revealed the resulting circulation of the central bay, as waters circulated in a counterclockwise flow.

### Eulerian Analysis with Moored Wirewalkers Discloses Different Water Masses

Wirewalkers were moored at N1, S1, and E1 in water depths of 35-, 55-, and 250-meters, respectively [39] (**Fig. 2a**). Diel patterns associated with vertical movement of the internal tide layer are captured twice daily by all Wirewalkers in fluorescence and light measurements (**Fig. S2a)**. Among the moorings, N1 observed the freshest, warmest water, and lowest biomass. Day 1 at N1 was characterized by intermediate salinity, consistent with aged, upwelled water that had mixed with coastal transition zone water and Bay water and contained a stratified surface mixed layer. An LRAUV survey captured the influx of high-salinity, recently upwelled water entering the study area immediately before the start of ship sampling (**Fig. S2b)**. By Day 2, N1 observed strong destratification indicated by the disappearance of the low salinity signature due to either (a) local wind-driven turbulent mixing (b) advection of a front past the mooring, or (c) nighttime cooling in conjunction with wind mixing from a cold front **(Fig. S2c).** This destratification was observed at all three moorings. Mooring E1 observed homogenous chlorophyll at the surface and internal tides that brought high chlorophyll water to depths below 100 meters. Dissolved oxygen patterns revealed near-surface increases in oxygen concentrations, suggesting productivity-related oxygen increases.

**Fig. 2.**
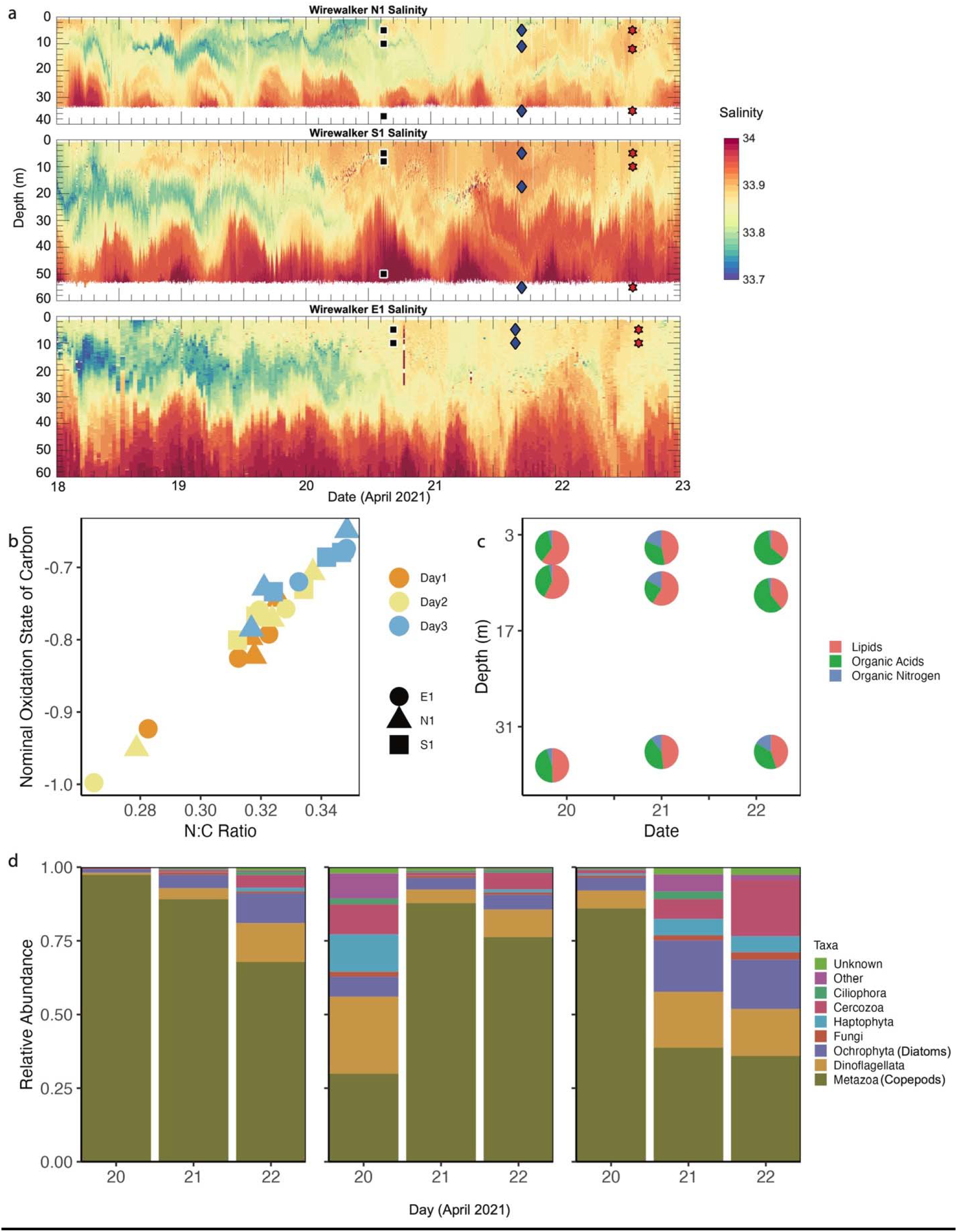
(a) Salinity profiles from each Wirewalker instrument over three days, in which reds indicate higher salinity and blues indicate lower salinity. Ship-sampling times are indicated with the overlaid symbols. (b) Weighted average nominal oxidation state of carbon (NOSC) of metabolite features vs weighted average elemental Nitrogen to elemental Carbon ratio (N:C) of metabolite features captures a shift toward less labile dissolved organic matter. (c) The three most abundant molecule superclasses at station N1 were lipids, organic acids, and organic nitrogen compounds. (d) Select taxa by 18SV9 metabarcoding at station N1 at three depths, (left) surface, (middle) SCM, and (right) deep.

Discrete CTD samples showed differential physical properties and nutrient conditions between the moorings (**Fig. S2c, S2d**). E1 had the highest absolute nutrients likely due to internal tidal nutrient transport over Monterey Canyon. N1 showed a unique pattern of reduced Si-limitation over time as indicated by the increasing Si:N ratio. The weighted average elemental Nitrogen to elemental Carbon ratio (N:C) vs. Nominal Oxidation State of Carbon (NOSC) of total metabolite features at the moorings showed an increase in N:C and NOSC over the three days, likely due water mass mixing (**Fig. 2b**). A less negative NOSC value is indicative of more potential energy gained during catabolism and a higher N:C ratio is indicative of molecules with relatively more nitrogenous groups such as amines and amides [40]. The lowest N:C and NOSC occurred at S1, while those of N1 and E1 were more widely distributed. Neither N1 nor S1 had a distinct SCM on the second and third days, despite elevated but decreasing chlorophyll over time.

This mixing signature was mirrored by the similarity in surface and SCM metabolomes at N1 (**Fig. 2c**). The most abundant molecule superclasses at N1 were lipids, organic acids, and organic nitrogen compounds [41, 42], and there was a distinct shift on Day 3 at N1 surface and SCM from lipid-dominated to organic acid-dominated compounds. In the deep samples, the amount of organic nitrogen compounds increased over time. Changes in chemical composition over time at N1 may have been related to changing nutrient availability and increased mixing. N1 also experienced a shift in community composition over time: surface and deep samples showed an increase in the dinoflagellate to copepod ratio and diatom to copepod ratio over time (**Fig. 2d**). There were proportionally more dinoflagellates and diatoms at the surface over time, in agreement with low nutrient levels. The lowest chlorophyll-*a* and the highest tDA at N1 were at the surface on the first day (**Fig. S2e, S2f**). Interestingly, this tDA measurement was the highest in the entire study.

### *Pseudo-nitzschia* Biosynthesizes Low Levels of Domoic Acid

Unlike the spring of 2015 when a bloom dominated by *Pseudo-nitzschia australis* caused a highly toxic HAB event, the spring of 2021 was not characterized by abundant *Pseudo-nitzschia* diatoms nor copious, acute levels of DA toxin. 18S V9 rRNA gene amplicon metabarcoding revealed *Pseudo-nitzschia* as the third most abundant diatom genus (**Fig. 3a**). ITS2 amplicon sequencing revealed twelve *Pseudo-nitzschia* species in ship samples; of those detected, *australis, seriata, and multiseries* are some of the most robust DA-producers, but made up only a small fraction of the community dominated by *P. americana* (**Fig. 3b**, **Fig. 3c**) [6, 43]. Samples taken at depth, about five meters above the sea floor, had the highest ITS2 alpha diversity (p<0.05) **(Fig. S3a**). ITS2 alpha diversity increased as 18SV9 alpha diversity increased with a linear relationship, suggesting that samples with a more diverse eukaryotic community also had a more diverse *Pseudo-nitzschia* community, though no community change was observed over time (**Fig. S3b, S3c**). tDA and *Pseudo-nitzschia*, as identified by 18SV9 metabarcoding, imperfectly overlap spatially (**Fig. 3c, 3d**). *Pseudo-nitzschia* abundance has been reported in other studies to be decoupled from DA concentration over time since the cellular quota of DA can vary [44].

**Fig. 3.**
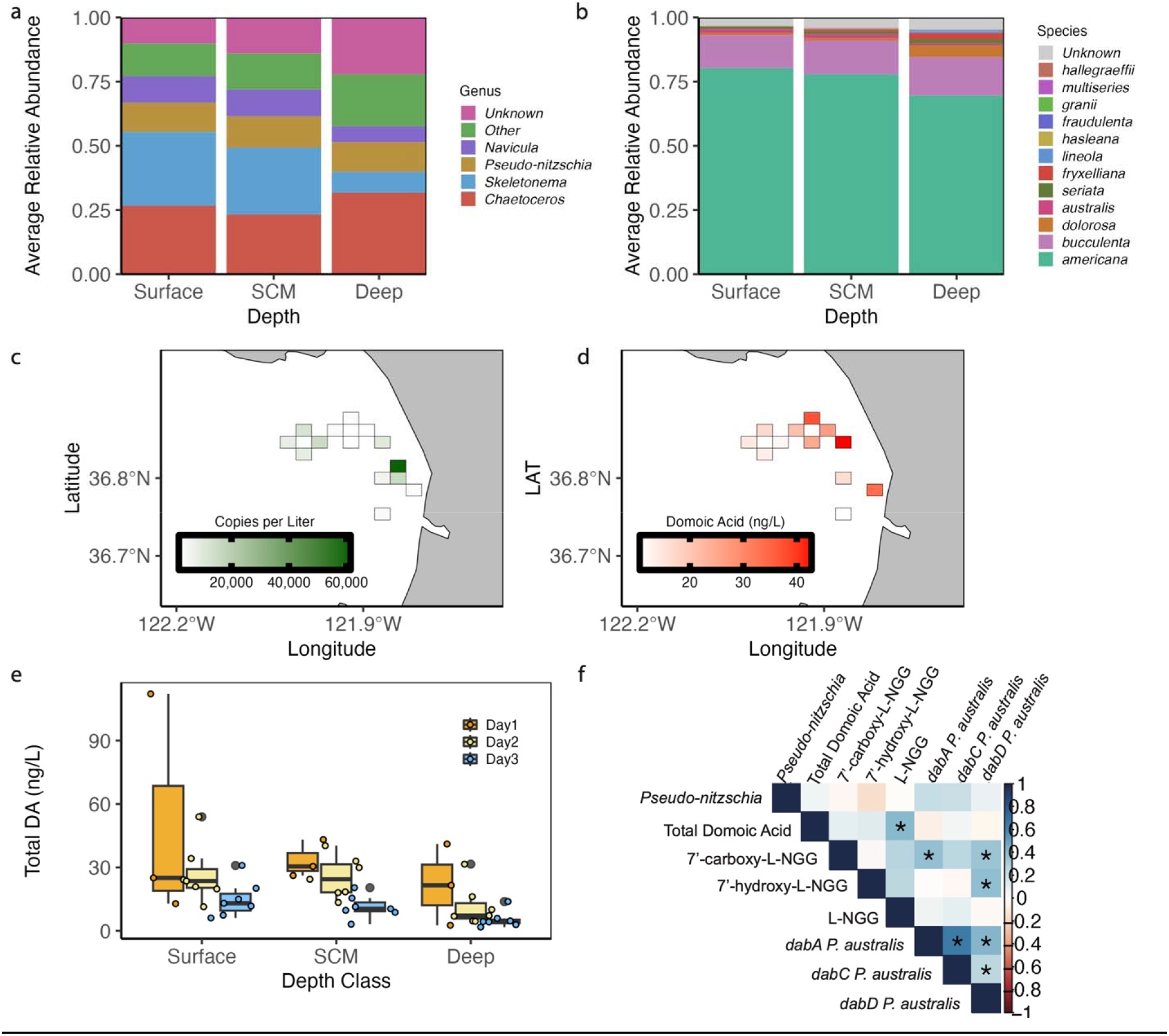
(a) 18SV9 rRNA gene sequencing of shipboard samples showed the four most abundant diatom genera are *Chaetoceros*, *Skeletonema, Pseudo-nitzschia,* and *Navicula*. (b) ITS2 sequencing of shipboard samples identified twelve *Pseudo-nitzschia* species, predominantly species *americana*. (c,d) *Pseudo-nitzschia* distributions (18SV9 copies/L) overlap spatially with total domoic acid (ng/L). The color bar in (d) shows peak area after blank subtraction and normalization. (e) Total domoic decreased each day in all three depth classes among all stations among all 52 shipboard samples. (f) Spearman rank correlations identified relationships within the domoic acid biosynthetic pathway from ship samples, where values closer to 1 (dark blue) are positive correlations, and values closer to −1 (dark red) are negative correlations, and an asterisk (*) represents significant correlations.

While all samples contained tDA at varying levels, tDA was highest nearshore and in the northeast region of the Bay (**Fig. 3d**). tDA was the highest on the first day of sampling and decreased daily in all three depth classes (**Fig. 3e**). Among the moorings, it is likely that tDA was highest at N1 because S1 and E1 were already mixed, whereas N1 retained upwelled water for longer. Notably, tDA in the spring of 2021 was two orders of magnitude lower than particulate DA (pDA) quantified in spring of 2015 during the historic DA event. Furthermore, DA biosynthetic intermediates were identified in the environment, and were also present in similar locations as tDA (**Fig. S3d**). Modified versions of DA were also detected, such as methyl-DA (*m/z* + 14.016) (**Fig. S3e)**. In shipboard data, *P. australis* expression of the DA biosynthetic genes *(dabA*, *dabC*, and *dabD)* were positively significantly correlated with each other, as were genes and the molecules they produce: *dabD* which is transcribed to produce the protein that makes 7’-hydroxy-L-NGG and 7’-carboxy-L-NGG was significantly positively correlated with those two molecules (**Fig. 3f**). Importantly, L-NGG showed the strongest correlation and significance with tDA, the first time this correlation has been reported in the environment (**Fig. 3f**). The implication is that L-NGG and *dabA*, the gene that encodes the enzyme that produces it, should be considered as informed targets for environmental monitoring and toxin prediction because they are produced prior to DA.

### Northeastern Bay Ecology Reveals the Decline of Domoic Acid

Nutrient influx from upwelled water in early April likely relieved nutrient limitation and promoted phytoplankton growth, which in turn stimulated grazing. Total DA and biomass-normalized tDA were the highest at N1 with a peak at the surface on Day 1, decreasing in tDA concentration as the water mass was advected offshore (**Fig. 4a, S4a**). Overall, tDA was elevated in higher temperature, lower salinity water, consistent with the highest temperature and relatively low salinity measured at N1 surface Day 1. These data support several hypotheses to explain the decreased tDA at the surface of N1 including decreases in Si limitation, iron limitation, and grazing pressure. Si:N and Si_ex_ were the lowest on Day 1 and increased over time, indicating an easing of Si and iron limitation (**Fig. 4b**). Notably, Si:N and Si_ex_ never became particularly low, yet their lowest levels coincided with the greatest tDA. The overall nutrient availability was higher on Day 1 and dropped quickly thereafter, so although Si:N increased, there was lower absolute nutrient abundance. Even though nutrient concentrations were not inherently limiting, the ratios suggest an imbalance that could induce cellular toxicity. At the surface of N1, the lowest Si_ex_ value was measured on Day 1 and increased over time, suggesting relatively less iron on the first day of sampling. Since this value was positive, however, true diatom iron limitation was not confirmed. At N1, 18SV9 rRNA abundances of copepods to *Pseudo-nitzschia* declined over time, suggesting decreased grazing pressure over time (**Fig. 4c**). In the whole dataset, higher copepod abundance was correlated with lower *Pseudo-nitzschia* abundance and higher tDA, and the highest copepod abundance was observed at the surface of N1 on Day 1, concurrent with the highest tDA (**Fig. 4d**).

**Fig. 4.**
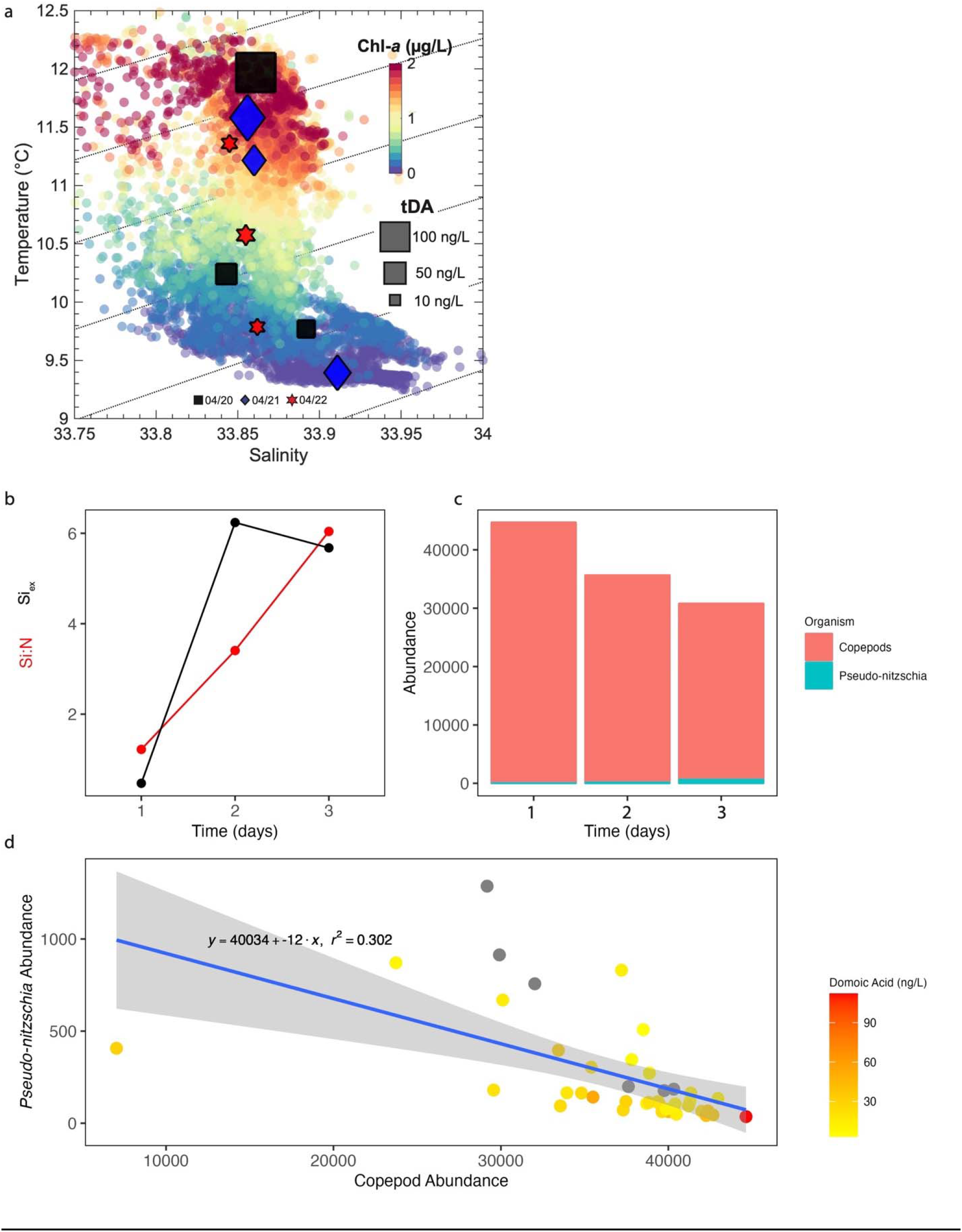
(a) Total domoic acid relative to temperature and salinity, as measured by discrete ship based CTD water samples near the Wirewalker at station N1. The color of the background dots, as indicated with the colorbar, represents the log_10_ of the relative chlorophyll fluorescence as measured by the Wirewalker stationed at N1. The colored shapes (black square, blue diamond, red star) overlayed represent the three days of sampling of total domoic acid (tDA), and the size of the shape represents the concentration of tDA. Curved dashed lines represent isopycnals, lines of constant density. (b) Iron limitation at N1 surface was measured by the Si_ex_ metric, a proxy for diatom iron limitation (black). The dissolved silicic acid to nitrate (Si:N) ratio at N1 surface increased over time (red). (c) The copepod to *Pseudo-nitzschia* ratio at the surface of N1, as determined by 18SV9 metabarcoding, decreased over the three days. (d) A linear relationship exists between copepod and *Pseudo-nitzschia* abundance in samples from the surface and SCM, and the sample from the N1 surface, indicated by the darkest red circle, showed the highest number of copepods and low *Pseudo-nitzschia*. Axes abundance values in (c) and (d) are read counts per sample, rarefied to even sequencing depth.

### Quasi-Lagrangian Analysis by 3G ESP/LRAUV Reveals *dab* Gene Expression **Patterns**

Water samples (n=24, depth range 1.5 to 14.7) collected by the robotic fleet as the water mass moved offshore coupled shipboard samples to investigate the expression of genes involved in DA biosynthesis. Gene expression of the DA biosynthetic gene cluster (*dabA, dabB, dabC, and dabD*) was detected in 16 samples with >90% nucleic acid identity to known sequences (**Fig. S5a, S5b, S5c, S5d, S5e**). Among both shipboard and ESP samples, *dab* was detectable only in the top 70 meters of the bay, which is concurrent with the euphotic zone where populations of *Pseudo-nitzschia* reside [11]. The highest gene expression of *dabA*, which initiates the committed biosynthesis of DA, was found at N1, where the highest tDA was also measured. Spearman’s rank correlation analysis was conducted between the *dab* genes, biosynthetic intermediate molecules of DA, and *Pseudo-nitzschia* abundance. Significant positive correlations were observed in gene expression of: *dabA* with *dabC*, *dabA* with *dabD*, and *dabC* with *dabD* (p<0.05) (**Fig. S5f**). Furthermore, significant positive correlations between tDA and L-NGG abundance, 7’-carboxy-L-NGG and *dabA,* 7’-carboxy-L-NGG and *dabD*, and 7’-hydroxy-L-NGG and *dabD* were observed.

## Discussion

Our data suggest that *dab* gene expression increases with response to recently upwelled waters containing freshly upwelled nutrients (**Fig. 5a**, **Fig. 5b**). *P. australis* expression of *dabA* varied along a hydrographic gradient, such that higher expression levels accompanied higher-salinity recently upwelled water, across both the ship and 3G-ESP/LRAUV platforms. Downstream genes *dabC* and *dabD* from *P. australis* were measured in relatively fresher waters, indicating expression in more aged, upwelled waters. Furthermore, temperature and salinity measurements from the LRAUV over time indicate that *dab* gene expression peaked at physical transitions and frontal features where different water types meet and mix, identified by abrupt changes in temperature and salinity reflecting boundaries of recently upwelled water and resident bay water (**Fig. 5c**). This same association was indicated in the Wirewalker time series, which showed the highest DA detection where a water type boundary transition was detected at N1. Nutrient pulses at fronts may relieve nitrogen limitation, allowing for growth and production of the nitrogen-rich metabolite DA, under sustained silica-limitation. The California Current Ecosystem (CCE) experiences oceanic fronts in the spring and summer upwelling season that contain elevated microbial and grazing activity and organic matter [45–48]. N1, where the highest tDA was measured, also contained relatively high *dabA* from *P. australis*, in the upper 10% of ship samples, and the highest salinity measurements. Even though upwelling was weak during the sampling period, strong persistent upwelling in the Bay earlier in the month brought upwelled water to station N1 where *dabA* and tDA were maximal. We hypothesize that *dab* expression peaked at the high salinity side of physical transition zones because *Pseudo-nitzschia* cells were exposed to nutrients from recently upwelled water, stimulating cellular processes. Several samples showing elevated *dab* expression coincided with where the LRAUV drift reversed course in a counterclockwise loop, consistent with influx, stirring, and mixing of regional water types **(Fig. S5g).** Perhaps, under DA outbreak conditions, such *dab* expression and DA production could be further aggravated by a combination of sustained Si limitation, iron limitation and grazing that did not manifest in this study.

**Fig. 5.**
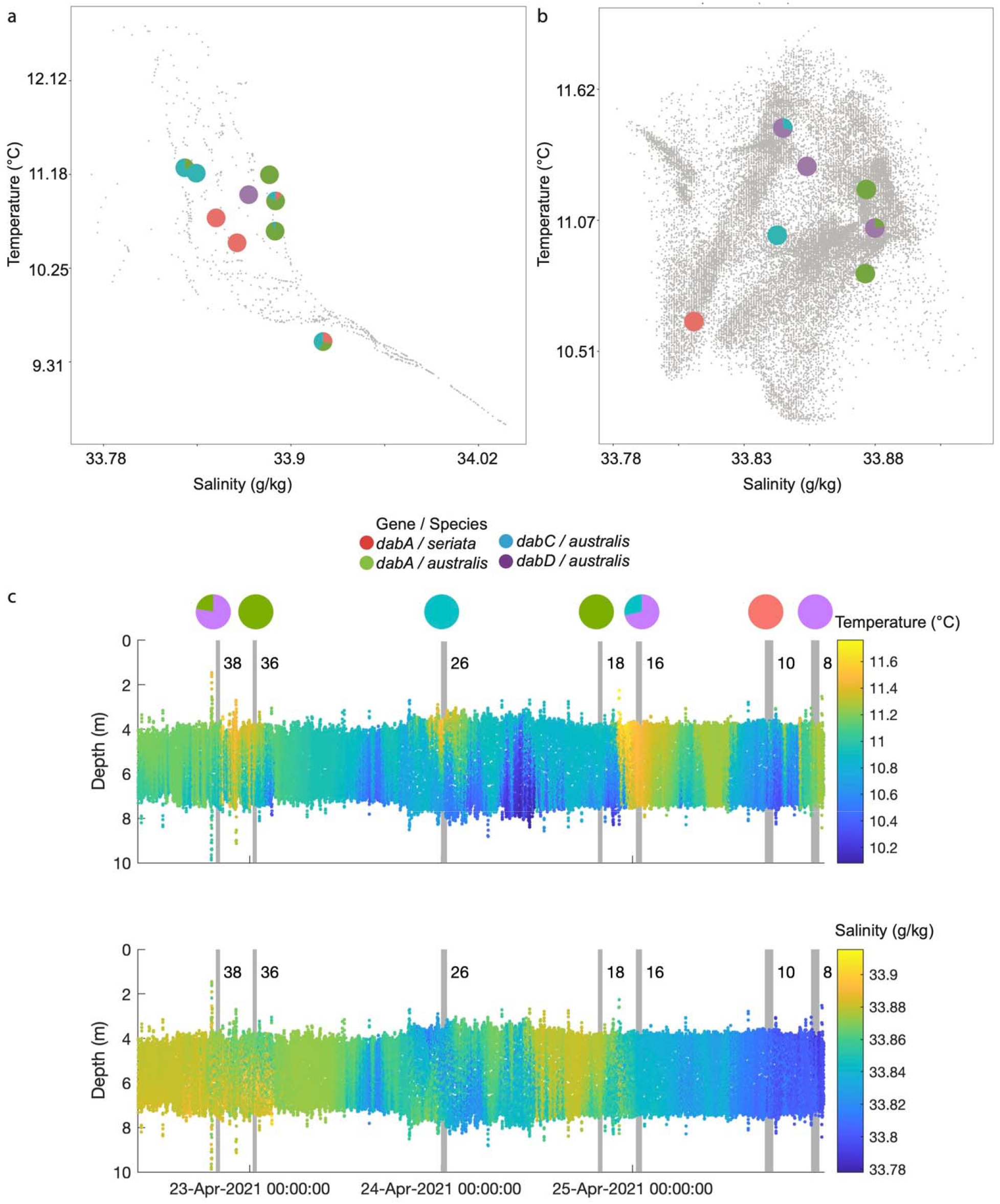
Domoic acid biosynthetic (*dab*) gene expression in expressed gene copies per liter of seawater (CPL) as measured in (a) Ship sampling by CTD at the SCM and as measured in (b) 3G-ESP/LRAUV sampling. Gray dots indicate all temperature and salinity measurements taken by the respective platform, and pie charts indicate the proportion of each gene’s expression in copies per liter of seawater. (c) Temperature (top) and salinity (bottom) measurements from the LRAUV over time indicate that *dab* gene expression occurs primarily in frontal features where there are abrupt changes in temperature and salinity. Gray vertical bars indicate samples taken by the 3G-ESP/LRAUV where *dab* gene expression is found, and the width of the gray bars represents the duration of water sampling. Considering the importance of *dabA* from *P. australis*, the three rightmost pie charts in (b) containing *dabA* from *P. australis* expression occur on the high salinity side of frontal features (Samples 38, 36, and 26).

The finding of gene expression variability due to the responses of freshly upwelled water at frontal features was enabled by specialized missions and rapid *in situ* sampling of the 3G-ESP/LRAUV. *In situ,* autonomous sampling has been valuable to other gene expression studies [50]. The key advantage of *in situ* 3G-ESP/LRAUV sampling over traditional ship sampling is reduced transport time of collection and processing of the water that could lead to changes in the cell’s gene expression profile.

The differential strength and persistence of upwelling and nutrient limitation differentiated the spring of 2021 from the spring of 2015, contributing to the stark differences of non-*Pseudo-nitzschia*-bloom and *Pseudo-nitzschia*-bloom conditions [14]. While DA at the surface of N1 was high relative to the other samples in the 2021 dataset, they were low when compared to 2015. Total DA values in 2021 (<10^1^ ng/L) were consistent with pDA values reported in early April of 2015 [14]. Particulate DA values in 2015 then rose rapidly to greater than 10^4^ ng/L by mid-April, an increase of two orders of magnitude.

Because DA is detected where *dab* expression is not found, we hypothesize that DA is an intrinsically stable metabolite and not entirely degraded by the heterotrophic bacterial community. Thus, DA could significantly contribute to the refractory pool of dissolved organic matter in the ocean. This intrinsic background signal of DA may drift in the environment asynchronously of *Pseudo-nitzschia* and *dab* gene expression. Related work has linked microbial taxa to chemotaxonomic “fingerprints” of metabolites [51]. Studies like these have improved our ability to connect taxa to metabolites that they take up, modify or produce. A targeted approach with a lower *m/z* cutoff would augment our untargeted metabolomics study. Profiling both endometabolites and exometabolites would also augment this study, as utilized by Brisson et al [52].

## Conclusions and Future Directions

We present results that tracked ecosystem change at the molecular level over the course of one week in the Eastern North Pacific Ocean. We demonstrated that high intensity, synchronous studies of Monterey Bay combining ship, 3G-ESP/LRAUV, and Wirewalker sampling improved the resolution of the taxonomy of microbial communities and their infochemicals. We observed that the microbial community did not significantly shift within a water mass as it moved offshore. We sampled during the transition from upwelling to relaxation as DA production was decreasing, in the absence of the conditions needed to spark a widespread and toxic bloom. A multi-omic analysis of a non-blooming population of *Pseudo-nitzschia* revealed its baseline metabolism associated with DA production in the marine ecosystem, a status rarely studied in HAB research. We report, for the first time, relative abundances of DA biosynthetic intermediate molecules and absolute abundances of *dab* genes in the ocean and how they correlate to *Pseudo-nitzschia* and DA.

Notably DA biosynthetic (*dab*) gene expression along the LRAUV track peaked at the high salinity side of boundary interfaces where recently upwelled water and resident bay water meet and mixed. The data suggests that exposure to recently upwelled water as inferred by higher salinity causes expression of *dabA* by *P. australis*. Strikingly, even in non-DA-outbreak conditions, the same factors induce DA production as in a highly toxic environmental bloom. There may be a natural role for DA production in a non-disturbed state: Just as understanding a wildfire’s natural role in a forest ecosystem allows us to improve wildfire management and plan for routine lower intensity, lower frequency burns, understanding DA’s natural role in the marine ecosystem may provide information for prevention and management of toxic and destructive events. Indeed, a recent study illustrates the correspondence between *Pseudo-nitzschia* blooms and productivity of pelagic and benthic habitats during low-DA conditions in Monterey Bay, highlighting the management value to identifying DA vectors during subacute DA events [53]. Complimentarily, our study serves to advance understanding of the molecular biology of *Pseudo-nitzschia* that underlies devastating toxic events.

Future directions to this work will test the hypothesis that *dab* gene expression occurs at the high salinity side of frontal features by sampling fronts during a larger *Pseudo-nitzschia* bloom in Monterey Bay during the spring of 2023. *In situ* molecular sampling of microbial communities is typically limited by discrete sampling and lack of integration of multiple ‘omic datasets, but future studies that couple fine-resolution measurements in a quasi-Lagrangian and Eulerian matter improve understanding of water mass evolution in coastal regions. Our results corroborate that most molecules in the ocean are uncharacterized compounds, emphasizing the limitation of uncultured oceanic phytoplankton. Integrated multi-omic studies like this one will improve our ability to uncover the “microbe-metabolite network” in the ocean [54]. The results of these studies provide fundamental science to drive environmental monitoring of HABs and to understand the largely understudied microbial metabolism.

## Acknowledgements

Thank you to Zachary Quinlan, Ralph Torres, Lihini Aluwihare, Katherine Barbeau, Yasin El Abiead, Jayme Smith, and Steffaney Wood for helpful discussions and guidance. Thank you to Chloe Weinstock, Kevan Yamahara, and the crew of the *R/V* Rachel Carson for help with sample collection. We thank the crew of the R/V Paragon for help with deploying the Wirewalker, LRUAV, and Waveglider and the watch-standers during LRAUV and Waveglider operations. Thank you to Brian Bill at the NOAA Northwest Fisheries Science Center for use of *Pseudo-nitzschia australis* strain NWFSC-722. Thank you to the NOAA ECOHAB Award NA19NOS4780181 for support of the project.

This study was supported by grants to AEA from the National Science Foundation (NSF-OCE-2224726, NSF*-*OCE*-*1756884, NSF IOS-2103715), the National Ocean and Atmospheric Administration (NA19NOS4780181), and the Simons Foundation (970820). MT was supported by NA19NOS4780181. ATA was supported by Moore Foundation. CMP was supported by David and Lucile Packard Foundation. ZF was supported by Simons Collaboration on Principles of Microbial Ecosystems (PriME, Grant ID 970820 to AEA). WU was supported by NA19NOS4780181 and David and Lucile Packard Foundation. AJL was supported by NA19NOS4780181. CRA was supported by NA19NOS4780181. CAS was supported by NA19NOS4780181 and David and Lucile Packard Foundation through support allocated by MBARI. JPR was supported by NA19NOS4780181 and David and Lucile Packard Foundation through support allocated by MBARI.

## Data Accession

Mass spectrometry data were deposited in a public repository (MassIVE MSV000089287). The molecular networking job for EcoHAB 2021 FBMN can be publicly accessed <u>here</u>. Data collected by the robotic fleet and their movement during the campaign are publicly available at https://stoqs.mbari.org/stoqs_canon_april2021/.

The data reported in this paper have been deposited in the NCBI sequence read archive (BioProject accession no. PRJNA1025262; BioSample accession nos. SAMN37718780-SAMN37718866; SAMN37718681-SAMN37718764; SAMN37767034-SAMN37767117; SAMN37769100-SAMN3776918.

## References

1. Azam F, Fenchel T, Field JG, Gray JS, Meyer-Reil LA, Thingstad F. The ecological role of water-column microbes in the sea. Mar Ecol Prog Ser 1983; 10: 257–263.

2. Paerl HW Justic, D. Treatise on estuarine and coastal science. Treatise on Estuarine and Coastal Science. 2011. pp 23–42.

3. Huyer A. Coastal upwelling in the California current system. Prog Oceanogr 1983; 12: 259–284.

4. Bruland KW, Rue EL, Smith GJ. Iron and macronutrients in California coastal upwelling regimes: Implications for diatom blooms. Limnol Oceanogr 2001; 46: 1661–1674.

5. Moran MA, Kujawinski EB, Schroer WF, Amin SA, Bates NR, Bertrand EM, et al. Microbial metabolites in the marine carbon cycle. Nat Microbiol 2022; 7: 508–523.

6. Bates SS, Hubbard KA, Lundholm N, Montresor M, Leaw CP. *Pseudo-nitzschia, Nitzschia*, and domoic acid: New research since 2011. Harmful Algae 2018; 79: 3–43.

7. Hampson DR, Manalo JL. The activation of glutamate receptors by kainic acid and domoic acid. Nat Toxins 1998; 6: 153–158.

8. McCabe RM, Hickey BM, Kudela RM, Lefebvre KA, Adams NG, Bill BD, et al. An unprecedented coastwide toxic algal bloom linked to anomalous ocean conditions. Geophys Res Lett 2016; 43: 10,366–10,376.

9. Teitelbaum JS, Zatorre RJ, Carpenter S, Gendron D, Evans AC, Gjedde A, et al. Neurologic sequelae of domoic acid intoxication due to the ingestion of contaminated mussels. N Engl J Med 1990; 322: 1781–1787.

10. Pulido OM. Domoic acid toxicologic pathology: a review. Mar Drugs 2008; 6: 180–219.

11. Brunson JK, McKinnie SMK, Chekan JR, McCrow JP, Miles ZD, Bertrand EM, et al. Biosynthesis of the neurotoxin domoic acid in a bloom-forming diatom. Science 2018; 361: 1356–1358.

12. Smith J, Connell P, Evans RH, Gellene AG, Howard MDA, Jones BH, et al. A decade and a half of *Pseudo-nitzschia spp.* and domoic acid along the coast of southern California. Harmful Algae 2018; 79: 87–104.

13. Lewitus AJ, Horner RA, Caron DA, Garcia-Mendoza E, Hickey BM, Hunter M, et al. Harmful algal blooms along the North American west coast region: History, trends, causes, and impacts. Harmful Algae 2012; 19: 133–159.

14. Ryan JP, Kudela RM, Birch JM, Blum M, Bowers HA, Chavez FP, et al. Causality of an extreme harmful algal bloom in Monterey Bay, California, during the 2014–2016 northeast Pacific warm anomaly. Geophys Res Lett 2017; 44: 5571–5579.

15. Bowers HA, Ryan JP, Hayashi K, Woods AL, Marin R, Smith GJ, et al. Diversity and toxicity of *Pseudo-nitzschia* species in Monterey Bay: Perspectives from targeted and adaptive sampling. Harmful Algae 2018; 78: 129–141.

16. Wells ML, Trick CG, Cochlan WP, Hughes MP, Trainer VL. Domoic acid: The synergy of iron, copper, and the toxicity of diatoms. Limnol Oceanogr 2005; 50: 1908–1917.

17. Franck V, Bruland K, Hutchins D, Brzezinski M. Iron and zinc effects on silicic acid and nitrate uptake kinetics in three high-nutrient, low-chlorophyll (HNLC) regions. Mar Ecol Prog Ser 2003; 252: 15–33.

18. Olson M, Lessard E, Wong C, Bernhardt M. Copepod feeding selectivity on microplankton, including the toxigenic diatoms *Pseudo-nitzschia spp.*, in the coastal Pacific Northwest. Mar Ecol Prog Ser 2006; 326: 207–220.

19. Olson MB, Lessard EJ. The influence of the *Pseudo-nitzschia* toxin, domoic acid, on microzooplankton grazing and growth: A field and laboratory assessment. Harmful Algae 2010; 9: 540–547.

20. Tammilehto A, Nielsen TG, Krock B, Møller EF, Lundholm N. Induction of domoic acid production in the toxic diatom *Pseudo-nitzschia seriata* by calanoid copepods. Aquat Toxicol 2015; 159: 52–61.

21. Harðardóttir S, Pančić M, Tammilehto A, Krock B, Møller EF, Nielsen TG, et al. Dangerous relations in the Arctic marine food web: Interactions between toxin producing *Pseudo-nitzschia* diatoms and *Calanus* copepodites. Mar Drugs 2015; 13: 3809–3835.

22. Selander E, Berglund EC, Engström P, Berggren F, Eklund J, Harðardóttir S, et al. Copepods drive large-scale trait-mediated effects in marine plankton. Sci Adv 2019; 5: eaat5096.

23. Rosenfeld LK, Schwing FB, Garfield N, Tracy DE. Bifurcated flow from an upwelling center: a cold water source for Monterey Bay. Cont Shelf Res 1994; 14: 931–964.

24. Graham WM, Largier JL. Upwelling shadows as nearshore retention sites: the example of northern Monterey Bay. Cont Shelf Res 1997; 17: 509–532.

25. Ryan JP, McManus MA, Kudela RM, Lara Artigas M, Bellingham JG, Chavez FP, et al. Boundary influences on HAB phytoplankton ecology in a stratification-enhanced upwelling shadow. Deep Sea Res Part II Top Stud Oceanogr 2014; 101: 63–79.

26. Kudela RM, Bickel A, Carter ML, Howard MDA, Rosenfeld L. The Monitoring of harmful algal blooms through ocean observing: The development of the California harmful algal bloom monitoring and alert program. In: Liu Y, Kerkering H, Weisberg RH (eds). Coastal Ocean Observing Systems. 2015. Academic Press, Boston, pp 58–75.

27. Anderson CR, Kudela RM, Kahru M, Chao Y, Rosenfeld LK, Bahr FL, et al. Initial skill assessment of the California Harmful Algae Risk Mapping (C-HARM) system. Harmful Algae 2016; 59: 1–18.

28. Anderson CR, Berdalet E, Kudela RM, Cusack CK, Silke J, O’Rourke E, et al. Scaling up from regional case studies to a global harmful algal bloom observing system. Front Mar Sci 2019; 6: 250.

29. Sison-Mangus MP, Jiang S, Kudela RM, Mehic S. Phytoplankton-associated bacterial community composition and succession during toxic diatom bloom and non-bloom events. Front Microbiol 2016; 7: 1433.

30. Macintyre HL, Stutes AL, Smith WL, Dorsey CP, Abraham A, Dickey RW. Environmental correlates of community composition and toxicity during a bloom of *Pseudo-nitzschia spp*. in the northern Gulf of Mexico. J Plankton Res 2011; 33: 273–295.

31. Petras D, Koester I, Da Silva R, Stephens BM, Haas AF, Nelson CE, et al. High-resolution liquid chromatography tandem mass spectrometry enables large scale molecular characterization of dissolved organic matter. Front Mar Sci 2017; 4: 405.

32. Greenfield DI, Marin III R, Doucette GJ, Mikulski C, Jones K, Jensen S, et al. Field applications of the second-generation Environmental Sample Processor (ESP) for remote detection of harmful algae: 2006-2007. Limnol Oceanogr Methods 2008; 6: 667–679.

33. Harvey JBJ, Ryan JP, Zhang Y. Influences of extreme upwelling on a coastal retention zone. Front Mar Sci 2021; 8: 648994.

34. Kolody BC, McCrow JP, Allen LZ, Aylward FO, Fontanez KM, Moustafa A, et al. Diel transcriptional response of a California Current plankton microbiome to light, low iron, and enduring viral infection. ISME J 2019; 13: 2817–2833.

35. Anderson CR, Brzezinski MA, Washburn L, Kudela R. Circulation and environmental conditions during a toxigenic *Pseudo-nitzschia australis* bloom in the Santa Barbara Channel, California. Mar Ecol Prog Ser 2006; 327: 119–133.

36. King AL, Barbeau KA. Dissolved iron and macronutrient distributions in the southern California Current System. J Geophys Res Oceans 2011; 116: C03018.

37. Hogle SL, Dupont CL, Hopkinson BM, King AL, Buck KN, Roe KL, et al. Pervasive iron limitation at subsurface chlorophyll maxima of the California Current. Proc Natl Acad Sci USA 2018; 115: 13300–13305.

38. Paduan JD, Rosenfeld LK. Remotely sensed surface currents in Monterey Bay from shore-based HF radar (Coastal Ocean Dynamics Application Radar). J Geophys Res Oceans 1996; 101: 20669–20686.

39. Omand MM, Cetinić I, Lucas AJ. Using bio-optics to reveal phytoplankton physiology from a Wirewalker autonomous platform. Oceanography 2017; 30: 128–131.

40. Wegley Kelly L, Nelson CE, Petras D, Koester I, Quinlan ZA, Arts MGI, et al. Distinguishing the molecular diversity, nutrient content, and energetic potential of exometabolomes produced by macroalgae and reef-building corals. Proc Natl Acad Sci USA 2022; 119.

41. Dührkop K, Nothias L-F, Fleischauer M, Reher R, Ludwig M, Hoffmann MA, et al. Systematic classification of unknown metabolites using high-resolution fragmentation mass spectra. Nat Biotechnol 2021; 39: 462–471.

42. Dührkop K, Fleischauer M, Ludwig M, Aksenov AA, Melnik AV, Meusel M, et al. SIRIUS 4: a rapid tool for turning tandem mass spectra into metabolite structure information. Nat Methods 2019; 16: 299–302.

43. Lelong A, Hégaret H, Soudant P, Bates SS. *Pseudo-nitzschia* (Bacillariophyceae) species, domoic acid and amnesic shellfish poisoning: revisiting previous paradigms. Phycologia 2012; 51: 168–216.

44. Smith J, Gellene AG, Hubbard KA, Bowers HA, Kudela RM, Hayashi K, et al. *Pseudo-nitzschia* species composition varies concurrently with domoic acid concentrations during two different bloom events in the Southern California Bight. J Plankton Res 2018; 40: 29– 45.

45. Marchesiello P, McWilliams JC, Shchepetkin A. Equilibrium Structure and Dynamics of the California Current System. J Phys Oceanogr 2003; 33: 753–783.

46. Schwing FB, Gaxiola-Castro G, Goméz-Valdéz J, Kosro PM, Mantyla AW, Smith RL, et al. The state of the California Current, 2001-2002: Will the California Current System keep its cool, or is El Niño looming? CalCOFI Rep 2002; 43.

47. Samo TJ, Pedler BE, Ball GI, Pasulka AL, Taylor AG, Aluwihare LI, et al. Microbial distribution and activity across a water mass frontal zone in the California Current Ecosystem. J Plankton Res 2012; 34: 802–814.

48. Stukel MR, Aluwihare LI, Barbeau KA, Chekalyuk AM, Goericke R, Miller AJ, et al. Mesoscale ocean fronts enhance carbon export due to gravitational sinking and subduction. Proc Natl Acad Sci USA 2017; 114: 1252–1257.

49. Smayda TJ. Turbulence, watermass stratification and harmful algal blooms: an alternative view and frontal zones as “pelagic seed banks”. Harmful Algae 2002; 1: 95–112.

50. Nowinski B, Feng X, Preston CM, Birch JM, Luo H, Whitman WB, et al. Ecological divergence of syntopic marine bacterial species is shaped by gene content and expression. ISME J 2023; 17: 813–822.

51. Durham BP, Boysen AK, Heal KR, Carlson LT, Boccamazzo R, Deodato CR, et al. Chemotaxonomic patterns in intracellular metabolites of marine microbial plankton. Front Mar Sci 2022; 9: 864796.

52. Brisson V, Swink C, Kimbrel J, Mayali X, Samo T, Kosina SM, et al. Dynamic *Phaeodactylum tricornutum* exometabolites shape surrounding bacterial communities. New Phytol 2023; 239: 1420–1433.

53. Bernstein S, Ruiz-Cooley RI, Kudela R, Anderson CR, Dunkin R, Field JC. Stable isotope analysis reveals differences in domoic acid accumulation and feeding strategies of key vectors in a California hotspot for outbreaks. Harmful Algae 2021; 110: 102117.

54. Moran MA, Kujawinski EB, Schroer WF, Amin SA, Bates NR, Bertrand EM, et al. Microbial metabolites in the marine carbon cycle. Nat Microbiol 2022; 7: 508–523.

