## Supplementary material for "High Resolution Biological and Physical Sampling Reveals Expression of Domoic Acid Biosynthetic Genes at Frontal Zones": EH21_Supplementary.docx

### **Materials and Methods**

#### **Accession of Environmental Data and Rationale**

Biologically Effective Upwelling Transport Index (BEUTI) values were accessed from NOAA at latitude = 36° N (https://oceanview.pfeg.noaa.gov/erddap/). The BEUTI index was utilized because it incorporates all relevant conditions and processes to estimate nutrient supply including nutricline depth, upwelling due to divergence at the coast and wind stress curl. BEUTI is Coastal Upwelling Transport Index (CUTI) times the concentration of nitrate at the base of the mixed layer. CUTI is the sum of (1) vertical transport due to divergence/convergence in near-surface Ekman transport that is driven by alongshore winds and wind stress curl and (2) vertical transport associated with cross-shore geostrophic transport associated with an alongshore pressure gradient.

Wind speed and direction data was accessed from NOAA National Data Buoy Center at Station 46042 36.785 N 122.396 W (<https://www.ndbc.noaa.gov/station_history.php?station=46042>).

Temperature and Salinity data was accessed from Monterey Bay Aquarium Research Institution (MBARI) CTD measurements at “Station M1” Mooring (36.75ºN, 122.03ºW) through the OPeNDAP Server (<https://dods.mbari.org/opendap/hyrax/data/OASISdata/netcdf/hourlyM1.nc.html>). Station M1 is near the mouth of Monterey Bay in water about 1,000 meters deep (**Figure 1b**). Near-continuous data has been collected at Station M1 since the early 1990s.

#### **Sampling Strategy and Sample Collection:**

Ship sampling was conducted aboard the *R/V* Rachel Carson over three days: April 20, 21, and 22 of 2021 in Monterey Bay using a conductivity, temperature, and depth (CTD) instrument and Niskin bottle rosette. Ship sampling provided data from two methods of observation: Eulerian and quasi-Lagrangian. The Eulerian time series was at three locations where moored Wirewalkers were placed, and these three locations were sampled once daily by the ship [1]. These moorings were in the domoic acid source region based on pre-experimental reconnaissance of *Pseudo-nitzschia* species and domoic acid measurements.

The quasi-Lagrangian time series was collected by the Ship drifting with a robotic fleet of a 3G-ESP/LRAUV and a Waveglider. The methods of tracking the subsurface 3GESP/LRAUV with a Waveglider, demonstrated for open ocean studies, are described in . Four daily stations for sample collection were placed at a fixed distance around the 3G-ESP/LRAUV drifting vehicle in a 2-km grid: North, South, East, and West of the instrument. Ship sampling at both Eulerian and Lagrangian stations took place at three depths: Shallow (approximately 5 meters), Subsurface Chlorophyll Maximum (SCM, variable), and Deep (variable). Eulerian moored stations are referred to as N1, E1, and S1, and stations N2, S2, E2, W2 refer to the Lagrangian stations. On April 21 and 22, four 2-km Lagrangian stations were sampled, while on April 20 only three of the four stations were sampled. The 2-km grid around the ESP is not perfectly symmetric on a map because the vehicle drifted between sampling each station, and the positions were defined relative to the position of the vehicle at the time of sampling.

The Lagrangian approach was employed to focus on temporal variation and remove spatial variation by moving with the moving phytoplankton population. This approach was initially challenged due to the tracked phytoplankton patch shoaling and causing the 3G-ESP/LRAUV to surface. The LRAUV mission was then adapted to remain at a fixed depth within the mixed layer / upper thermocline. Because during this period, neither strong upwelling nor strong environmental gradients were present, the Lagrangian grid sampled at 2 km points around the drifting ESP, appropriate to the weaker gradients present. This sampling strategy was designed to provide a richer data set given the environmental conditions and available observing assets.

##### **3G ESP Filter Collection and Sample Preservation:**

The position, temperature, salinity, chlorophyll relative fluorescence, light intensity, and backscatter were measured every two seconds by the 3G-ESP/LRAUV instrument. 3G-ESP/LRAUV’s filter samples were collected during the three-day ship-sampling and for three days thereafter to create a 6-day dataset.

The ESP/LRAUV collected and preserved samples as previously described [2–4]. Here, 25 mm 0.2 µm hydrophilic PVDF filters were loaded into cartridges. When the LRAUV directed the ESP to sample, the ESP’s internal sample loop was flushed with 110 mL of seawater. Seawater was directed through a filter contained in a cartridge and filtration continued until a 1 L volume was reached or the filter clogged (filtration rate < 0.2mL/second). At the end of filtration, the volume was recorded and the sample preserved using RNAlater as previously described [2, 3]. Samples collected on the ESP/LRAUV remained on board until after field operations, when samples were transferred to 2ml screw top tubes and stored at -80°C until processed.

##### **Sterivex Sample Collection For DNA and RNA:**

Two peristaltic pump motors were set up, each equipped with three single channel pump heads and six lines of tubing: One pump was used for DNA and the other for RNA. Whole ocean water samples for DNA and RNA were collected by filtration of a recorded volume of about 2 L and 4 L respectively onto separate 0.22 µm Sterivex-GP filters as follows (MilliporeSigma, Burlington, MA, USA): 4 liters of water was collected from CTD rosettes in amber opaque bottles for RNA filtering, and 2 liters of whole sea water was collected in clear bottles from CTD rosettes for DNA filtering. Bottles were rinsed with sea water three times prior to collecting samples. Bottles were filled slightly over the 2-liter or 4-liter mark. Lines were primed with excess seawater. Then, pumps were paused, labeled 0.22 µm Sterivex-GP filters (MilliporeSigma, Burlington, MA, USA) were attached, and the pump was started at 150 rpm, collecting biomass onto the Sterivex filter. Start time was noted, followed by the end time either when the volume was depleted, the filter was clogged, or thirty minutes had passed. The pump was paused, the line was released, and the filter was removed. A 50 mL syringe was used to purge extra water out of the filter three times. The ends of the Sterivex were plugged with a sterile luer-lock plug and hematocrit sealant, wrapped in aluminum foil, and were flash frozen in liquid nitrogen and stored in the cryo-can until transport to shore and storage in a -80 °C freezer. Lines were flush lines with 1 liter of Milli-Q water between samples.

##### **Chlorophyll Sample Collection and Processing:**

A vacuum pump and manifold containing three 47 mm filter holders with 500 mL collection reservoirs were set up. A 47 mm GF/F filter was placed on each filter holder. Whole sea water was collected from the CTD rosette in a 1 L brown opaque bottle. The bottle was rinsed with sea water three times prior to collecting samples. 500 mL of seawater was poured into the vacuum manifold collection reservoir. The vacuum pump was started, and the valves were opened. When filtration was completed, valves were closed, and the pump turned off. Flat forceps were used to remove filters, fold the filter in half, and place the filter into a 5 mL cryovial. The cryovial was flash frozen in liquid nitrogen and stored in a cryo-can until transport to shore and storage in a -80 °C freezer. Reservoirs were rinsed with filtered natural sea water between samples.

The protocol for extraction and quantification of chlorophyll is available here: <https://calcofi.org/sampling-info/methods/> [5–7]. In short, filters were placed in a 10 mL screw top culture tube containing 8.0 mL of 90% HPLC-grade aqueous acetone and placed at -20°C for 24 hours. After 24 hours, samples were brought to room temperature for one hour and the fluorescence was read on a Turner 10 AU fluorometer to calculate the concentration of chlorophyll-a and phaeopigments. Samples were acidified with 100 µL of 10% by volume of 37% hydrochloric acid to degrade chlorophyll to phaeopigments and the fluorescence was read again on a fluorometer to calculate the concentration of phaeopigments and chlorophyll-b. A chlorophyll standard was used to calibrate the fluorometer and a blank of 90% acetone was used (Sigma Aldrich C6144).

##### **Flow Cytometry Sample Collection and Processing:**

1 mL whole water was transferred to 1.8 mL cryovials, and 100 µL glycerol in pH 8.0 TE buffer (10mM Tris-HCl containing 1mM EDTA Na_2_) was added to each cryovial for preservation. Each vial was mixed gently by inverting and incubated for 1 minute at ambient temperature. The cryovial was flash frozen in liquid nitrogen and stored in a cryo-can until transport to shore and storage in a -80 °C freezer. Samples were not processed or analyzed for this study but are available upon request.

##### **Nutrient Analysis:**

Nutrient samples were collected and processed following methods used in other studies in Monterey Bay [8]. Nutrient samples were collected alongside all CTD samples, were frozen on the ship, and were processed downstream on an AlpChem autoanalyzer to quantify nitrate, nitrite, phosphate, and silicate concentrations [9].

#### **DNA Extraction and Preparation of Amplicon Libraries**

**DNA Extraction with Internal Standard DNA addition**

Sterivex filters were extracted following published methods [10, 11].

Single-use aliquots of internal standard gDNA of organisms not expected in a marine dataset were added to the extraction lysis buffer mix just prior to extraction [12]. *Schizosaccharomyces pombe (S. pombe)* gDNA (ATCC [Manassas, VA, USA] 24843D-5) at 2.12 ng per sample (1 µL of 2.12 ng/µL gDNA) was added as the 18S rRNA gene internal standard, and *Thermus thermophilus (T. thermophilus)* gDNA (ATCC 27634D-5) at 4.09 ng per sample (1 µL of 4.09 ng/µL) was added as the 16S rRNA gene internal standard. gDNA stock solutions were quantified using a Qubit fluorometer prior to addition (Thermo Fisher Scientific, Waltham, MA, USA).

DNA was extracted and purified with the NucleoMag Plant Kit (Macherey-Nagel, Düren, Germany) on an epMotion 5057TMX (Eppendorf, Hamburg, Germany) as described on protocols.io: <https://doi>.org/10.17504/protocols.io.bc2hiyb6. DNA extraction size and quality was verified on a 1.8% agarose gel after extraction.

**Amplicon Library Construction and Sequencing**

Amplicon libraries were generated as described doi: dx.doi.org/10.17504/protocols.io.bmuck6sw [10, 13]. 16S rRNA genes were amplified by PCR with V4-V5 primer set 515 F (5’-GTGYCAGCMGCCGCGGTAA-3’) and 926 R (5’-CCGYCAATTYMTTTRAGTTT-3’) and 18S rRNA genes were amplified with PCR using V9 primer set 1389 F (5’-TTGTACACACCGCCC-3’) and 1510 R (5’-CCTTCYGCAGGTTCACCTAC-3’) [14–16]. The ITS2 region was amplified from both the ship and the AUV samples using an unpublished primer (5.8SF- 5’-TGCTTGTCTGAGTGTCTGTGGA-3’; 28SR- 5’-TATGCTTAAATTCAGCGGGT-3’ that were designed to specifically target *Pseudo-nitzschia* ITS2 [17].

Traditional methods of identification of *Pseudo-nitzschia* species have involved robust morphological identification by light microscopy of structural features [18–20]. Despite its accuracy, morphological identification is time and labor-intensive, requires specialized instrumentation, cannot reliably identify cryptic species and subspecies, and is not feasible for environmental sampling with low concentrations of cells. Therefore, molecular phylogeny is often utilized for *Pseudo-nitzschia* species delineation. Due to the low 18S sequence variability among *Pseudo-nitzschia* species, 18S amplicons are not accurate in delineating species specificity [21]. Other methods to identify *Pseudo-nitzschia* species include quantitative polymerase chain reaction (qPCR), large-subunit ribosomal RNA (LSU rRNA)-targeted oligonucleotide probes, sandwich hybridization probes, automated ribosomal intergenic spacer analysis (ARISA), and molecular barcoding of internal transcribed spacer 1, mitochondrial *cox1,* and the nuclear-encoded large subunit (LSU) [21–27]. The phylogeny of *Pseudo-nitzschia* has also been resolved using internal transcribed spacer 2 (ITS2) and matches morphological species delineations [27].

Primer pairs contain unique combinations of two barcorded indices, allowing downstream demultiplexing. DNA was amplified with a one-step PCR using the TruFi DNA Polymerase PCR kit (Azura Genomics, Raynham, MA, USA). Each reaction was performed with an initial denaturing step at 95°C for 1 min followed by 30 cycles of 95°C for 15 sec, 56°C for 15 sec, and 72°C for 30 sec for the 16S, 18SV9, and ITS2 amplicons.

Amplification was confirmed by running 2.5 μL of each PCR reaction on a 1.8% agarose gel. PCR amplicon products were purified using AMPure XP beads (Beckman Coulter) following the standard 1.5x PCR clean-up protocol. PCR quantification was performed in duplicate using Quant-iT PicoGreen dsDNA Assay kit (Invitrogen). Samples were then pooled in equal proportions followed by another 0.8x AMPure XP bead purification. Pools were evaluated on an Agilent 2200 TapeStation and quantified with Qubit HS dsDNA (Thermo Fisher Scientific). Each pool was sequenced at the University of California, Davis Sequencing Core on a single Illumina MiSeq lane (2×300bp for 16S, 18SV4, and ITS2, 2×150bp for 18SV9) with a 15% PhiX spike-in.

#### **Bioinformatic Assembly, Taxonomic Classification and Visualization of Amplicon Libraries**

Six amplicon datasets were analyzed: 16S, 18SV9, and ITS2 from both the ship and the AUV sampling, referred to here on as “16S-AUV”, “18SV9-AUV”, “ITS2-AUV”, “16S-Ship”, “18SV9-Ship”, and “ITS2-Ship.”

Amplicons were analyzed with Qiime2 (version qiime2-2023.2) [20–24][28]. Demultiplexed paired-end reads were trimmed to remove adapter and primer sequences with cutadapt, and denoised with DADA2 to yield amplicon sequence variants (ASVs) [28–30]. Taxonomic annotation was performed with SILVA and PR2 for 16S and 18S amplicons respectively, and with a classifier built and trained on a custom database for ITS2 [27, 31–34]. Sequences classified as chloroplasts within the 16S datasets were removed and analyzed as a separate plastid dataset using a qiime2 naïve-bayes classifier trained on the PhytoREF database for annotation [35].

Analysis was carried out using R, particularly the PhyloSeq version 1.34.0 [36].

**Statistical analysis**

Statistical analysis was performed using R. To determine the statistical significance of PcoA plots (weighted unifrac), the phyloseq package in R was used to calculate distances and then the vegan package adonis 2 function in R was used to run PERMANOVA [36, 37].

To determine the statistical significance of boxplots, a histogram of the data was first plotted. To test for normality, a Shapiro-Wilk test was used. If the histogram was not normal, statistical tests were performed that do not assume normality. For categorical data, a Kruskal-Wallis test (the non-parametric equivalent of an ANOVA) was used. For data with only two levels, a Wilcoxon rank sum test (non-parametric equivalent of t-test) was run. For example, a Kruskal-Wallis test was used for box plots comparing alpha diversity between depths. Pairwise tests were performed between depth classes (i.e., deep vs. shallow, shallow vs. SCM, SCM vs. deep) using the Wilcoxon Rank Sum Test.

**Internal standards** were utilized in DNA extractions to aid in quantitative microbial community profiling following published methods [12]. To calculate the total number of 16S or 18S rRNA gene copies spiked into each sample, it was assumed that the average weight of a base pair is 650 Daltons (650 g/mol), the 16S rRNA gene copy number per cell for *T. thermophilus* gDNA was 2, the 18S rRNA gene copy number per cell for *S. pombe* was 110, the genome size of *T. thermophilus* was 2.13 Mb, and that the genome size of *S. Pombe* was 13.8 Mb. The total number of internal standard rRNA gene copies spiked into each sample was calculated as 1.57*10^7 18S *S. pombe* gene copies per sample, and 3.56*10^6 16S *T. thermophilus* gene copies per sample. Subsequently, abundance of each out a gene copies per mL was calculated using the number of internal standard rRNA gene copies spiked into each sample, the number of gene standard reads sequenced in each sample, the number of reads of toutOTU sequenced, and the volume of seawater filtered in each sample (2 L).

**16S-AUV amplicons** were rarefied to 47022 OTUs and all samples were retained. 5490 OTUs no longer remained in the dataset.

Jupyter Notebook “EH21_AUV_16S.ipynb” file is provided for data processing and DADA2 parameters.

**18SV9-AUV amplicons** were rarefied to 24470 OTUs and all but one sample was retained. Sample EH28_A114 (Collected by the AUV on 4/23/21 at 18:55 at position -122.0182338, 36.81509781) was not included in the AUV 18SV9 amplicon analysis due to poor sequencing depth. 334 OTUs no longer remained in the dataset.

Jupyter Notebook “EH21_AUV_18SV9.ipynb” file is provided for data processing and DADA2 parameters.

**ITS2-AUV amplicons** were rarefied to 17321 OTUs and all samples were retained. 128 OTUs no longer remained in the dataset.

Jupyter Notebook “230504_EH21_ITS2_AUV_dada2.ipynb” file is provided for data processing and DADA2 parameters.

**16S-Ship amplicons** were rarefied to 33601 reads, maintaining all samples: 4300 OTUs no longer remained in the dataset. 16S quantitative libraries were rarefied to 322795 gene copies per mL seawater, maintaining all 60 samples and 3410/3710 OTUs: 300 OTUs no longer remained in the dataset.

Jupyter Notebook “EcoHAB2021_16S_dada2.ipynb” file is provided for data processing and DADA2 parameters.

**18SV9-Ship amplicons** were rarefied to 46750 reads, maintaining 57/60 of our samples and 3054/3304 OTUs: 3 samples were removed due to small library size and 250 OTUs no longer remained in the dataset. 18SV9 quantitative libraries were rarefied to 136150 gene copies per mL seawater, maintaining 57/60 samples and 3075/3079 OTUs: 3 samples were removed due to small library size and 4 OTUs no longer remained in the dataset.

The three samples that were removed due to small library size were resequenced and analyzed with Qiime2 as described above. Dada2-tables and Dada2-rep-seqs tables were merged from the original dataset with the three re-sequenced samples, and rarefied to 46756 reads, such that all samples had higher quality sequencing data. With rarefication, the three samples with low sequencing depth were removed, as were 840 OTUS, retaining 3803 taxa and 60 samples. The merged data tables were analyzed with the Qiime workflow, statistics, and in R with Phyloseq.

Jupyter Notebook “EcoHAB2021_18SV9_dada2.ipynb”, and “EH21_Ship_18SV9_3resequenced.ipynb” files are provided for data processing and DADA2 parameters.

**ITS2-Ship amplicons** were rarefied to 10 reads, maintaining all samples: 678 OTUs no longer remained in the dataset.

Jupyter Notebook “230504_EH21_ITS2_Ship_dada2.ipynb” file is provided for data processing and DADA2 parameters.

#### **Metatranscriptomes:**

**RNA Extraction and cDNA generation from 3G-ESP Samples**

RNA was extracted with the NucleoMag RNA kit (Macherey-Nagel, Düren, Germany) on an epMotion 5072 TMX (Eppendorf, Hamburg, Germany) [11].

RNA spikes #2 and #7 from the Invitrogen ArrayControl RNA Spikes kit (Catalog# Invitrogen AM1780) were added to the lysis buffer during extraction [38].

cDNA was generated from total RNA using random hexamer primers using the AzuraFlex cDNA synthesis kit (Azura Genomics, Raynham, Massachusetts). This cDNA was utilized for amplicon library preparation for the ESP samples.

**Preparation and sequencing of polyA-enriched RNA**

polyA-enriched RNA was prepared from total RNA using oligo dT primers using the NEBNext® Poly(A) mRNA Magnetic Isolation Module (for 96 samples) kit (New England Biolabs, Ipswich, MA, USA; Catalog E7490L) and the NEBNext® Ultra™ II Directional RNA Library Prep with Sample Purification Beads kit (Catalog# E7765L).

**Assembly and annotation of RNA sequencing libraries**

Raw reads were adapter- and quality-trimmed using fastp v0.23.2 (trimmomatic option [39, 40]. rRNA-matched trimmed were further filtered out using BBduk ([https://jgi.doe.gov/data-and-tools/software-tools/bbtools/;](https://jgi.doe.gov/data-and-tools/software-tools/bbtools/) rDNA databases PR2 v4.12.0, RFAM v14.1, SILVA v138). Metatranscriptomes were assembled by-sample and by-group using MEGAHIT v1.2.9, and the longest contigs were retained via CD-HIT-est v4.8.1 clustering (0.95 sequence identity) [41, 42]. Open reading frames were determined and translated by FragGeneScan followed by annotation against common protein domain databases Pfam v35.0, PANTHER v15.0, TIGRFAM v15.0, KEGG v30-01-2023, and EggNOG v5 using a combination of tools (diamond v2.0.15, eggNOG-mapper v2.1.10, Interproscan v5.57-90.0, kofamscan v1.3.0 [43–48]. Lineage Probability Index (LPI) was calculated from top 100 diamond blastp hits by dividing the sum of probabilities of each taxonomic term by the normalization factor corresponding to its taxonomic level in the lineage and choosing the term with the highest index [49].

**Quantification of transcripts (Spike IDs, TPM, CPL)**

Following cell lysis, 5×10e9 molecules of each RNA spike #2 and #7 were added to samples, allowing for internal standard normalization. ORF read map counts for the metatranscriptomic assembly were obtained by Bowtie2 v2.5.0 (--local mode, otherwise default setting keeping best read alignments only) [50]. Before downstream processing, transcript counts per liter were calculated [51]. Briefly, ORFs corresponding to RNA spikes #2 and #7 were identified in the assembly by BLASTN v2.13.0+ (-evalue 1e-3, hits with bitscore < 50 discarded), their raw counts summed and the respective rows removed from the metatranscriptomic matrix. Transcript per million (TPM) values were calculated, first normalizing to ORF size in kbp (RNA spike size for spike counts) and then to the reads per kilobase total. Counts per liter were counted using the spike coefficient (sum of spike molecules added 1×10e10 / sum of spike TPM) and divided by sample volume in liters.

Relative expression of ORFs within a taxonomic group was calculated as a fraction of their CPL values in the CPL-total of all ORFs of that group as determined by their LPI annotation.

**Identification of *dab* genes and Alignments**

Full-length and partial ORFs encoding DAB biosynthesis enzymes were identified by BLAST searches in both DNA and protein space with published reference sequences as queries [52, 53]. Protein sequences were aligned by MUSCLE v5.1, visually checked, and filtered from paralogs based on phylogeny (IQ-TREE v2.0.3 with LG4X substitution matrix and ultra-fast bootstrapping) [54, 55] (**Figure S5a).**

**Identification of conserved expression modules (WCGNA)**

The Weighted Gene Correlation Network Analysis (WGCNA) R package was used to identify modules of ORFs in the metatranscriptomes with similar expression patterns [56, 57]. For this analysis, only metatranscriptomics from ORFs from Ship Surface, Ship SCM, or AUV samples were considered: Ship Deep and all blanks and positive/negative controls were excluded from analysis. Samples were normalized by copies per liter (CPL) prior to analysis. To retain statistically relevant data, samples were then filtered by only considering those with at least 50 raw counts in at least 40% of samples, leaving 15574 OTUs for WGCNA analysis. A Pearson correlation matrix was constructed with this dataset, and an adjacency matrix was constructed from that by applying a power function (AF(s)=s^b), where “b” represents the soft-thresholding parameter. B=16 was found to be the lowest b value that allowed for a scale-free topology R-squared value above 0.8, and was chosen as recommended in the WGCNA user manual, in order to optimize the mean number of connections of the network while preserving scale independence. A signed Topological Overlap Matrix (TOM) was constructed from the correlation matrix to measure dissimilarity between each pair of nodes based on shared neighbors. A dendrogram of genes was created using average linkage hierarchical clustering using the “hclust” function, and the minimum module size was defined as 30 ORFs. Modules of co-expression were identified using the function cuttreeDynamic and were labeled with colors. Subsequently, modules with similar expression profiles were merged by using the “moduleEigengenes” function with default parameters to calculate “eigengenes” (a measure of “average” expression calculated as the first principal component of the ’odule's expression matrix) for each module. Modules with correlated eigengenes were merged by setting a cut height threshold on the dendrogram to 0.5, yielding seven modules.

Driver genes for each module were identified by calculating the student asymptotic p-value for the given correlations between modules and genes, and examining those with the lowest p-values, indicating the highest significance. The top fifty genes highly associated with each module are provided in *Supplementary Information.*

“MEGrey” module contains remaining contigs that did not fit into a module of co-expression contigs with low variance explained by the model (n=311). “MEhoneydew1” contained 12328 ORfs, “MEindianred4” contained 1387 ORfs, “MEblack” contained 753 ORfs, “MEdarkolivegreen4” contained 356 ORFs, and module “MEmediumpurple2” contains 321 ORFs. Module “MEpalevioletred3” contains 77 ORFs that are mostly unannotated.

#### **Metabolites**

Whole sea water was concentrated by solid phase extraction and was queried with liquid chromatography tandem mass spectrometry. Features were analyzed and annotated with Global Natural Products Social Network (GNPS) [58]. SIRIUS and CANOPUS were used to predict molecular formulas based on isotope patterns in MS spectra and MS/MS fragmentation patterns and compound classes based on MS/MS fragmentation pattern libraries, respectively [59, 60]. Toxin data were derived from both an Environmental Sample Processor (ESP) with a Surface Plasmon Resonance (SPR) instrument on an autonomous underwater vehicle (AUV) that followed the high-biomass region, and from ship-based whole water and filter sampling for LC-MS/MS metabolomics analysis.

**Particulate Domoic Acid Sample Collection and Processing:**

A vacuum pump and manifold 3 × 25 mm filter holders with 200 mL collection reservoirs were set up. A 25 mm GF/F filter was placed on each filter holder. Whole sea water was collected from the CTD rosette in a 1 L brown opaque bottle. The bottle was rinsed with sea water three times before collecting samples. 200 mL of seawater was poured into the vacuum manifold collection reservoir. The vacuum pump was started, and the valves were opened. When filtration was completed, valves were closed, and the pump turned off. Flat forceps were used to remove filters, fold the filter in half, and place the filter into a 2 mL cryovial. Cryovial was flash frozen in liquid nitrogen and stored in the cryo-can until transport to shore and storage in a -80 °C freezer. Reservoirs were rinsed with filtered natural sea water between samples.

Samples were processed by thawing filters on ice and extracting with 1200 µL of 50% aqueous methanol (LC-MS grade) in the cryovial. Samples were sonicated for 5 minutes in a sonication bath and vortexed for 10 minutes. The filter was removed, and the extract was transferred to a 0.2 μm filter tube and centrifuge for 5 minutes at 13,000× g. The flowthrough was dried in a speedvac. The dried extract was resuspended in 400 µL of a solution of 50% LC-MS grade methanol, 49% LC-MS grade water, and 1% LC-MS grade formic acid. Sample was transferred to an LC-MS vial for analysis.

**Particulate Domoic Acid Data Analysis:**

High resolution liquid chromatography mass spectrometry (HRMS) measurements were carried out on an Agilent Technologies 1200 Series system with a diode-array detector coupled to an Agilent Technologies 6530 accurate-mass Q-TOF LCMS run in positive ionization mode. Compounds were separated by reversed-phase HPLC chromatography on a Phenomenex Kinetex 5 mm C18 100 Å 150 × 4.6 mm LC column with water + 0.1% formic acid (solvent A) and acetonitrile + 0.1% formic acid (solvent B) as eluents. The following gradient was applied at a flow rate of 0.5 mL/min: hold at 5% B for 0.5 minutes, 5% to 62% B over 9.5 minutes, 62% to 100% B over 6 minutes, and 100% to 5% B for 6 minutes. To adequately desalt samples and prevent damage to the LCMS, the valve was diverted to waste for 0-5 minutes and 10-22 minutes. The MS/MS was targeted at 312.14 m/z and 314.15 m/z. Samples were run alongside a standard of domoic acid.

Quantifications were conducted in GNPS Dashboard by integrating the peak area of extracted ion chromatograms [61].

**Total Metabolomics Sample Collection and Processing:**

Whole sea water was concentrated with solid phase extraction following published methods [62–64]. This method has been used in environmental studies [65, 66]. One liter of whole sea water from the CTD rosette was collected into a methanol-and-Milli-Q-washed 1 liter bottle and stored on wet ice for transport to the laboratory. Samples were acidified with 1 mL of 37% trace metal grade hydrochloric acid to a pH of 2.0. Along with each daily sample set, 1 liter of LC-MS water was transferred into a clean 1-liter bottle and acidified with 1 mL 37% trace metal grade hydrochloric acid to a pH of 2.0 as a procedural blank.

For solid phase extraction, materials were prepared by soaking adapters in 3.7% hydrochloric acid for 24 hours and rinsing in Milli-Q water. The solid phase extraction station was set up following described methods on protocols.io. 200 mg PPL cartridges (BondElut, Agilent) were washed to activate the hydrophobic beads with 9 mL methanol (3× column volume), 9 mL pH 2 water (prepared as 1 mL 37% trace metal grade hydrochloric acid into 1 L of LC-MS grade water), 9 mL methanol, 9 mL pH 2 water. Samples were pulled into the column reservoir slowly at a rate of about 1 drop every 2 seconds. Once the sample volume went through the column or the column was clogged and the volume filtered was noted, the PPL cartridge was flushed with 2 column volumes (~6 mL) pH 2 water to remove salt from the cartridge. The sample was frozen at -80 °C and thawed before drying under Nitrogen gas until the color of the cartridge changed to a light yellow. Cartridges were individually wrapped in foil and frozen at -80 °C until elution. Samples were eluted from the PPL cartridge into an LC-MS vial with 2 mL LC-MS grade methanol with gravity followed by a syringe to push the eluent into the capturing LC vial. LC vials were capped and dried in a Centrivap overnight. Dried sample extracts were resuspended in 100 uL of MeOH/H_2_O/Formic Acid (79/20/1), collected with a pipette tip and dispensed back into the vial with an insert (200-250 uL).

**Total Metabolomes Sample Processing and Data Analysis:**

**UHPLC-MS/MS Analysis**

UHPLC-MS/MS analysis follows protocols developed [62, 65]. A Vanquish reverse-phase UHPLC system was used for liquid chromatography. The column used for chromatography had a stationary phase of C18 porous core shell column (Kinetex C18, 150 × 2 mm, 1.8 μm particle size, 100 Å pore size, Phenomenex). The column was conditioned before use with a 99% acetonitrile/1% water to 5% acetonitrile/95% water gradient over 10 incremental 5-minute steps. Solvents used as the mobile phase were H_2_O + 0.1% FA (solvent A) and acetonitrile + 0.1% FA (solvent B), run over the stationary phase at a flow rate was set of 0.5 mL min^−1^. The liquid chromatography method followed a linear gradient from 0 to 0.5 min, 5% B, 0.5 to 8 min, 5% to 50% B, 8 to 10 min, 50% to 99% B, followed by a 2-min washout phase at 99% B and a 3-min re-equilibration phase at 5% B.

A Q-Exactive Orbitrap (Thermo Fisher Scientific) was used for MS/MS analysis using electrospray ionization (ESI) in positive mode. Data-dependent acquisition (DDA) of MS/MS spectra was performed in positive mode. ESI parameters were set to 52 AU sheath gas flow, 14 AU auxiliary gas flow, 0 L min^−1^ sweep gas flow, and 400°C auxiliary gas temperature. The spray voltage was set to 3.5 kV and the inlet capillary to 320°C. Then, 50-V S-lens level was applied. MS scan range was set to 150–1500 m/z with a resolution at m/z 200 (*R*_m/z 200_) of 70,000 with one microscan. The maximum ion injection time was set to 100 ms with automated gain control (AGC) target of 1.0 × 10^6^. Up to 5 MS/MS spectra per MS1 survey scan were recorded in DDA mode with R_m/z 200_ of 17,500 with one microscan. The maximum ion injection time for MS/MS scans was set to 100 ms with an AGC target of 3.0 × 10^5^ ions and a minimum 5% C-trap filling. The MS/MS precursor isolation window was set to m/z 1. Normalized collision energy was set to a stepwise increase from 20% to 30% to 40% with *z* = 1 as default charge state. MS/MS scans were triggered at the apex of chromatographic peaks within 2–15 s from their first occurrence. Dynamic precursor exclusion was set to 5 s. Ions with unassigned charge states were excluded from MS/MS acquisition as well as isotope peaks. 100 charge states were excluded from MS/MS acquisition as well as isotope peaks. Samples were run alongside a domoic acid standard (Canadian Reference Materials, Halifax, Canada) to verify retention time and fragmentation.

Chromatography drift and fragmentation quality were verified with a quality control mix of six standards run at the beginning and end of the sequence: Sulfamethazine, Sulfamethizole, Sulfachloropyridazine, Sulfadimethoxine, Amitryptiline, and Coumarin-314. Solvent blanks were run to identify background noise, and an IODA exclusion list was generated and applied to the MS method to prevent fragmentation of MS1 peaks present in the blanks [67].

**Data Conversion and Public Access**

Raw data from the Q-Exactive Orbitrap mass spectrometer (Thermo Fisher Scientific) was converted from the .raw format to the universal format .mzml using MS Convert with a 32-bit binary encoding precision and with peak picking applied to request centroid conversion. All .raw and .mzml files were uploaded to the MASSIVE database as public datasets for data sharing purposes.

**Classical Molecular Networking**

Data was run through a Classical Molecular Networking job using GNPS [58]. A molecular network was created using the online workflow (https://ccms-ucsd.github.io/GNPSDocumentation/) on the GNPS website (http://gnps.ucsd.edu). The precursor ion mass tolerance was set to 0.01 Da and a MS/MS fragment ion tolerance of 0.01 Da. A network was then created where edges were filtered to have a cosine score above 0.5 and more than 5 matched peaks. Further, edges between two nodes were kept in the network if each of the nodes appeared in each other's respective top 10 most similar nodes. Finally, the maximum size of a molecular family was set to 100, and the lowest scoring edges were removed from molecular families until the molecular family size was below this threshold. The spectra in the network were then searched against GNPS' spectral libraries. The library spectra were filtered in the same manner as the input data. All matches kept between network spectra and library spectra were required to have a score above 0.5 and at least 5 matched peaks.

**Data Deconvolution**

Feature finding was performed using MZMine 3 [68–70]. The processing wizard workflow was used to define parameters: the MS1 cutoff was set at 5.0E4, the MS2 cutoff at 1.0E3, and the minimum feature height at 1.0E5. The scan-to-scan m/z tolerance was set to 10 ppm, the feature-to-feature m/z tolerance was set to 3 ppm, and the sample-to- sample m/z tolerance to 5 ppm. HPLC Parameters included a cutoff of the first 0.3 minutes before injection, and the last 5 minutes of washing, and the maximum peaks per chromatogram was set to 15. Post-processing data was exported without smoothing, and with outputs for: feature based molecular networking, ion identity networking, and Sirius. The parameters used in MZmine 3 are listed in (**Table S2).**

**Metadata**

Metadata was compiled using ReDU [71].

**Feature Based Molecular Networking and Ion Identity Networking**

A molecular network was created with the Feature-Based Molecular Networking (FBMN) workflow [58, 72]. The mass spectrometry data were first processed with MZMINE2 and the results were exported to GNPS for FBMN analysis [69]. The data was filtered by removing all MS/MS fragment ions within +/- 17 Da of the precursor m/z. MS/MS spectra were window filtered by choosing only the top 6 fragment ions in the +/- 50 Da window throughout the spectrum. The precursor ion mass tolerance was set to 0.01 Da and the MS/MS fragment ion tolerance to 0.01 Da. A molecular network was then created where edges were filtered to have a cosine score above 0.7 and more than 6 matched peaks. Further, edges between two nodes were kept in the network if and only if each of the nodes appeared in each other’s respective top 10 most similar nodes. Finally, the maximum size of a molecular family was set to 100, and the lowest scoring edges were removed from molecular families until the molecular family size was below this threshold. The spectra in the network were then searched against GNPS spectral libraries [58, 73]. The library spectra were filtered in the same manner as the input data. All matches kept between network spectra and library spectra were required to have a score above 0.7 and at least 6 matched peaks. The DEREPLICATOR was used to annotate MS/MS spectra [74]. The molecular networks were visualized using Cytoscape software [75].

**Data Quality Checking Post-Processing**

Data was manually inspected to verify that features of interest were retained after data deconvolution and feature based molecular networking. Relevant technical blank and media blank signals were removed from all corresponding samples by using a blank subtraction application (Dorresteinappshub.ucsd.edu:5000). The output from MZMine 3 was inputted into the blank subtraction application, and the non-blank minimum threshold was set to 3. Datasets for which volume was not corrected for during the sample preparation volume-correction resuspension step were normalized by total ion chromatogram. The output quant table from MZMine 3 analysis was merged with the output feature table from feature based molecular networking analysis to determine relative abundances of annotated features. Post-processed data then was used for downstream analysis and plotting using R, Excel, and Python.

**Molecular Formula and Compound Class Predictions**

SIRIUS and CANOPUS were used to predict molecular formulas based on isotope patterns in MS spectra and MS/MS fragmentation patterns and compound classes based on MS/MS fragmentation pattern libraries, respectively [59, 60]. Limitations to this approach include biases in ionization efficiency and molecules that differentially bind the PPL during solid phase extraction. This approach therefore does not holistically reflect true total dissolved organic matter. However, changes that we do detect between samples are significant because all methods are consistent.

Molecular formula assignment of tandem mass spectrometry data was conducted with the SIRIUS computational annotation tool (version 4.0.1, linux build 5) on a cluster computer [76, 77]. In order to compute *de novo* molecular formulas by matching experimental and predicted isotopic patterns and by using fragmentation tree analysis of fragment ions, SIRIUS parameters were set as: molecular formula candidates retained was set to 20, and m/z >850 were excluded, profile was set to orbitrap, m/z maximum deviation to 10 ppm, all adducts were considered (–auto-charge), and adduct annotation from MZmine was trusted (–trust-ion-prediction). SIRIUS output was separated into each element in Excel, and this table was merged with the FBMN table and MZMine quant table. Blanks and standards were not analyzed by SIRIUS/CANOPUS, and nor were features without SIRIUS/CANOPUS annotations or annotations without quant information for those features. After this filtering, 7872 features remained of the initial 10416 features. Additional filtering was performed to remove features that were 0 in every sample, leaving 7552 features remaining. Elemental ratios of N:C, O:C, P:C, and H:C were calculated and used in analysis. Furthermore, nominal oxidation state of carbon (NOSC) assuming a net charge of 0 for each feature and standard Gibbs Potential Energy was calculated following described methods [78].

**Total Domoic Acid Analysis:**

Methods for whole water sampling for LC-MS/MS metabolomics and applications to environmental and ecological research have been described [62, 65]. The domoic acid feature was detected with FBMN and confirmed with retention time of a domoic acid standard that was run alongside at RT=3.25 minutes (Canadian Reference Materials). A standard curve of domoic acid was run non-concurrently to quantify domoic acid in the whole water samples. Six orders of magnitude of DA dilutions were run to identify an integrated peak area range of DA in the samples to identify a rough concentration range as an external titration. Subsequently, an internal titration standard addition methodology was conducted by adding different amounts of DA within that concentration range to the sample with the highest DA in the dataset, and calculated the change in integrated peak area with and without the DA to get the standard curve values. Quantitative analysis was performed in the GNPS Dashboard with extracted ion chromatogram m/z 312.12 +/- 0.01 for the mass of domoic acid in positive ion mode [61]. Integrated peak areas were exported as a .csv file. Methods for whole water sampling for LC-MS/MS metabolomics and applications to environmental and ecological research have been described [62, 63, 65].

**ESP Domoic Acid Analysis**

Toxin data were derived from both an Environmental Sample Processor (ESP) instrument on an autonomous underwater vehicle (AUV) that followed the high-biomass region. ESP methods for quantification particulate DA from the SPR, and ESP applications to HAB research in Monterey Bay and the Southern California Bight have been developed (Ussler, unpublished). In this study, 26 ESP samples were analyzed and 0 detected DA by SPR.

**Data Visualization:**

Data was visualized with Cytoscape 3.9.0, R Studio 2022.02.3 Build 492, Python Jupyter Notebooks, and Adobe Illustrator 27.0 2023 [75]. Correlations were visualized as correlograms using the CorRPlot package in R.

**Supplementary Text**

Functional Characterization of the Active Community

Weighted Gene Correlation Network Analysis (WGCNA) was used to define modules of co-expressed Open Reading Frames (ORFs) [56, 57]. 15,574 ORFs from ship surface, ship SCM, and 3G-ESP samples met statistical cut-offs and were included in analysis that identified seven modules of ORFs with similar expression profiles. (**Figs. S5h, S5i, S5j, S5k**). The largest module, “MEhoneydew1,” significantly positively correlated with samples from the 3G-ESP/LRAUV and with time (p<0.001) (**Fig. S5h**), while significantly negatively correlated with samples from the SCM and the Surface (p<0.001) (**Fig. S5h**). Genes highly associated with MEhoneydew1 include ORFs annotated as protein phosphatase, ATP synthase, NADP transhydrogenase, ribosomal proteins, bacteriorhodopsin, and DNA binding proteins. Genes highly associated with the second largest module “MEindianred4” included ORFs annotated as tropomyosin and troponin, ribosomal proteins, mitochondrial ATP synthase. The MEindianred4 module had a significant negative correlation with samples from the AUV, and a significant positive correlation with samples from the SCM. Genes highly associated with “MEblack” include ORFs annotated as ribosomal proteins (60S, 40S, 30S) and heat shock proteins. Genes highly associated with “MEdarkolivegreen4” included ORFs annotated as ribosomal proteins and translation factors. Genes highly associated with “MEmediumpurple2” included ORFs annotated as photosynthesis-related proteins, such as chlorophyll-binding proteins, thylakoid lumenal proteins, and serine proteases. MEpalevioletred2 had significant positive correlations with day progression (p<0.001) and samples from the AUV (p<0.01). Genes highly associated with “MEpalevioletred2” included ORFs annotated as ribosomal proteins (60S, 30S, 40S), cytochrome c oxidases, and heat shock proteins.

Taxonomic Structure of the Community Measured by Ship Sampling

DNA metabarcoding taxonomic assignments showed a diverse community of archaea, bacteria, and eukaryotes that were quite stable between samples of a depth classification, reflecting the sympatric nature of the region sampled, characteristic of the California Central Coast [79].

Phylogenetic analysis of 18SV9 rRNA amplicons revealed a community dominated by the phylum Metazoa which includes copepods at all depth classes (**Fig. S6a**). The second most abundant phylum was Dinoflagellata, followed by Ochrophyta which includes diatoms. The deep samples had a higher proportion of diatoms than did the surface and SCM samples, while the surface and SCM samples had a higher proportion of metazoans than did the deep samples. To avoid data misinterpretation due to 18S rRNA copy number variation, primer bias, or amplification efficiency bias, relative abundances of compositional data rather than absolute quantities were compared [80]. Nevertheless, read counts are significantly positively correlated with morphological abundance and carbon biomass, especially for highly abundant taxa [81–84]. The 18SV9 photosynthetic community was dominated by the phyla Dinoflagellata, Ochrophyta, and Haptophyta at all depth classes. (**Fig. S6b**). Dinoflagellata made up a higher proportion of the photosynthetic community in the deeper depths, whereas Ochrophyta made up a higher proportion of the photosynthetic community in the surface and SCM. PCA of the 18SV9 data based on the weighted unifrac distance metric suggested that surface samples are more similar to each other, and there was more diversity among the deep and SCM samples (**Fig. S6c**). Alpha diversity analysis (Shannon’s) suggests that there was higher diversity within in the deep depths than the surface and SCM (**Fig. S6d**).

Non-plastid 16S (V4-V5) rRNA amplicons included archaeal and bacterial amplicons. Phylogenetic analysis of 16S rRNA bacterial amplicons revealed a community dominated by the phylum Bacteroidota*,* followed by Acidobacteriota *and* Actinobacteriota at all depth classes (**Fig. S6e**). The proportion of dominance of phyla was similar among all three depth classes. The 16S community with photosynthetic potential was dominated by the phyla Acidobacteriota, Cyanobacteria, and Proteobacteria at all depth classes (**Fig. S6f**). Cyanobacteria made up a higher proportion of the photosynthetic community in the deeper depths, whereas Acidobacteriota made up a higher proportion of the photosynthetic community in the surface and SCM. PCA of the 16S non-plastid data (weighted unifrac) suggested that surface samples were more similar to each other, and there as more diversity among the deep and SCM samples (**Fig. S6g**. As with 18SV9, alpha diversity analysis (Shannon’s) suggested higher diversity in the deep than the surface and SCM (**Fig. S6h**), demonstrating synteny across microbes of all domains.

The 16S archaeal community was dominated significantly by the phylum Crenarchaeota in all samples at all depths, locations, and times, followed by Altiarchaeota and Nanoarchaeota.

Taxonomic Structure of the Community Measured by 3G-ESP/LRAUV Sampling

DNA metabarcoding taxonomic assignments from 3G-ESP/LRAUV sampling showed a wide range of prokaryotes and eukaryotes that were stable over space and time. The prokaryotic community from 16S metabarcoding did not change significantly over space, time of day, or day vs. night, nor did the bacterial community dominated by phyla Proteobacteria and Bacteriodota change over space or time (**Fig. S7a, S7b**). Along a cross-shore transect, alpha diversity peaked in the very nearshore and slowly increased again further offshore (**Fig. S7c**), however, with only two orders of archaea measured with 16S metabarcoding, archaeal diversity remained low at all stations (**Fig. S7d**). Spatial and temporal synteny also existed in the eukaryotic community as measured with 18SV9 metabarcoding (**Fig. S7e, Fig. S7f),** which identified Metazoa as the most abundant phylum across all samples. Unlike the 16S metabarcoding results, alpha diversity as lowest at the very nearshore stations yet evenly distributed across space and time at all other stations (**Fig. S7g**). Notably, *Pseudo-nitzschia* ITS2 amplicon abundance increased over the time of sampling, and amplicon sequencing resolved the following eight *Pseudo-nitzschia* species: *americana, australis, bucculenta, dolorosa, fraudulenta, fryxelliana, hasleana,* and *seriata* **(Fig. S7h).** The consistency of the community composition (16S, 18SV9, and ITS2) in both taxonomic composition and alpha and beta diversity reflected the quasi-Lagrangian sampling strategy. Because this was not a true Lagrangian drift, the alternative hypothesis is that the Bay was homogenous in community composition.

**References:**

1.	Omand MM, Cetinić I, Lucas AJ. Using bio-optics to reveal phytoplankton physiology from a Wirewalker autonomous platform. *Oceanography* 2017; **30**: 128–131.

2.	Truelove NK, Patin NV, Min M, Pitz KJ, Preston CM, Yamahara KM, et al. Expanding the temporal and spatial scales of environmental DNA research with autonomous sampling. *Environ DNA* 2022; **4**: 972–984.

3.	Yamahara KM, Preston CM, Birch J, Walz K, Marin R, Jensen S, et al. In situ Autonomous Acquisition and Preservation of Marine Environmental DNA Using an Autonomous Underwater Vehicle. *Front Mar Sci* 2019; **6**.

4.	Zhang Y, Ryan JP, Hobson BW, Kieft B, Romano A, Barone B, et al. A system of coordinated autonomous robots for Lagrangian studies of microbes in the oceanic deep chlorophyll maximum. *Sci Robot* 2021; **6**: eabb9138.

5.	Yentsch CS, Menzel DW. A method for the determination of phytoplankton chlorophyll and phaeophytin by fluorescence. *Deep Sea Res Oceanogr Abstr* 1963; **10**: 221–231.

6.	Holm-Hansen O, Lorenzen CJ, Holmes RW, Strickland JDH. Fluorometric Determination of Chlorophyll. *ICES J Mar Sci* 1965; **30**: 3–15.

7.	Lorenzen CJ. Determination of Chlorophyll and Pheo-Pigments: Spectrophotometric Equations1. *Limnol Oceanogr* 1967; **12**: 343–346.

8.	Timothy Pennington J, Chavez FP. Seasonal fluctuations of temperature, salinity, nitrate, chlorophyll and primary production at station H3/M1 over 1989–1996 in Monterey Bay, California. *Deep Sea Res Part II Top Stud Oceanogr* 2000; **47**: 947–973.

9.	Sakamoto CM, Friederich GE, Codispoti LA. MBARI procedures for automated nutrient analyses using a modified Alpkem Series 300 Rapid Flow Analyzer. *http://aquaticcommons.org/id/eprint/1971* 1990.

10.	James CC, Barton AD, Allen LZ, Lampe RH, Rabines A, Schulberg A, et al. Influence of nutrient supply on plankton microbiome biodiversity and distribution in a coastal upwelling region. *Nat Commun* 2022; **13**: 2448.

11.	Rabines A, Lampe R, Allen A. Sterivex DNA extraction. *protocols.io* 2020.

12.	Lin Y, Gifford S, Ducklow H, Schofield O, Cassar N. Towards Quantitative Microbiome Community Profiling Using Internal Standards. *Appl Environ Microbiol* 2019; **85**: e02634-18.

13.	Rabines A, Lampe R, Allen A. Amplicon Library Preparation. *protocols.io* 2020.

14.	Amaral-Zettler LA, McCliment EA, Ducklow HW, Huse SM. A Method for Studying Protistan Diversity Using Massively Parallel Sequencing of V9 Hypervariable Regions of Small-Subunit Ribosomal RNA Genes. *PLOS ONE* 2009; **4**: e6372.

15.	Parada AE, Needham DM, Fuhrman JA. Every base matters: assessing small subunit rRNA primers for marine microbiomes with mock communities, time series and global field samples. *Environ Microbiol* 2016; **18**: 1403–1414.

16.	Balzano S, Abs E, Leterme S. Protist diversity along a salinity gradient in a coastal lagoon. *Aquat Microb Ecol* 2015; **74**: 263–277.

17.	Lim HC, Tan SN, Teng ST, Lundholm N, Orive E, David H, et al. Phylogeny and species delineation in the marine diatom *Pseudo-nitzschia* (Bacillariophyta) using cox1, LSU, and ITS2 rRNA genes: A perspective in character evolution. *J Phycol* 2018; **54**: 234–248.

18.	H. Utermöhl. Zur Vervollkommung der quantitativen Phytoplankton Methodik. *Mitteilungen der Internationalen Vereinigung der Limnologen* 1958; **9**: 1–39.

19.	Lim H-C, Leaw C-P, Su SN-P, Teng S-T, Usup G, Mohammad-Noor N, et al. Morphology and Molecular Characterization of *Pseudo-Nitzschia* (bacillariophyceae) from Malaysian Borneo, Including the New Species *Pseudo-Nitzschia* Circumpora Sp. Nov. *J Phycol* 2012; **48**: 1232–1247.

20.	Lundholm N, Hasle GR, Fryxell GA, Hargraves PE. Morphology, phylogeny and taxonomy of species within the *Pseudo-nitzschia americana* complex (Bacillariophyceae) with descriptions of two new species, *Pseudo-nitzschia brasiliana* and *Pseudo-nitzschia linea*. *Phycologia* 2002; **41**: 480–497.

21.	Fitzpatrick E, Caron DA, Schnetzer A. Development and environmental application of a genus-specific quantitative PCR approach for *Pseudo-nitzschia* species. *Mar Biol* 2010; **157**: 1161–1169.

22.	Greenfield DI, Marin III R, Doucette GJ, Mikulski C, Jones K, Jensen S, et al. Field applications of the second-generation Environmental Sample Processor (ESP) for remote detection of harmful algae: 2006-2007. *Limnol Oceanogr Methods* 2008; **6**: 667–679.

23.	Bowers HA, Marin R, Birch JM, Scholin CA. Sandwich hybridization probes for the detection of *Pseudo-nitzschia* (Bacillariophyceae) species: An update to existing probes and a description of new probes. *Harmful Algae* 2017; **70**: 37–51.

24.	Miller PE, Scholin CA. Identification and enumeration of cultured and wild *Pseudo-nitzschia* (*Bacillariophyceae*) using species-specific LSU rRNA-targeted fluorescent probes and filter-based whole cell hybridization. *J Phycol* 1998; **34**: 371–382.

25.	Scholin C, Miller P, Buck K, Chavez F, Harris P, Haydock P, et al. Detection and quantification of *Pseudo-nitzschia australis* in cultured and natural populations using LSU rRNA-targeted probes. *Limnol Oceanogr* 1997; **42**: 1265–1272.

26.	Hubbard KA, Rocap G, Armbrust EV. Inter- and Intraspecific Community Structure Within the Diatom Genus *Pseudo-Nitzschia* (*Bacillariophyceae*) *J Phycol* 2008; **44**: 637–649.

27.	Lim HC, Tan SN, Teng ST, Lundholm N, Orive E, David H, et al. Phylogeny and species delineation in the marine diatom *Pseudo-nitzschia* (Bacillariophyta) using cox1, LSU, and ITS2 rRNA genes: A perspective in character evolution. *J Phycol* 2018; **54**: 234–248.

28.	Bolyen E, Rideout JR, Dillon MR, Bokulich NA, Abnet CC, Al-Ghalith GA, et al. Reproducible, interactive, scalable and extensible microbiome data science using QIIME 2. *Nat Biotechnol* 2019; **37**: 852–857.

29.	Martin M. Cutadapt removes adapter sequences from high-throughput sequencing reads. *EMBnet.journal* 2011; **17**: 10–12.

30.	Callahan BJ, McMurdie PJ, Rosen MJ, Han AW, Johnson AJA, Holmes SP. DADA2: High-resolution sample inference from Illumina amplicon data. *Nat Methods* 2016; **13**: 581–583.

31.	Guillou L, Bachar D, Audic S, Bass D, Berney C, Bittner L, et al. The Protist Ribosomal Reference database (PR2): a catalog of unicellular eukaryote small sub-unit rRNA sequences with curated taxonomy. *Nucleic Acids Res* 2013; **41**: D597-604.

32.	Yilmaz P, Parfrey LW, Yarza P, Gerken J, Pruesse E, Quast C, et al. The SILVA and ‘All-species Living Tree Project (LTP)’ taxonomic frameworks. *Nucleic Acids Res* 2014; **42**: D643-648.

33.	Ankenbrand MJ, Keller A, Wolf M, Schultz J, Förster F. ITS2 Database V: Twice as Much. *Mol Biol Evol* 2015; **32**: 3030–3032.

34.	Pruesse E, Quast C, Knittel K, Fuchs BM, Ludwig W, Peplies J, et al. SILVA: a comprehensive online resource for quality checked and aligned ribosomal RNA sequence data compatible with ARB. *Nucleic Acids Res* 2007; **35**: 7188–7196.

35.	Decelle J, Romac S, Stern RF, Bendif EM, Zingone A, Audic S, et al. PhytoREF: a reference database of the plastidial 16S rRNA gene of photosynthetic eukaryotes with curated taxonomy. *Mol Ecol Resour* 2015; **15**: 1435–1445.

36.	McMurdie PJ, Holmes S. phyloseq: An R Package for Reproducible Interactive Analysis and Graphics of Microbiome Census Data. *PLOS ONE* 2013; **8**: e61217.

37.	Dixon P. VEGAN, A Package of R Functions for Community Ecology. *J Veg Sci* 2003; **14**: 927–930.

38.	Cohen NR, Alexander H, Krinos AI, Hu SK, Lampe RH. Marine Microeukaryote Metatranscriptomics: Sample Processing and Bioinformatic Workflow Recommendations for Ecological Applications. *Front Mar Sci* 2022; **9**.

39.	Chen S, Zhou Y, Chen Y, Gu J. fastp: an ultra-fast all-in-one FASTQ preprocessor. *Bioinforma Oxf Engl* 2018; **34**: i884–i890.

40.	Bolger AM, Lohse M, Usadel B. Trimmomatic: a flexible trimmer for Illumina sequence data. *Bioinformatics* 2014; **30**: 2114–2120.

41.	Li D, Liu C-M, Luo R, Sadakane K, Lam T-W. MEGAHIT: an ultra-fast single-node solution for large and complex metagenomics assembly via succinct de Bruijn graph. *Bioinformatics* 2015; **31**: 1674–1676.

42.	Fu L, Niu B, Zhu Z, Wu S, Li W. CD-HIT: accelerated for clustering the next-generation sequencing data. *Bioinforma Oxf Engl* 2012; **28**: 3150–3152.

43.	Rho M, Tang H, Ye Y. FragGeneScan: predicting genes in short and error-prone reads. *Nucleic Acids Res* 2010; **38**: e191.

44.	Aramaki T, Blanc-Mathieu R, Endo H, Ohkubo K, Kanehisa M, Goto S, et al. KofamKOALA: KEGG Ortholog assignment based on profile HMM and adaptive score threshold. *Bioinforma Oxf Engl* 2020; **36**: 2251–2252.

45.	Buchfink B, Xie C, Huson DH. Fast and sensitive protein alignment using DIAMOND. *Nat Methods* 2015; **12**: 59–60.

46.	Cantalapiedra CP, Hernández-Plaza A, Letunic I, Bork P, Huerta-Cepas J. eggNOG-mapper v2: Functional Annotation, Orthology Assignments, and Domain Prediction at the Metagenomic Scale. *Mol Biol Evol* 2021; **38**: 5825–5829.

47.	Huerta-Cepas J, Szklarczyk D, Heller D, Hernández-Plaza A, Forslund SK, Cook H, et al. eggNOG 5.0: a hierarchical, functionally and phylogenetically annotated orthology resource based on 5090 organisms and 2502 viruses. *Nucleic Acids Res* 2019; **47**: D309–D314.

48.	Jones P, Binns D, Chang H-Y, Fraser M, Li W, McAnulla C, et al. InterProScan 5: genome-scale protein function classification. *Bioinforma Oxf Engl* 2014; **30**: 1236–1240.

49.	Delaye L, Vargas C, Latorre A, Moya A. Inferring Horizontal Gene Transfer with DarkHorse, Phylomizer, and ETE Toolkits. *Methods Mol Biol Clifton NJ* 2020; **2075**: 355–369.

50.	Langmead B, Salzberg SL. Fast gapped-read alignment with Bowtie 2. *Nat Methods* 2012; **9**: 357–359.

51.	Satinsky BM, Gifford SM, Crump BC, Moran MA. Use of internal standards for quantitative metatranscriptome and metagenome analysis. *Methods Enzymol* 2013; **531**: 237–250.

52.	Brunson JK, McKinnie SMK, Chekan JR, McCrow JP, Miles ZD, Bertrand EM, et al. Biosynthesis of the neurotoxin domoic acid in a bloom-forming diatom. *Science* 2018; **361**: 1356–1358.

53.	Steele TS, Brunson JK, Maeno Y, Terada R, Allen AE, Yotsu-Yamashita M, et al. Domoic acid biosynthesis in the red alga Chondria armata suggests a complex evolutionary history for toxin production. *Proc Natl Acad Sci* 2022; **119**: e2117407119.

54.	Edgar RC. Muscle5: High-accuracy alignment ensembles enable unbiased assessments of sequence homology and phylogeny. *Nat Commun* 2022; **13**: 6968.

55.	Minh BQ, Schmidt HA, Chernomor O, Schrempf D, Woodhams MD, von Haeseler A, et al. IQ-TREE 2: New Models and Efficient Methods for Phylogenetic Inference in the Genomic Era. *Mol Biol Evol* 2020; **37**: 1530–1534.

56.	Langfelder P, Horvath S. WGCNA: an R package for weighted correlation network analysis. *BMC Bioinformatics* 2008; **9**: 559.

57.	Zhang B, Horvath S. A general framework for weighted gene co-expression network analysis. *Stat Appl Genet Mol Biol* 2005; **4**: Article17.

58.	Wang M, Carver JJ, Phelan VV, Sanchez LM, Garg N, Peng Y, et al. Sharing and community curation of mass spectrometry data with GNPS. *Nat Biotechnol* 2016; **34**: 828–837.

59.	Dührkop K, Fleischauer M, Ludwig M, Aksenov AA, Melnik AV, Meusel M, et al. SIRIUS 4: a rapid tool for turning tandem mass spectra into metabolite structure information. *Nat Methods* 2019; **16**: 299–302.

60.	Dührkop K, Nothias L-F, Fleischauer M, Reher R, Ludwig M, Hoffmann MA, et al. Systematic classification of unknown metabolites using high-resolution fragmentation mass spectra. *Nat Biotechnol* 2021; **39**: 462–471.

61.	Petras D, Phelan VV, Acharya D, Allen AE, Aron AT, Bandeira N, et al. GNPS Dashboard: collaborative exploration of mass spectrometry data in the web browser. *Nat Methods* 2022; **19**: 134–136.

62.	Petras D, Koester I, Da Silva R, Stephens BM, Haas AF, Nelson CE, et al. High-resolution liquid chromatography tandem mass spectrometry enables large scale molecular characterization of dissolved organic matter. *Front Mar Sci* 2017; **4**: 405.

63.	Thukral M, Koester I, Petras D, Torres RR, Aron A, Gentry E, et al. Protocol for PPL Solid Phase Extraction for Dissolved Organic Matter (DOM) Sample Preparation for LC-MS/MS Analysis. *protocols.io* 2022.

64.	Dittmar T, Koch B, Hertkorn N, Kattner G. A simple and efficient method for the solid-phase extraction of dissolved organic matter (SPE-DOM) from seawater. *Limnol Oceanogr Methods* 2008; **6**: 230–235.

65.	Cancelada L, Torres RR, Garrafa Luna J, Dorrestein PC, Aluwihare LI, Prather KA, et al. Assessment of styrene-divinylbenzene polymer (PPL) solid-phase extraction and non-targeted tandem mass spectrometry for the analysis of xenobiotics in seawater. *Limnol Oceanogr Methods* 2022; **20**: 89–101.

66.	Petras D, Minich JJ, Cancelada LB, Torres RR, Kunselman E, Wang M, et al. Non-targeted tandem mass spectrometry enables the visualization of organic matter chemotype shifts in coastal seawater. *Chemosphere* 2021; **271**: 129450.

67.	Zuo Z, Cao L, Nothia L-F, Mohimani H. MS2Planner: improved fragmentation spectra coverage in untargeted mass spectrometry by iterative optimized data acquisition. *Bioinformatics* 2021; **37**: i231–i236.

68.	Schmid R, Heuckeroth S, Korf A, Smirnov A, Myers O, Dyrlund TS, et al. Integrative analysis of multimodal mass spectrometry data in MZmine 3. *Nat Biotechnol* 2023; 1–3.

69.	Pluskal T, Castillo S, Villar-Briones A, Orešič M. MZmine 2: Modular framework for processing, visualizing, and analyzing mass spectrometry-based molecular profile data. *BMC Bioinformatics* 2010; **11**: 395.

70.	Katajamaa M, Miettinen J, Orešič M. MZmine: toolbox for processing and visualization of mass spectrometry based molecular profile data. *Bioinformatics* 2006; **22**: 634–636.

71.	Jarmusch AK, Wang M, Aceves CM, Advani RS, Aguirre S, Aksenov AA, et al. ReDU: a framework to find and reanalyze public mass spectrometry data. *Nat Methods* 2020; **17**: 901–904.

72.	Nothias L-F, Petras D, Schmid R, Dührkop K, Rainer J, Sarvepalli A, et al. Feature-based molecular networking in the GNPS analysis environment. *Nat Methods* 2020; **17**: 905–908.

73.	Horai H, Arita M, Kanaya S, Nihei Y, Ikeda T, Suwa K, et al. MassBank: a public repository for sharing mass spectral data for life sciences. *J Mass Spectrom* 2010; **45**: 703–714.

74.	Mohimani H, Gurevich A, Shlemov A, Mikheenko A, Korobeynikov A, Cao L, et al. Dereplication of microbial metabolites through database search of mass spectra. *Nat Commun* 2018; **9**: 4035.

75.	Shannon P, Markiel A, Ozier O, Baliga NS, Wang JT, Ramage D, et al. Cytoscape: A Software Environment for Integrated Models of Biomolecular Interaction Networks. *Genome Res* 2003; **13**: 2498–2504.

76.	Böcker S, Letzel MC, Lipták Z, Pervukhin A. SIRIUS: decomposing isotope patterns for metabolite identification†. *Bioinformatics* 2009; **25**: 218–224.

77.	Böcker S, Dührkop K. Fragmentation trees reloaded. *J Cheminformatics* 2016; **8**: 5.

78.	Wegley Kelly L, Nelson CE, Petras D, Koester I, Quinlan ZA, Arts MGI, et al. Distinguishing the molecular diversity, nutrient content, and energetic potential of exometabolomes produced by macroalgae and reef-building corals. *Proc Natl Acad Sci* 2022; **119**.

79.	Kolody BC, McCrow JP, Allen LZ, Aylward FO, Fontanez KM, Moustafa A, et al. Diel transcriptional response of a California Current plankton microbiome to light, low iron, and enduring viral infection. *ISME J* 2019; **13**: 2817–2833.

80.	Potvin M, Lovejoy C. PCR-Based Diversity Estimates of Artificial and Environmental 18S rRNA Gene Libraries. *J Eukaryot Microbiol* 2009; **56**: 174–181.

81.	Harvey JBJ, Ryan JP, Zhang Y. Influences of extreme upwelling on a coastal retention zone. *Front Mar Sci* 2021; **8**: 648994.

82.	Hirai J, Kuriyama M, Ichikawa T, Hidaka K, Tsuda A. A metagenetic approach for revealing community structure of marine planktonic copepods. *Mol Ecol Resour* 2015; **15**: 68–80.

83.	Bucklin A, Yeh HD, Questel JM, Richardson DE, Reese B, Copley NJ, et al. Time-series metabarcoding analysis of zooplankton diversity of the NW Atlantic continental shelf. *ICES J Mar Sci* 2019; **76**: 1162–1176.

84.	McLaren MR, Willis AD, Callahan BJ. Consistent and correctable bias in metagenomic sequencing experiments. *eLife* 2019; **8**: e46923.

### **Supplementary Figures**

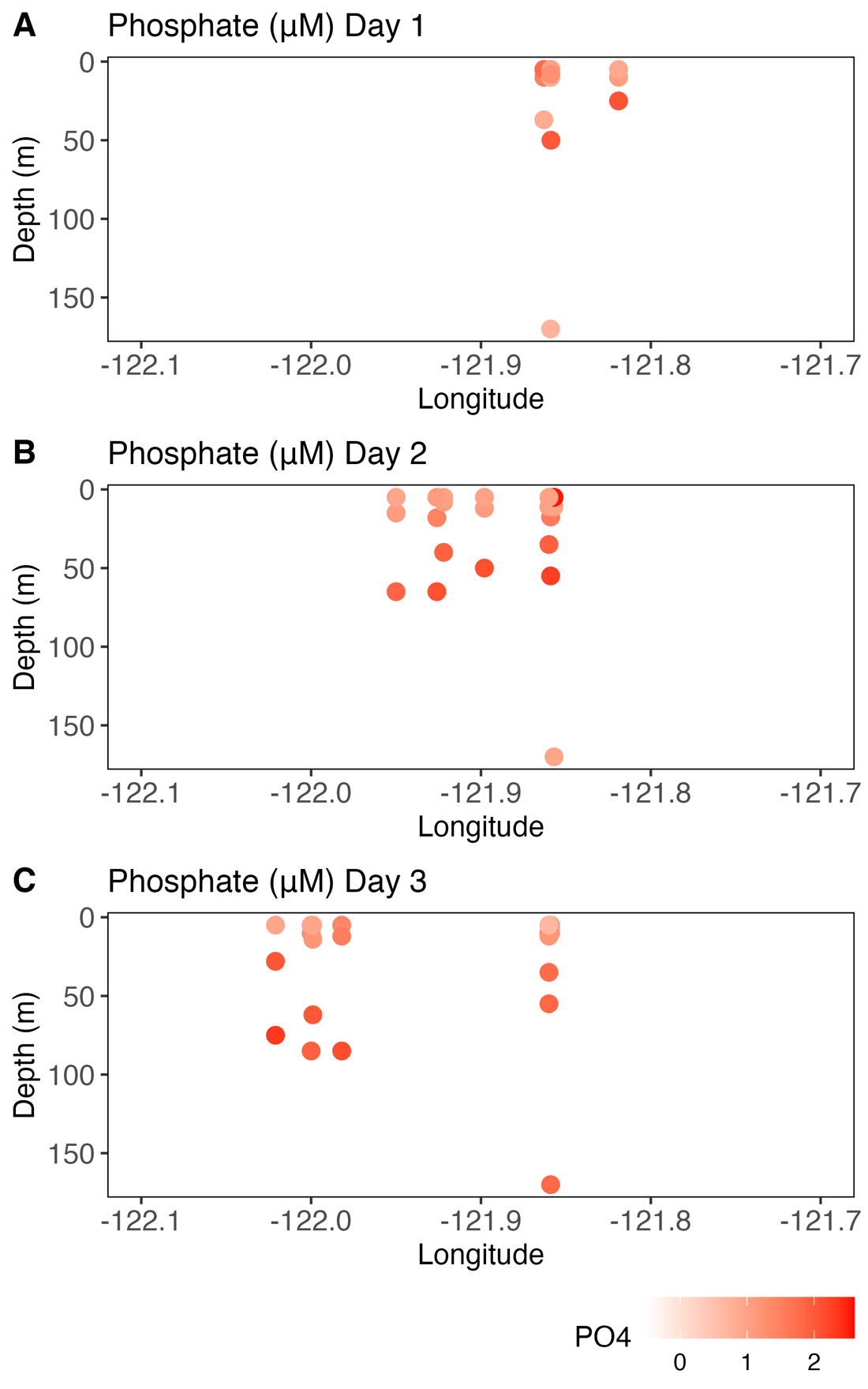

**Figure S1a.** Phosphate (µM) from ship sampling stations from (A) Day 1, (B) Day 2, and (C) Day 3.

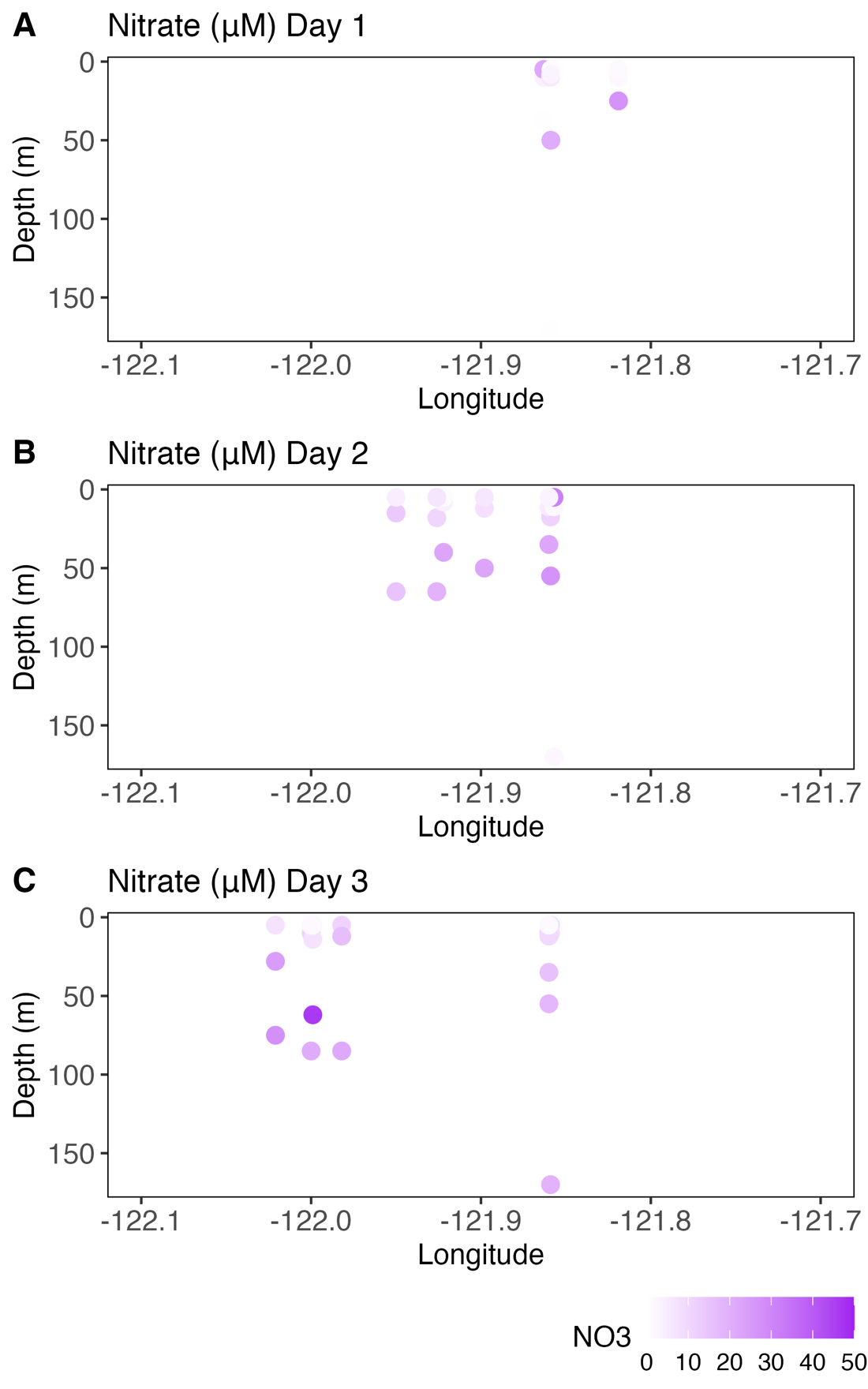

**Figure S1b.** Nitrate (µM) from ship sampling stations from (A) Day 1, (B) Day 2, and (C) Day 3. Plots show depth vs Longitude and darker colors indicate higher concentrations.

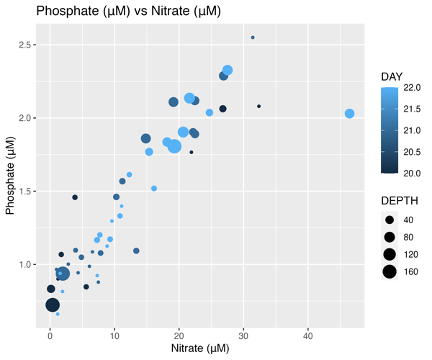

**Figure S1c.** Phosphate (µM) vs nitrate (µM) from ship sampling stations reveals a linear relationship across all days and depths sampled. The size of the dot indicates depth, and the color indicates day, with lighter colors indicating later days.

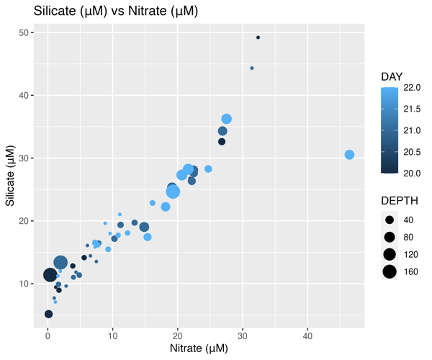

**Figure S1d.** Silicate (µM) vs nitrate (µM) from ship sampling stations reveals a linear relationship across all days and depths sampled. The size of the dot indicates depth, and the color indicates day, with lighter colors indicating later days.

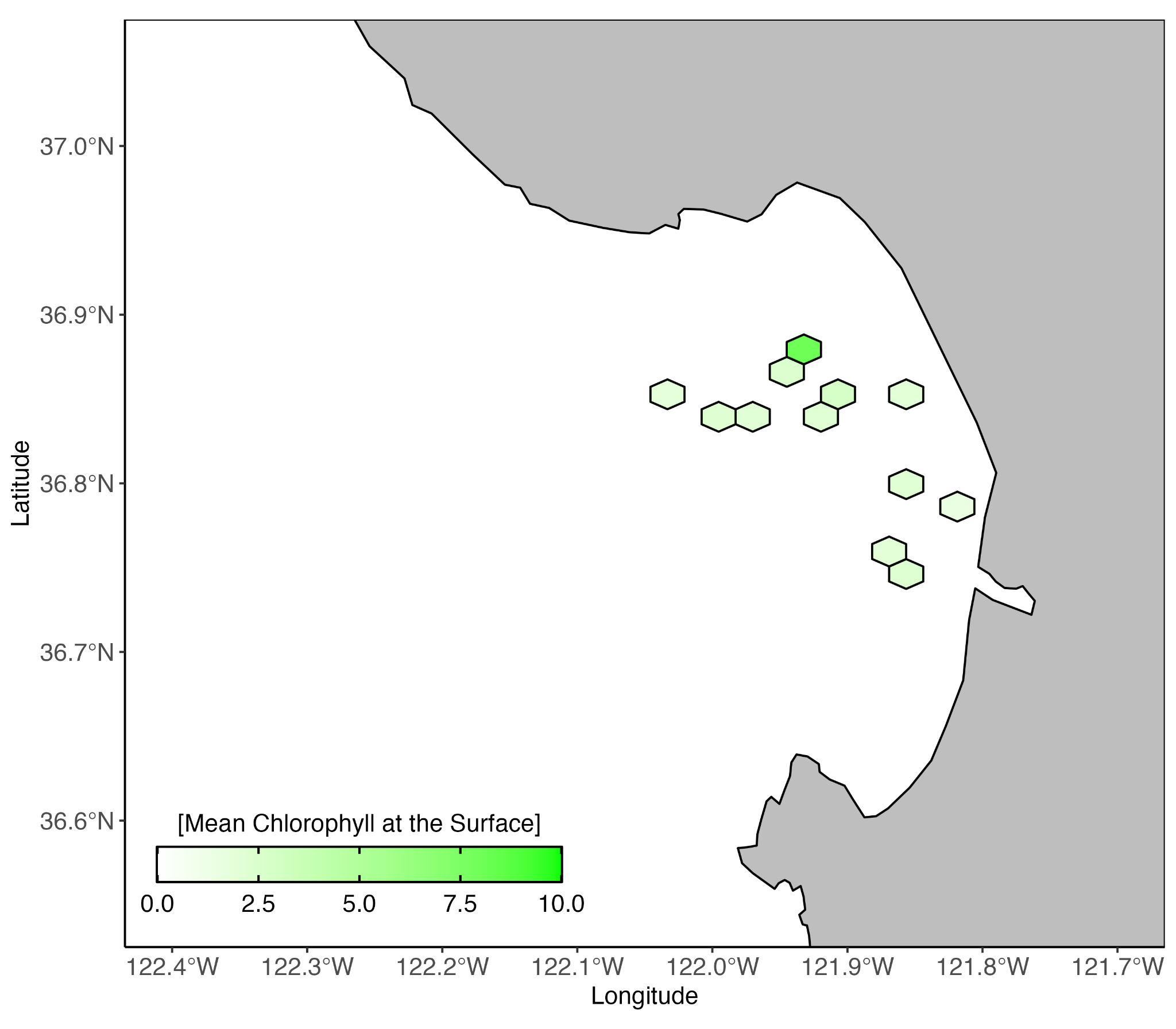

**Figure S1e.** Mean chlorophyll *a* (µg/L) at the surface from ship sampling stations. Chlorophyll values from samples taken within a hexagonal area are averaged.

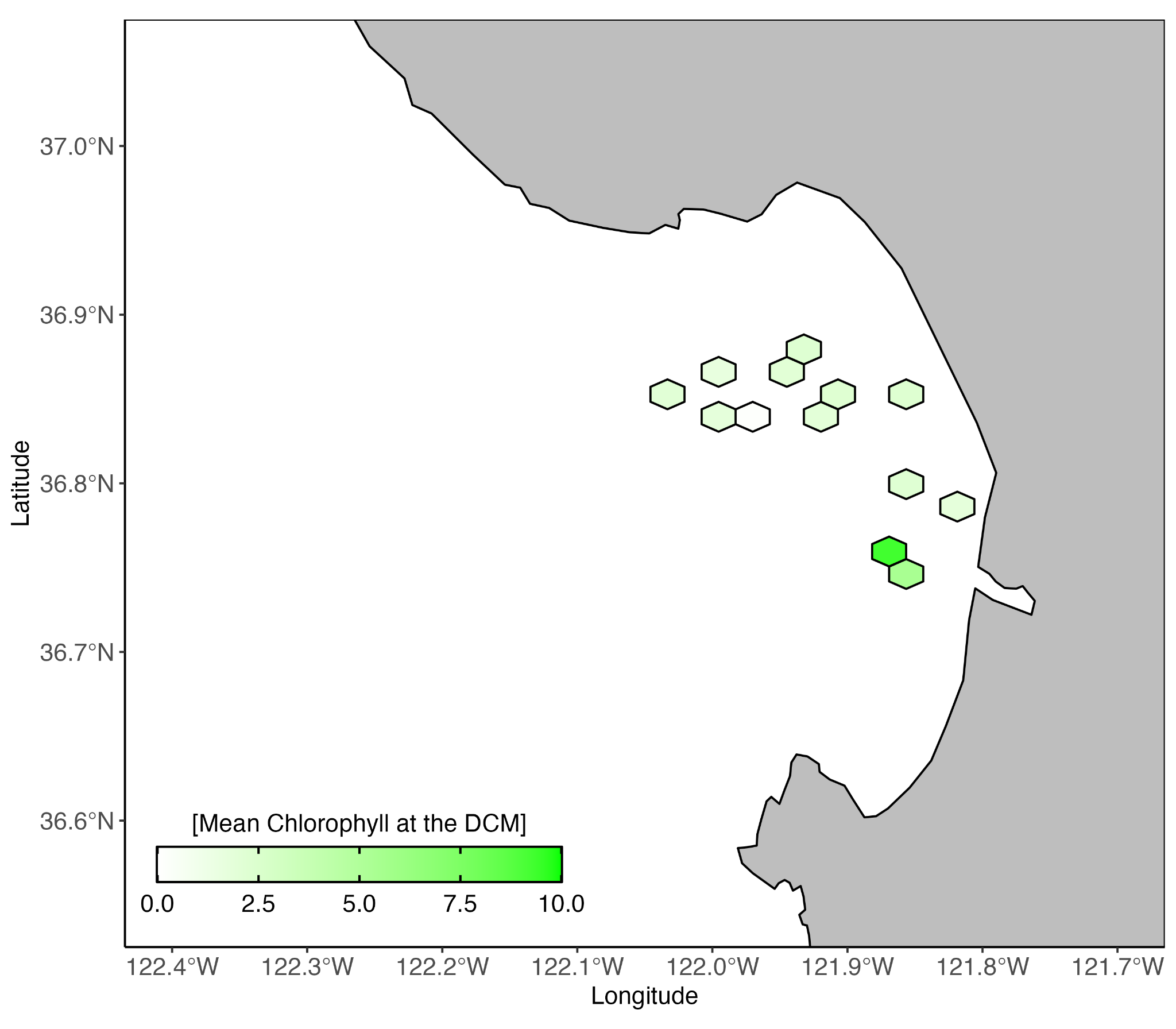

**Figure S1f.** Mean chlorophyll *a* (µg/L) at SCM from ship sampling stations. Chlorophyll values from samples taken within a hexagonal area are averaged.
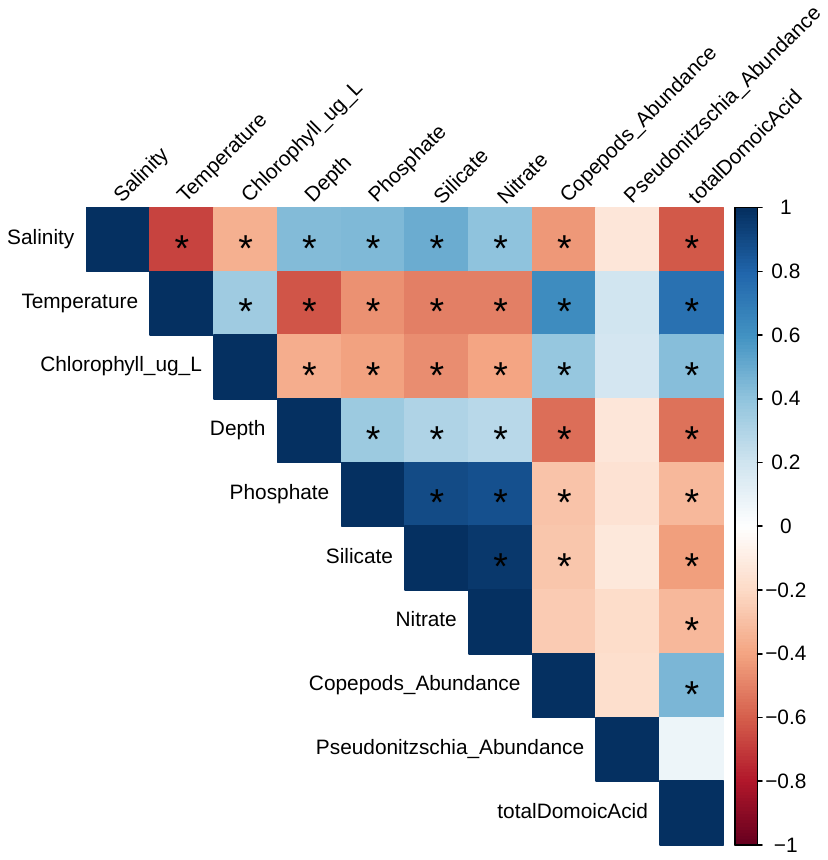

**Figure S1g**. Spearman rank correlations identified relationships between environmental variables, where values closer to 1 (dark blue) are positive correlations, and values closer to −1 (dark red) are negative correlations, and an asterisk (*) represents significant correlations (P < 0.05). Some correlations include positive significant correlations between total domoic acid and copepod abundance, temperature, and chlorophyll *a*, while a significant negative correlation between total domoic acid and salinity.

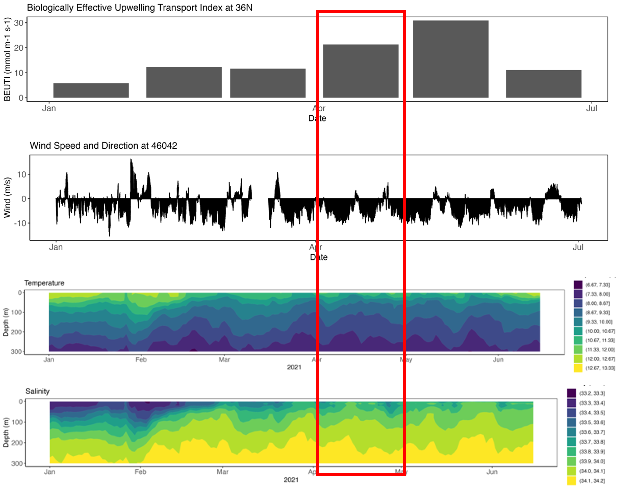

**Figure S1h.** Environmental Context and Regional Upwelling Conditions of Monterey Bay, CA, USA during January through June of 2021. (Top) Monthly upwelling index at 36°N, (Middle) wind speed from National Data Buoy Center (NDBC) Station 46042; negative values are upwelling favorable. (Bottom) Daily mean temperature and salinity from MBARI Mooring M1. The month of April is indicated with a red box, during which the field campaign took place.

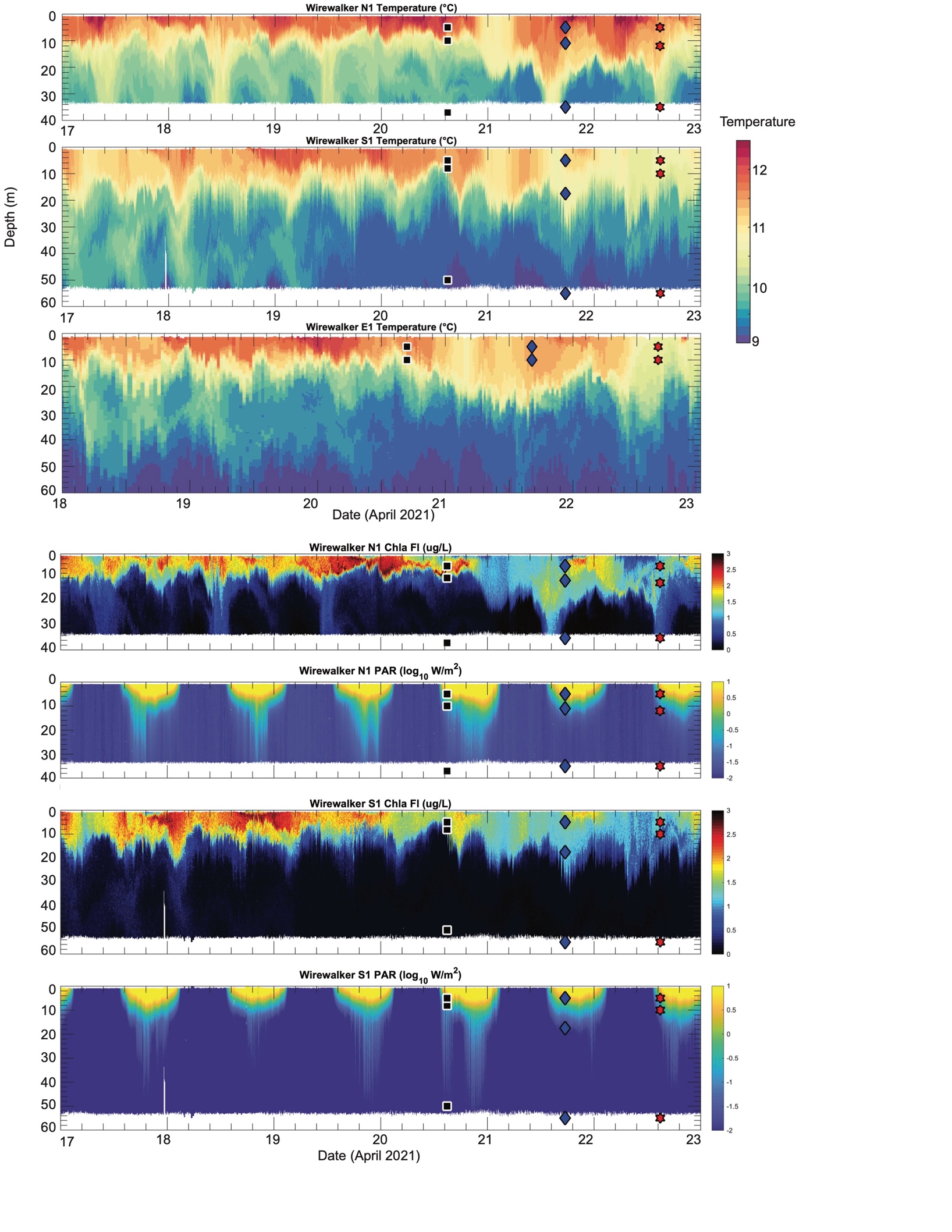

**Figure S2a.** Measurements Wirewalker instruments: Chlorophyll-*a* fluorescence, temperature, and PAR.

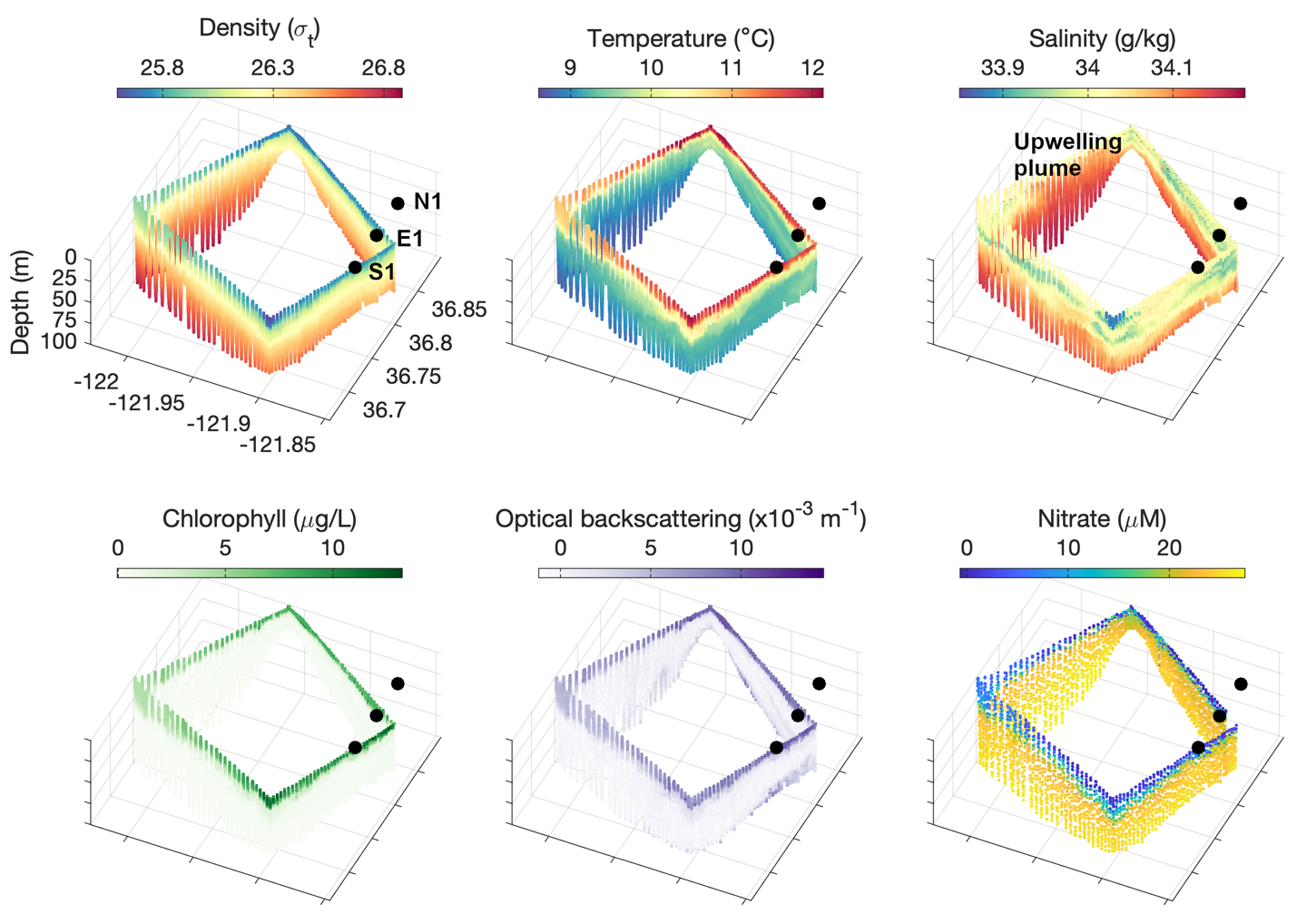

**Figure S2b.** AUV Survey in Monterey Bay from April 19 to April 20, 2021. The upwelling plume entering the study area from the northwest is identifiable by elevated salinity. Surface locations of the Wirewalker stations are indicated in the density plot.

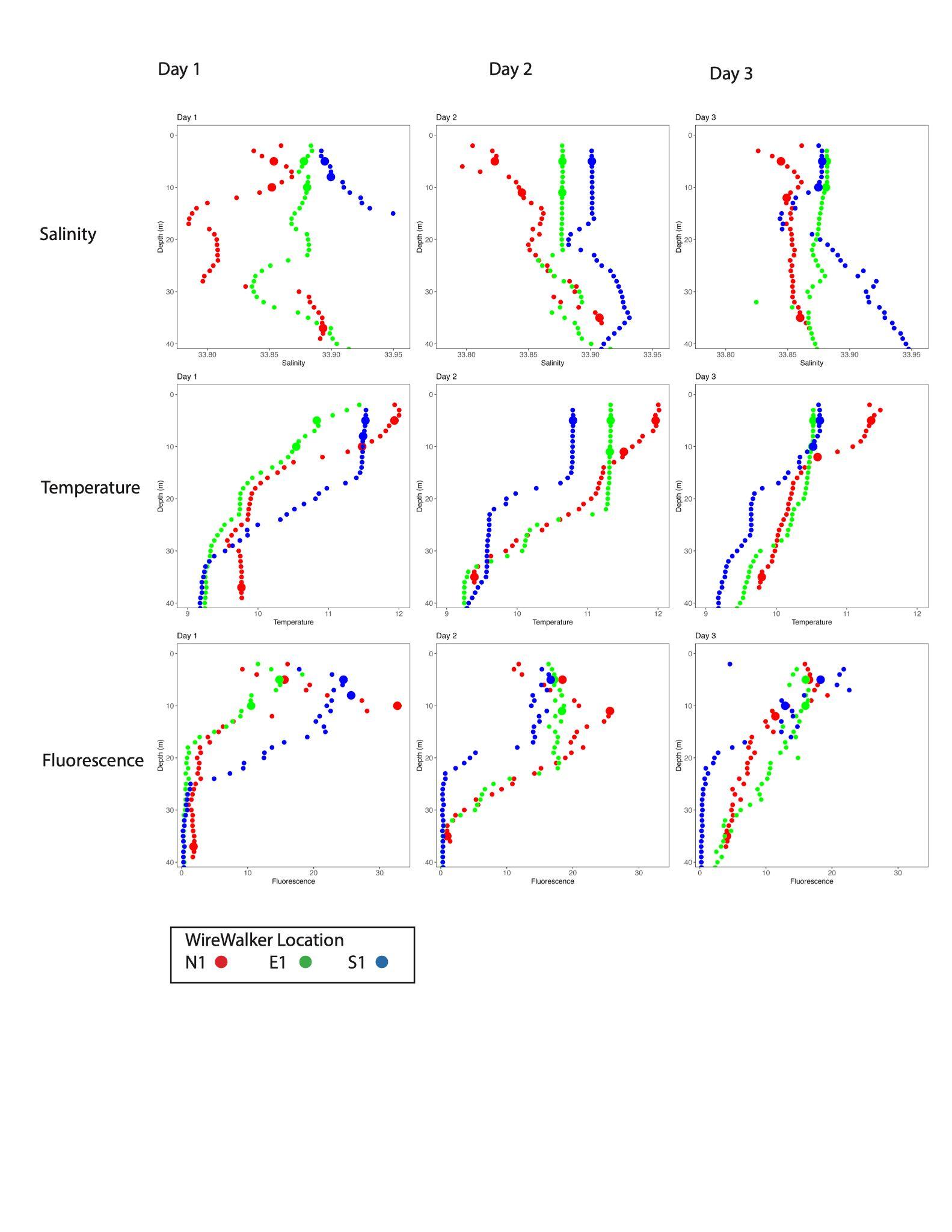

**Figure S2c.** CTD Profiles of Salinity, Temperature, and Fluorescence from Ship CTD casts in Eulerian sampling stations (N1, E1, and S1) over all three days.

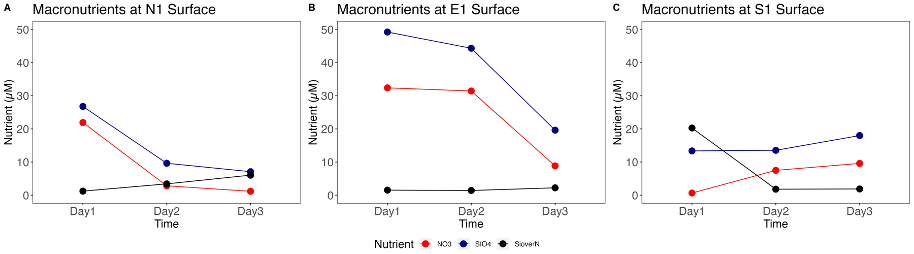

**Figure S2d.** Macronutrients Nitrate, Silicate, and Silicate to Nitrate ratio (µM) at the Surface of the three Eulerian sampling stations over the three days of ship sampling. Measurements were taken from discrete CTD water samples at the time of ship sampling. Station E1 has the highest absolute nutrient concentrations.

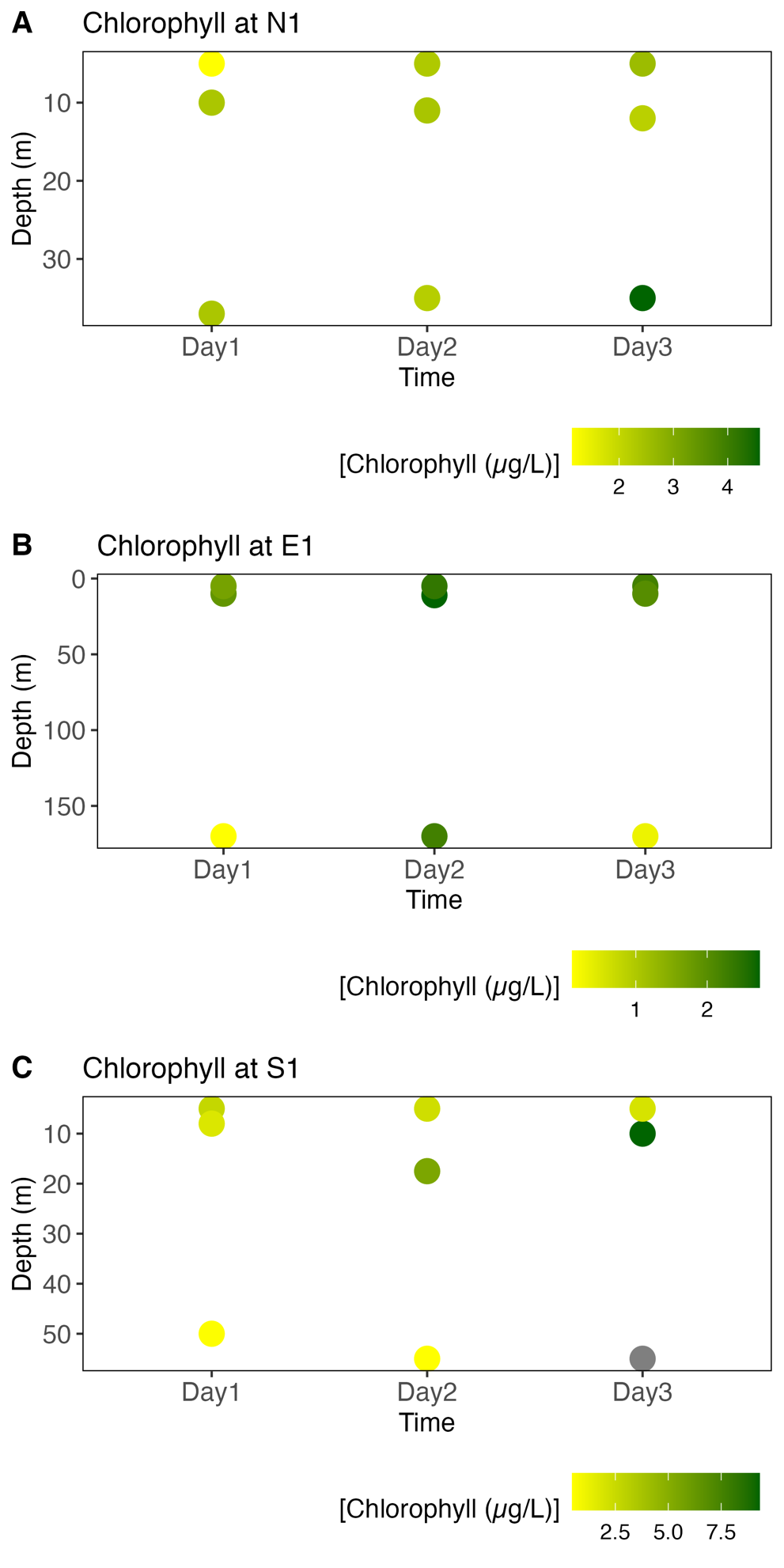

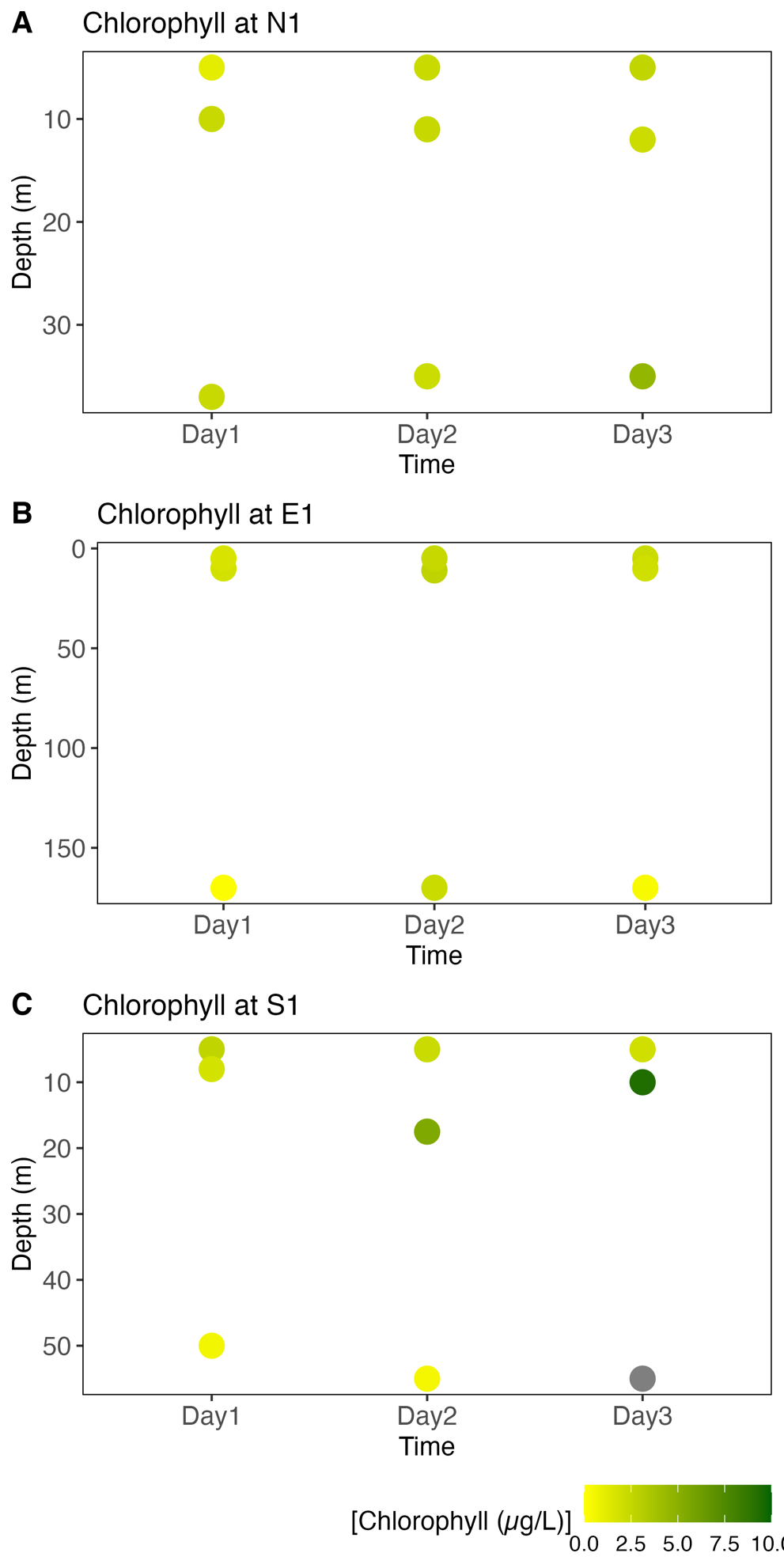

**Figure S2e:** Chlorophyll *a* from discrete measurements from Ship sampling at Eulerian sampling stations (N1, E1, and S1) over all three days. Plots show chlorophyll *a* on different scale bars (top) and on the same scale bar (bottom). Darker greens indicate a higher concentration of chlorophyll.

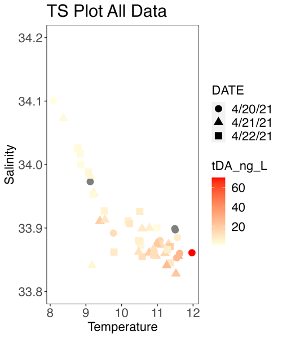

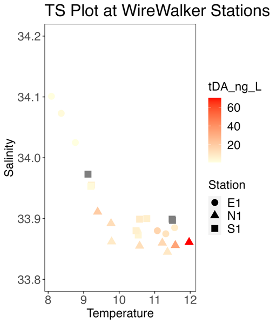

**Figure S2f.** Total Domoic acid (ng/L) in the context of temperature and salinity from all samples (left) and from Eulerian stations (right). Darker colors indicate more tDA, and the shapes indicate day (left) or station (right).

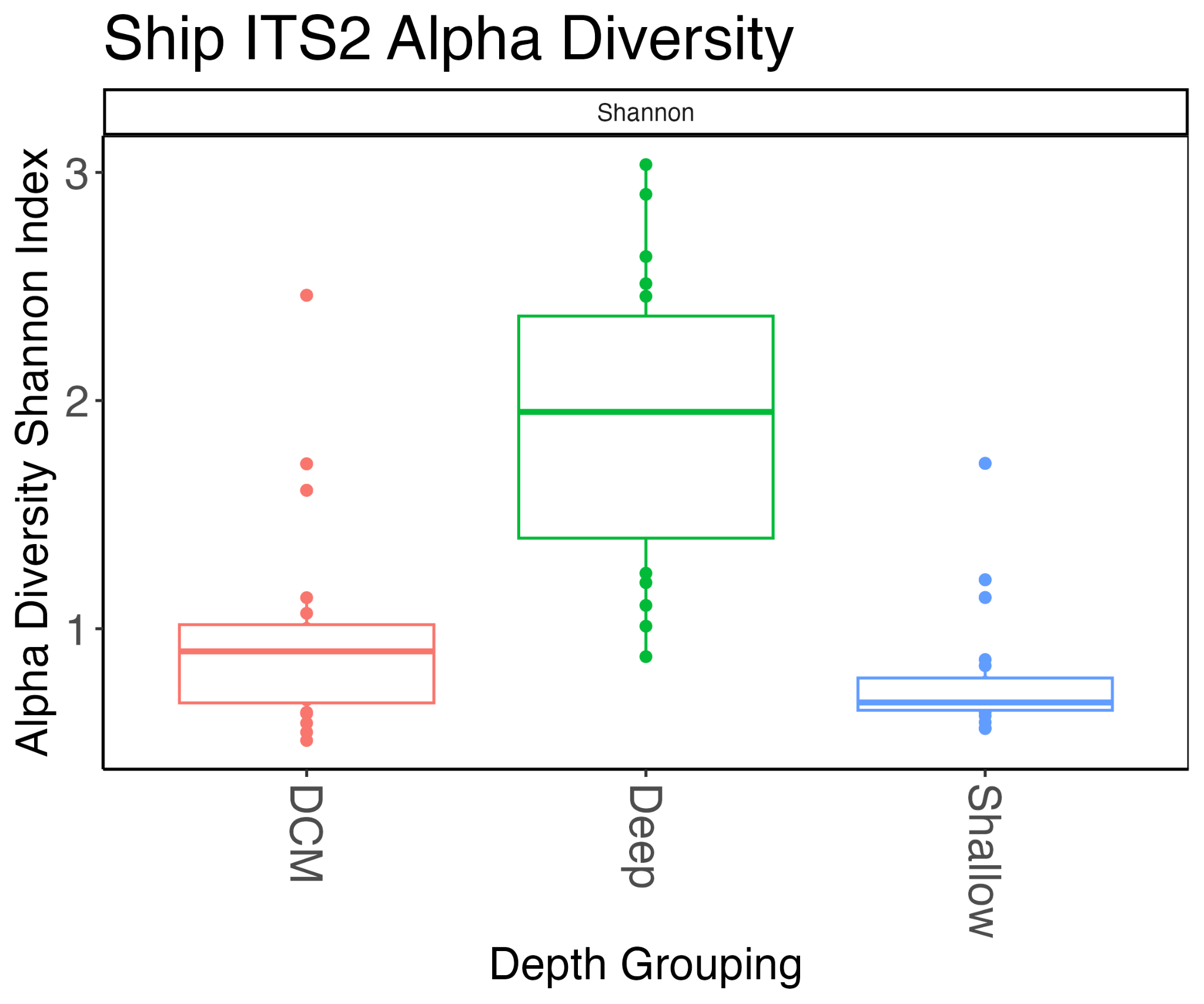

**Figure S3a:** ITS2 metabarcoding from ship samples filtered for *Pseudo-nitzschia* ITS2 only. Alpha diversity using the Shannon Index shows that the diversity is highest at depth relative to surface and SCM samples.

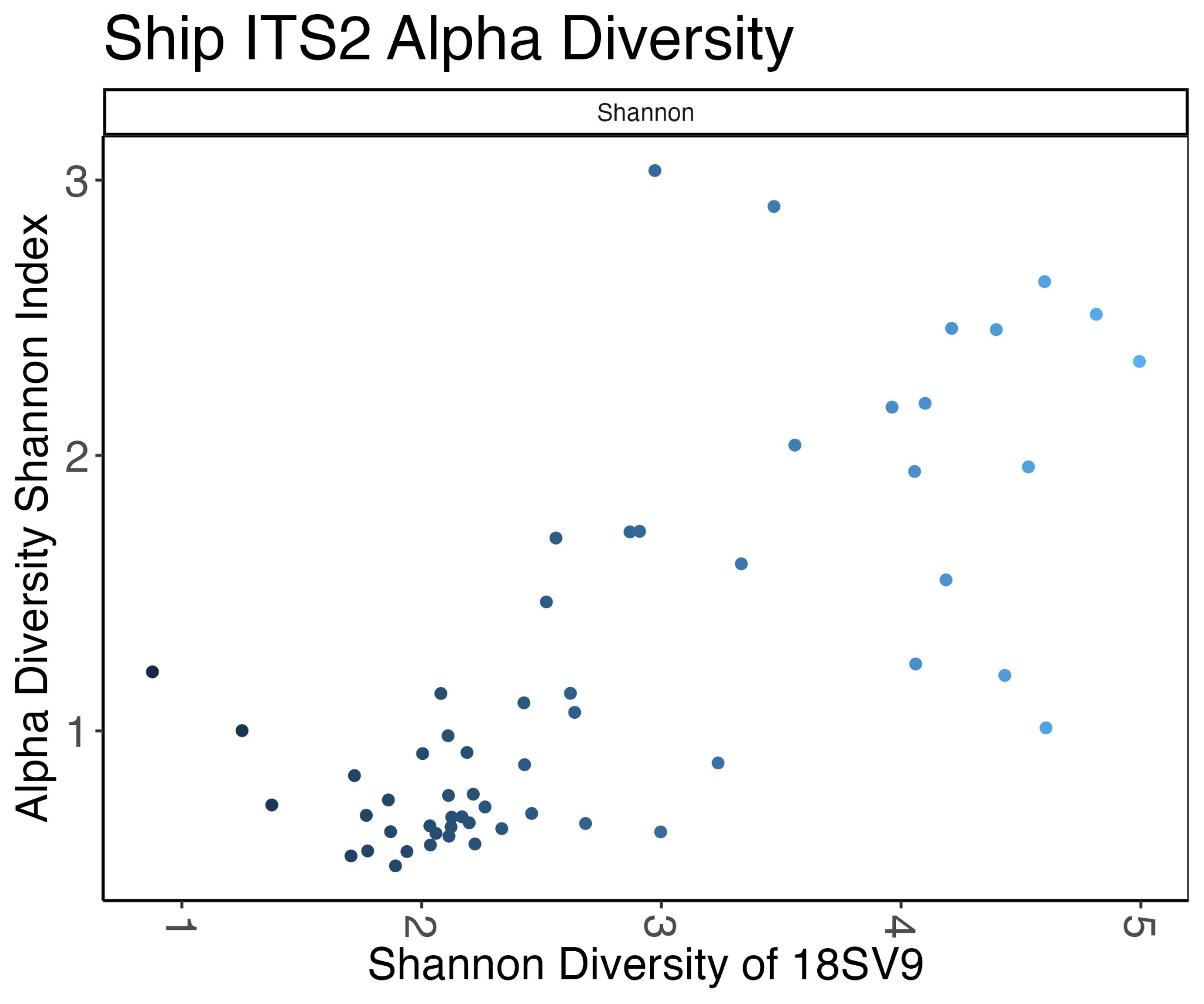

**Figure S3b:** ITS2 metabarcoding from ship samples filtered for *Pseudo-nitzschia* ITS2 only. Ship Sampling ITS2 metabarcoding Alpha diversity using the Shannon Index vs Ship Sampling 18SV9 Alpha diversity using the Shannon Index a linear relationship.

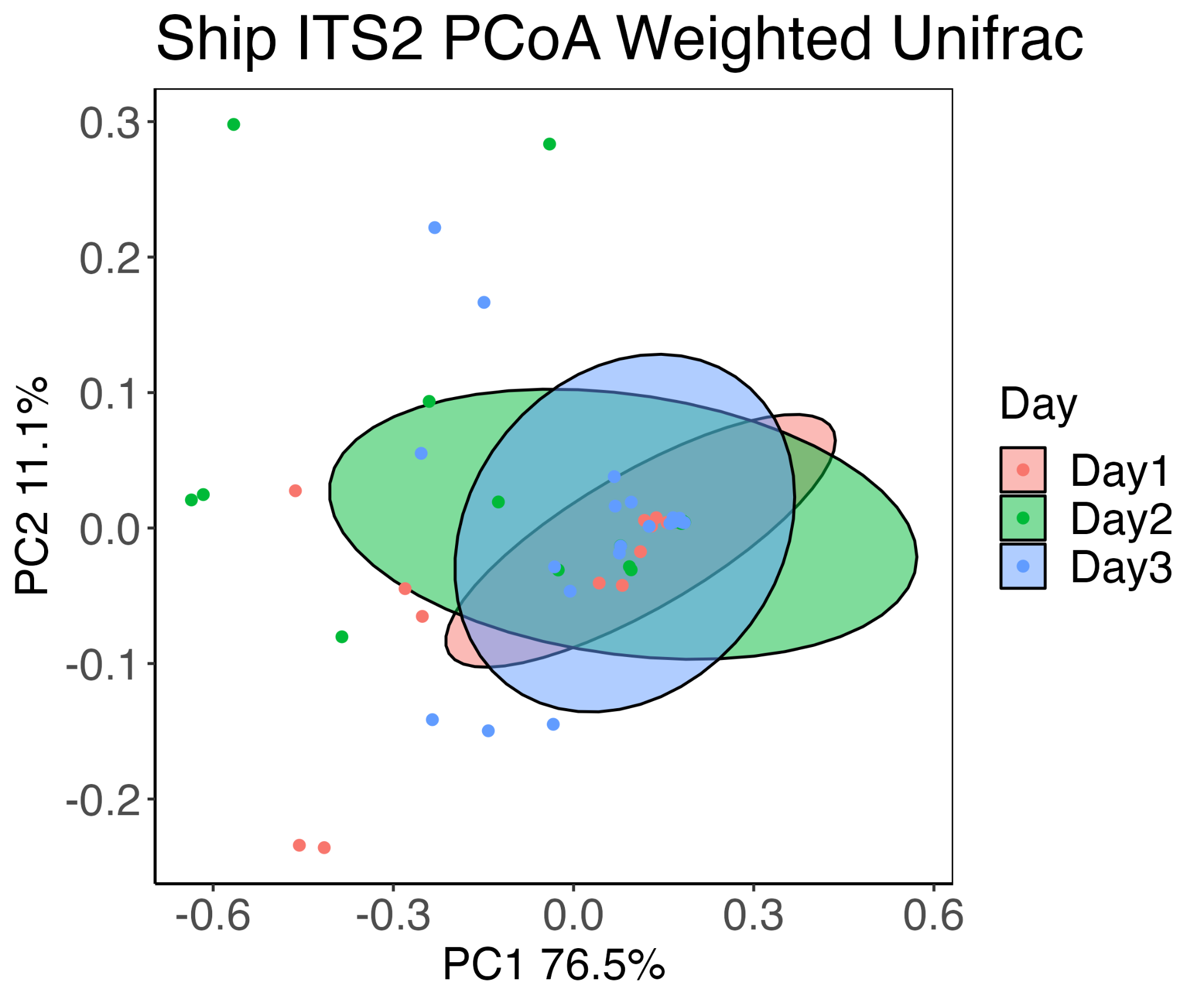

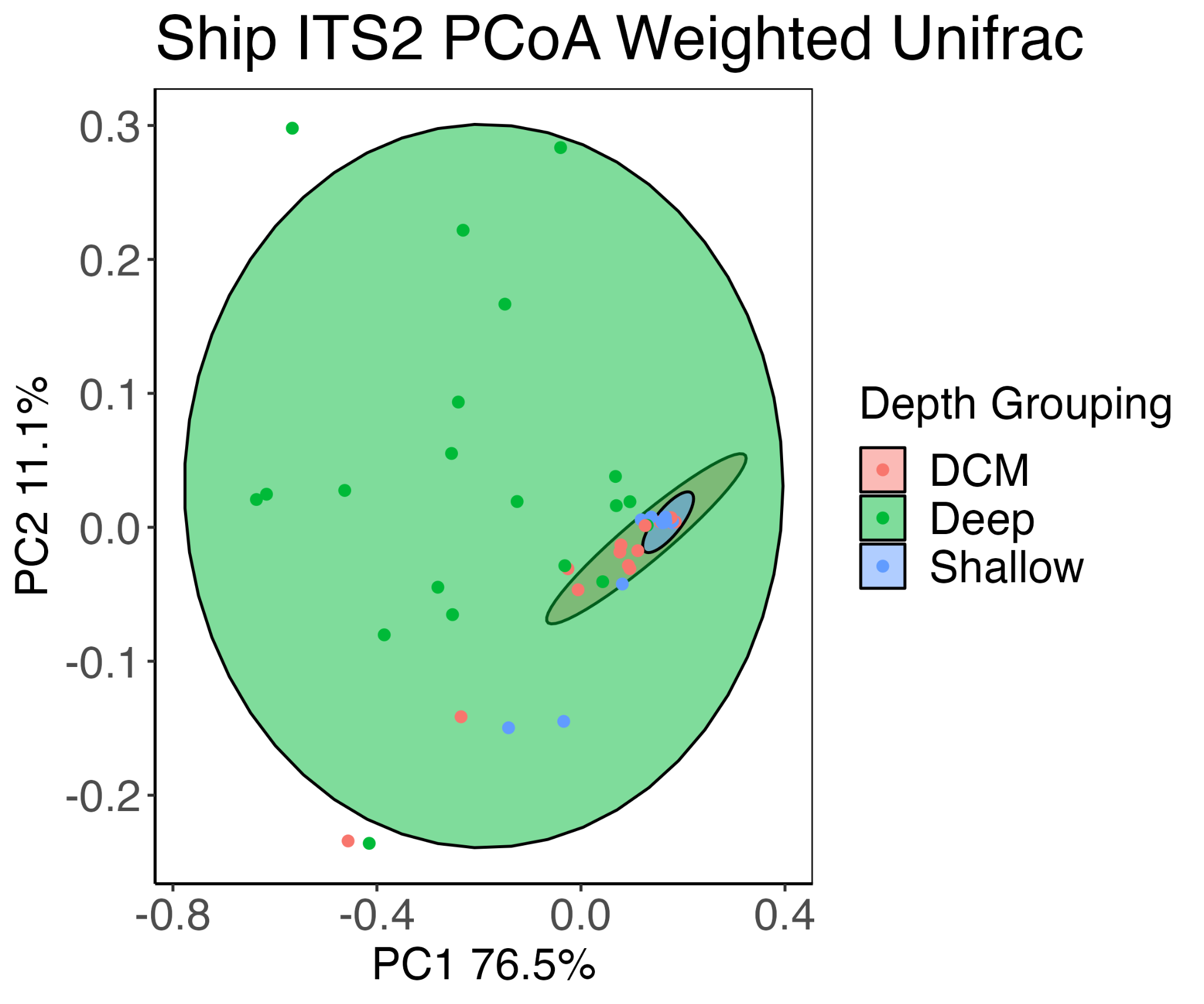

**Figure S3c:** ITS2 metabarcoding from ship samples filtered for *Pseudo-nitzschia* ITS2 only. Ship Sampling ITS2 metabarcoding Principal Component Analysis (PCA) does not reveal a difference in taxonomic composition over the three days (top). PCA shows that there are more unique ITS2 taxonomic compositions in deep samples, while the shallow and SCM samples are more similar to each other (bottom)

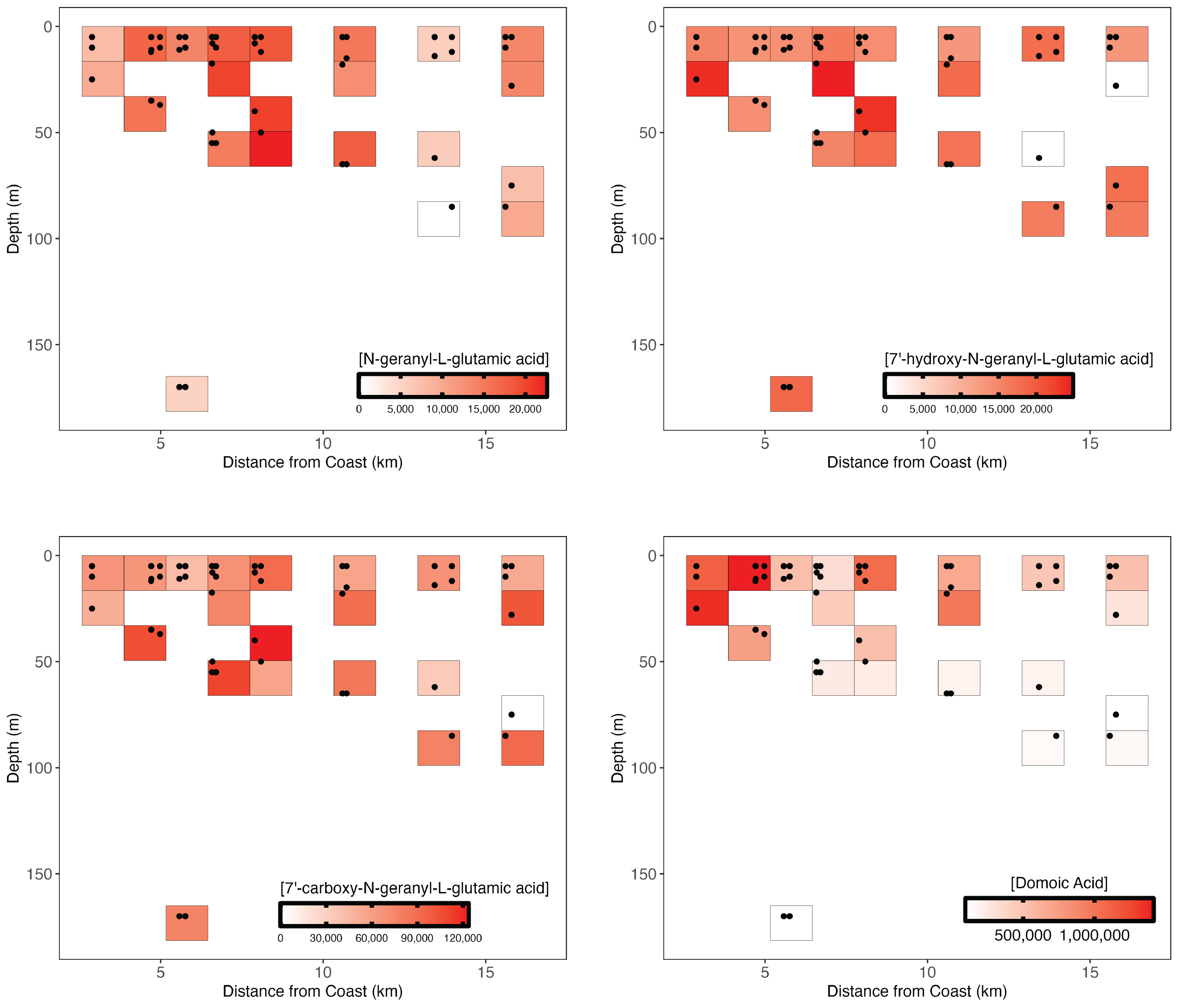

**Figure S3d.** The relative abundance of domoic acid biosynthetic intermediates spatially show spatial concurrence.

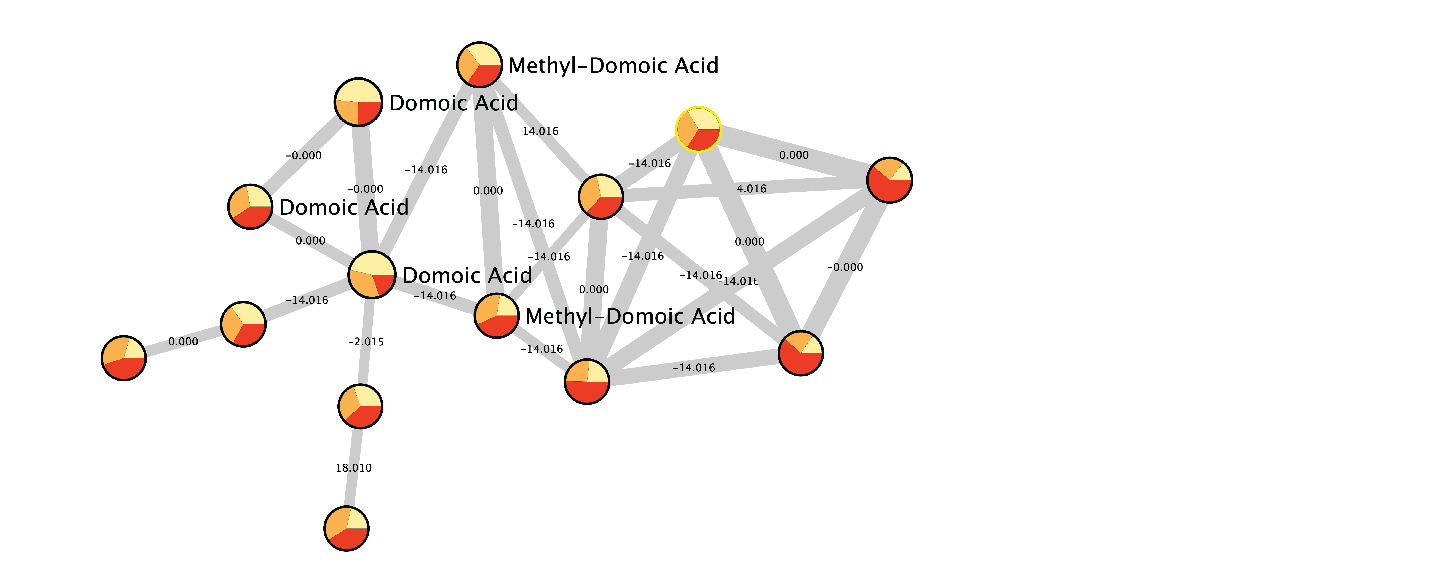

**Figure S3e:** Domoic acid and related molecules are observed in the environment using untargeted metabolomics. In this Global Natural Products Social Molecular Networking (GNPS) molecular network of all untargeted metabolomes, each node represents one compound and clustered nodes form networks of structurally related compounds. Thicker lines connecting nodes identify compounds with higher similarity. The pie charts forming each node are colored by abundance at each depth and sized by the intensity of the precursor ion (larger pies are more abundant in the dataset). The pies are colored by sampling day: yellow represents Day 1, orange Day 2, and red Day 3. The numbers connecting the nodes show a gain or loss of mass to charge (m/z) such as -14 indicating a loss of a methyl group.

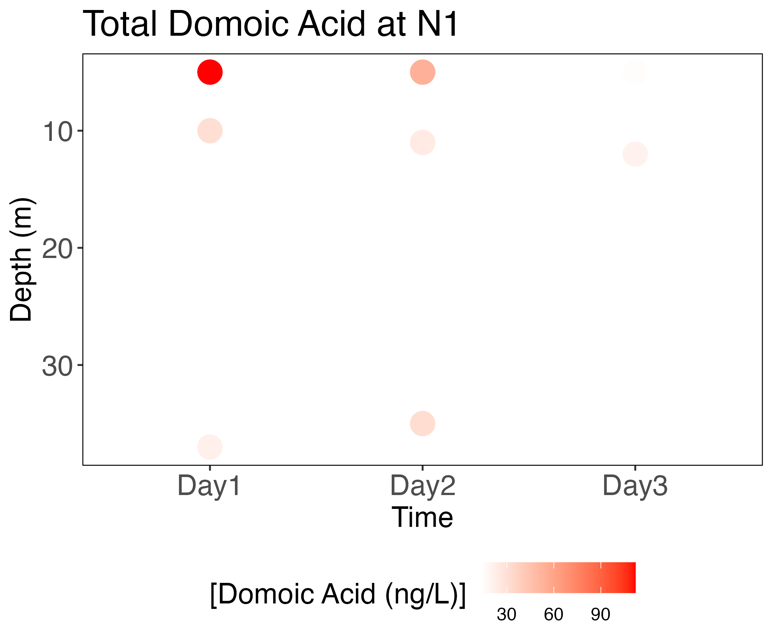

**Figure S4a.** Total DA at N1 is highest at the surface on the first day of sampling, and declined over time.

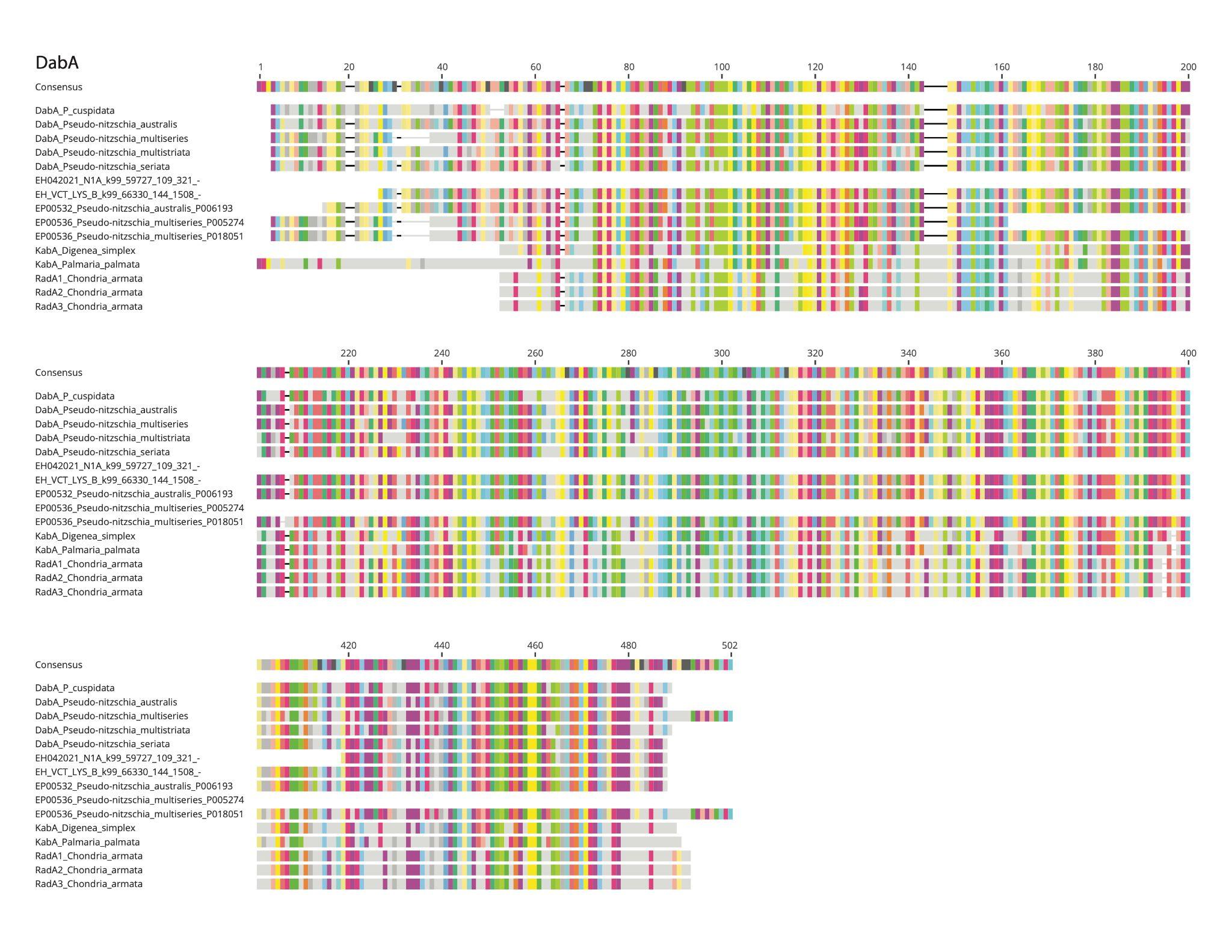

**Figure S5a:** MUSCLE5.1 alignments of reference Dab proteins with ORFs from the EcoHAB cruise, showing high conservation on the amino acid level. Only amino acids shared with the >25% consensus sequence are shown in color (1 of 3).

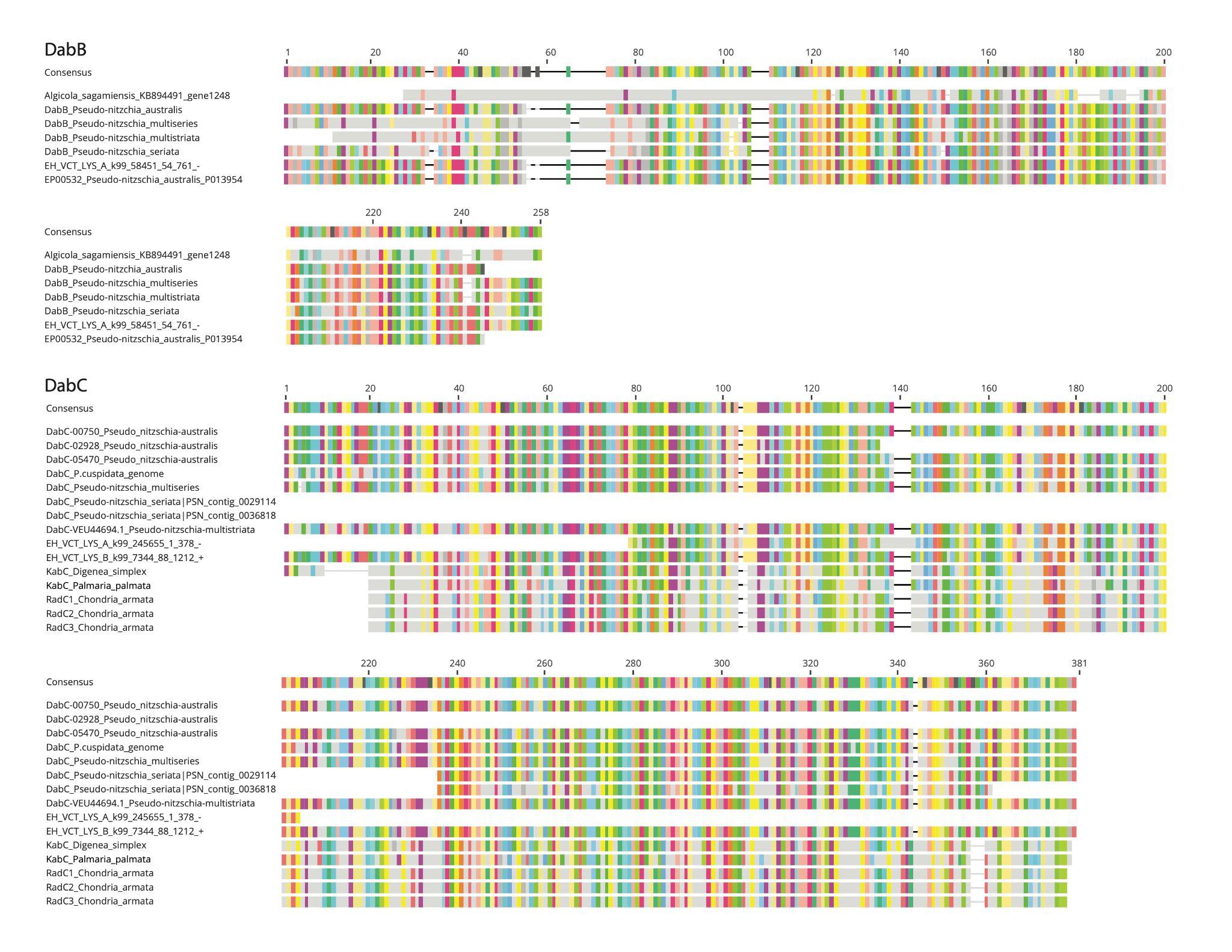
**Figure S5a:** MUSCLE5.1 alignments of reference Dab proteins with ORFs from the EcoHAB cruise, showing high conservation on the amino acid level. Only amino acids shared with the >25% consensus sequence are shown in color (2 of 3).

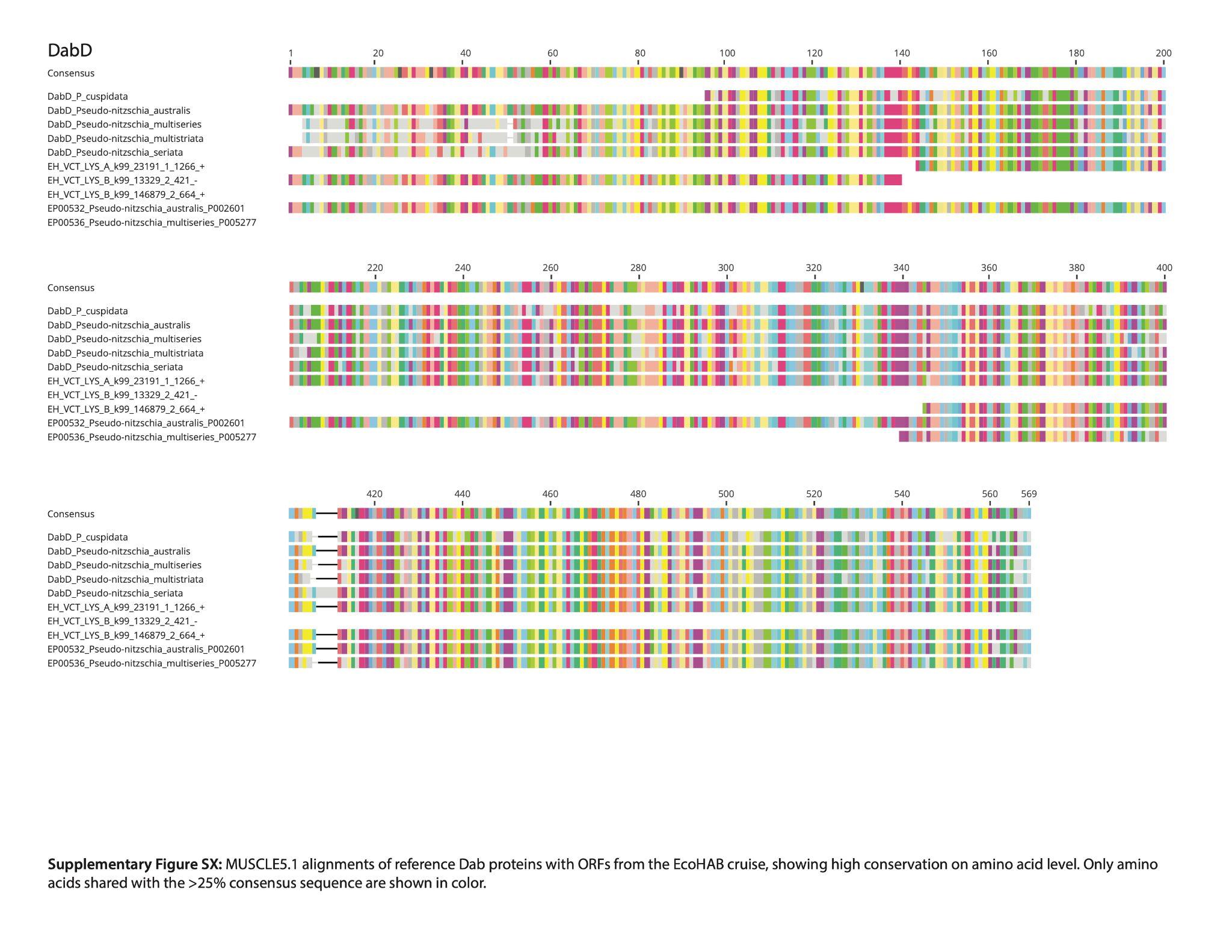
**Figure S5a:** MUSCLE5.1 alignments of reference Dab proteins with ORFs from the EcoHAB cruise, showing high conservation on the amino acid level. Only amino acids shared with the >25% consensus sequence are shown in color (3 of 3).

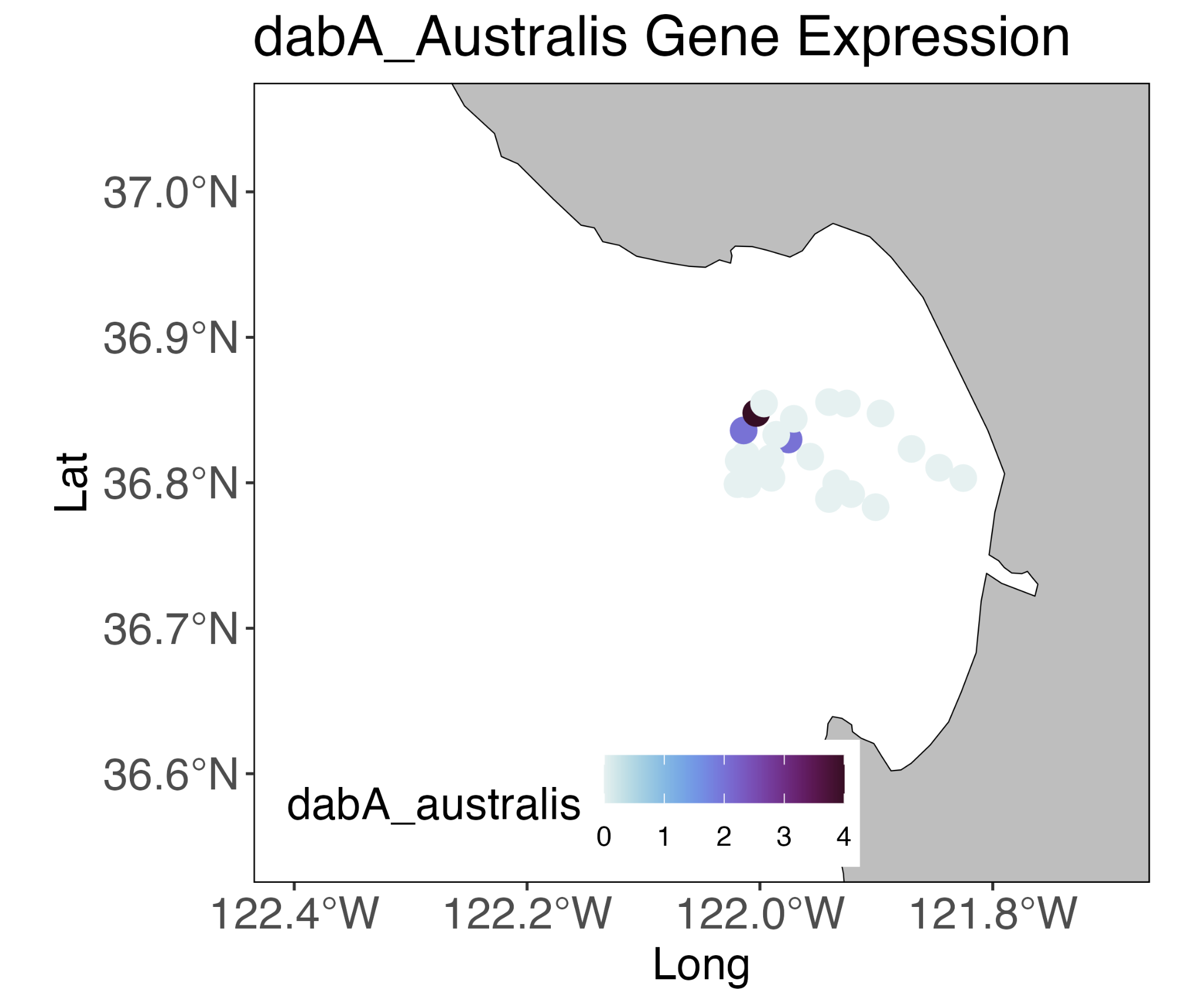

**Figure S5b.** *dabA* gene expression *P. australis* from samples from the 3G-ESP/LRAUV.

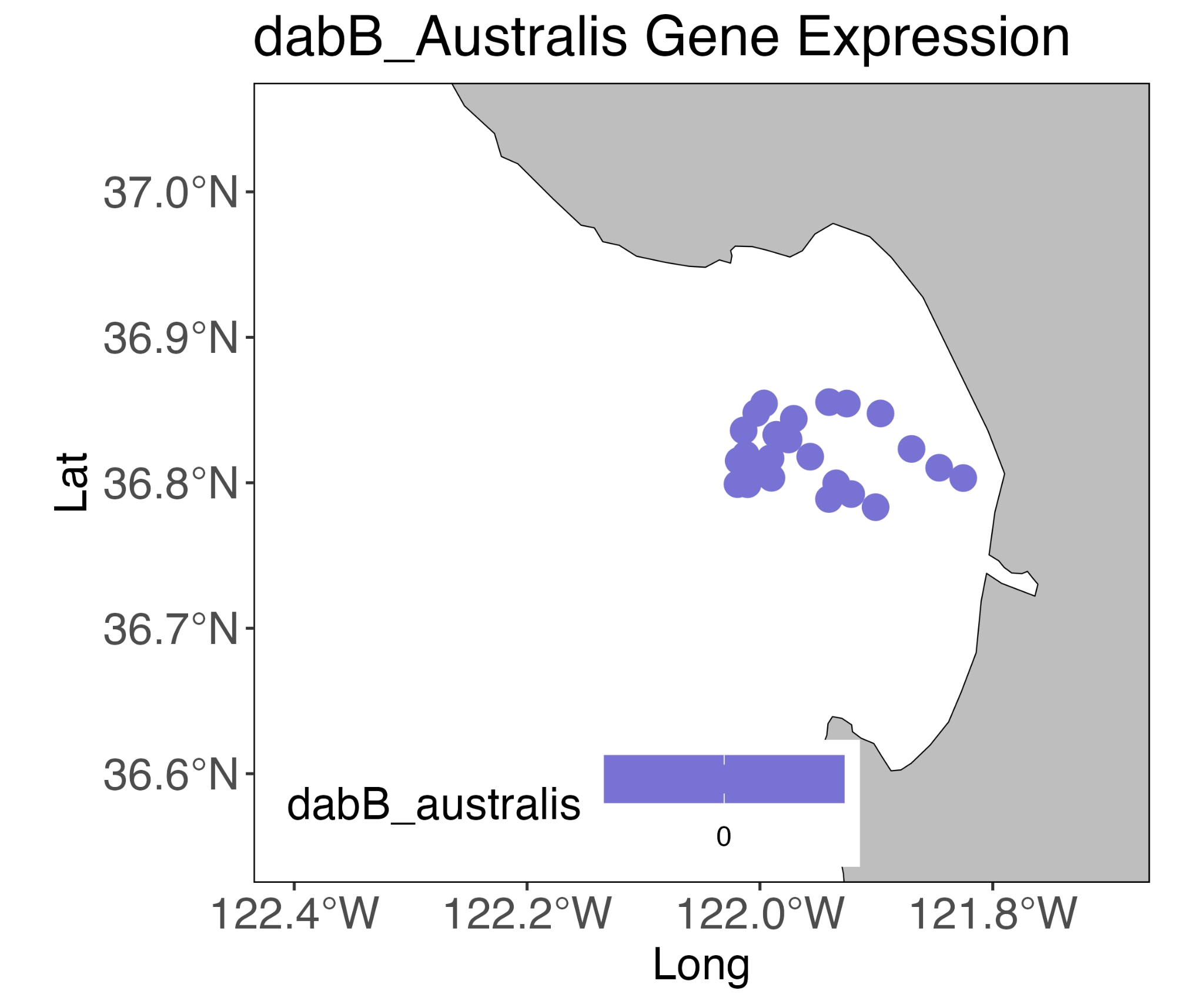

**Figure S5c.** *dabB* gene expression *P. australis* from samples from the 3G-ESP/LRAUV. No gene expression was detected.

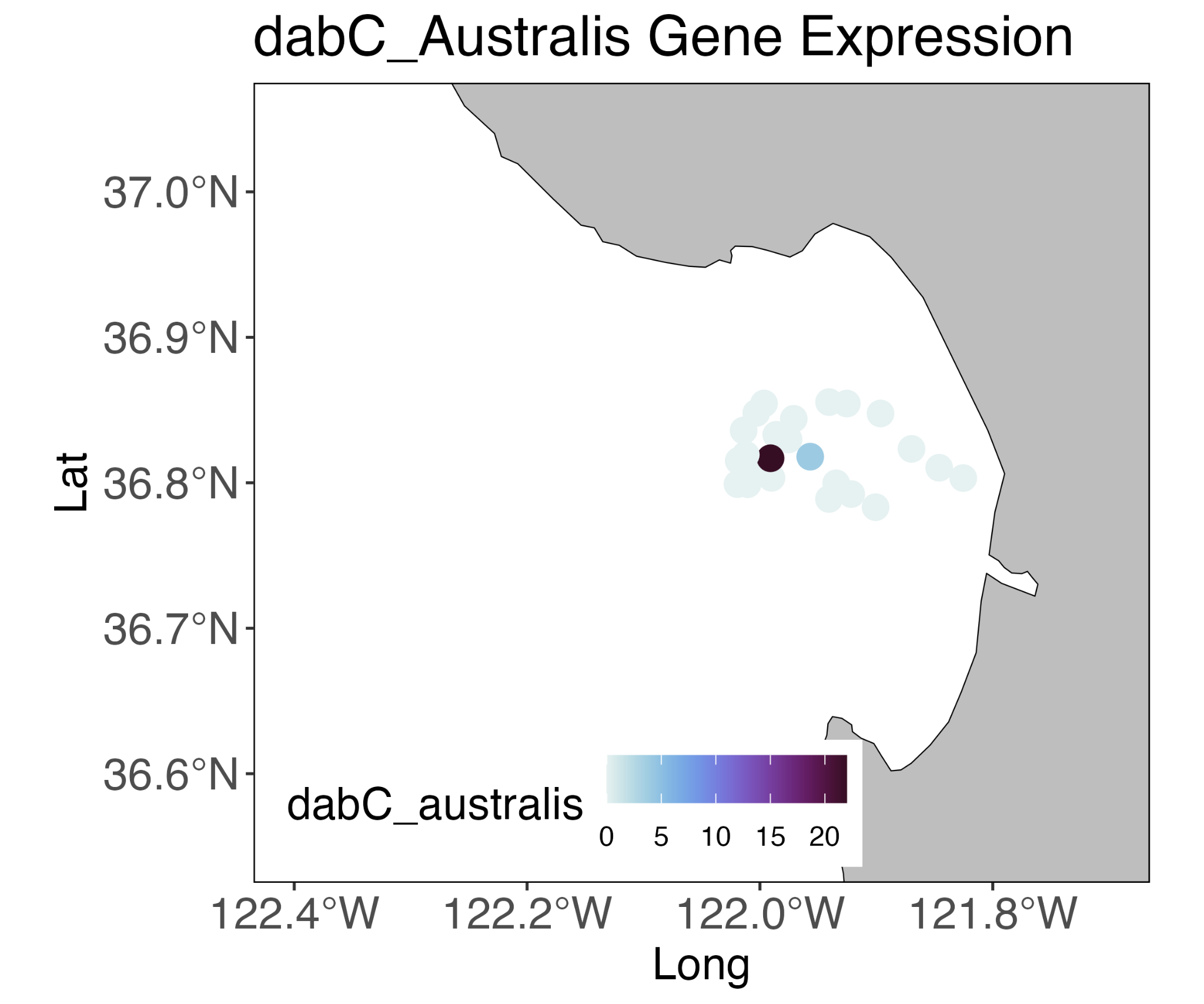

**Figure S5d.** *dabC* gene expression *P. australis* from samples from the 3G-ESP/LRAUV.

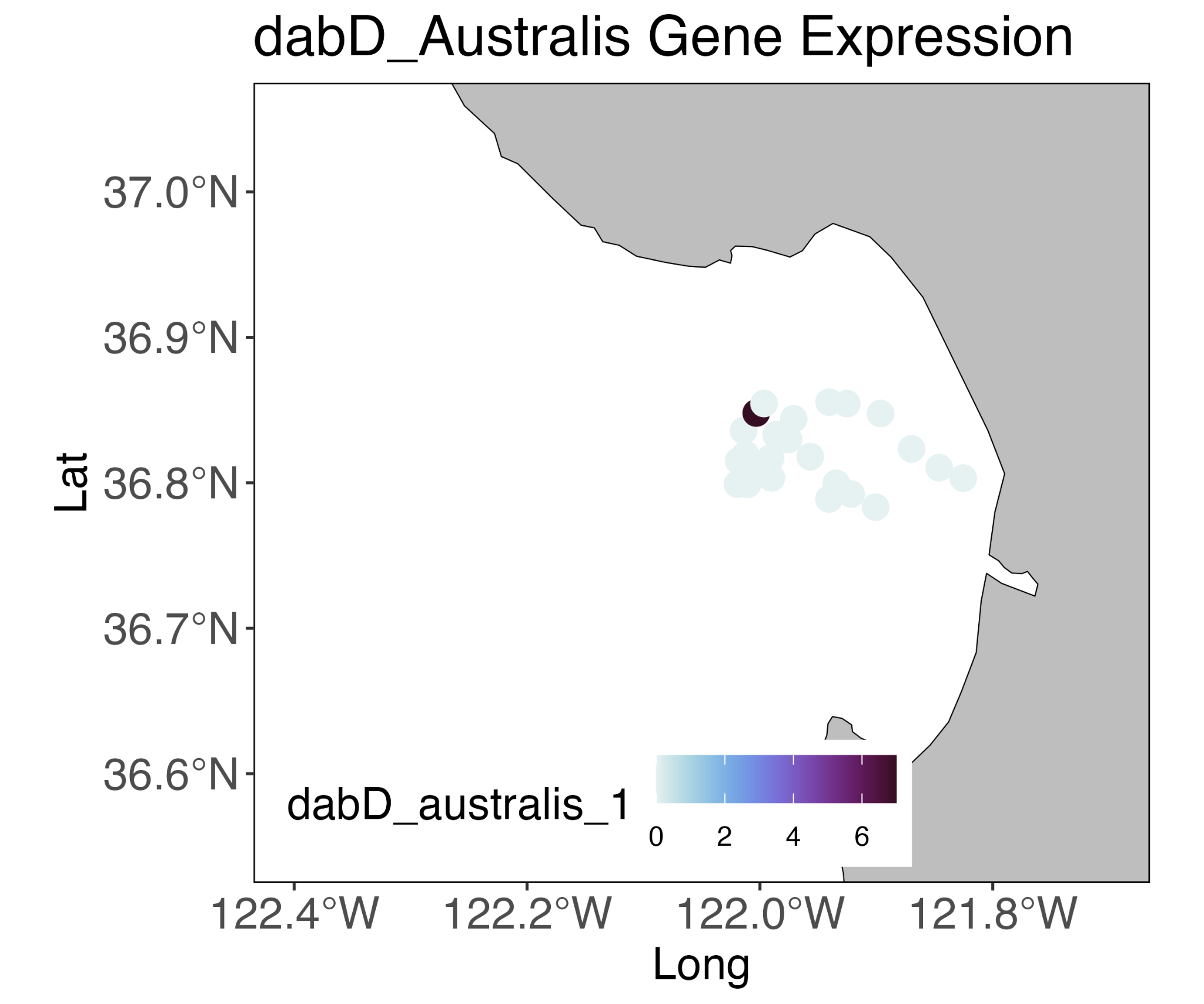

**Figure S5e.** *dabD* gene expression *P. australis* from samples from the 3G-ESP/LRAUV.

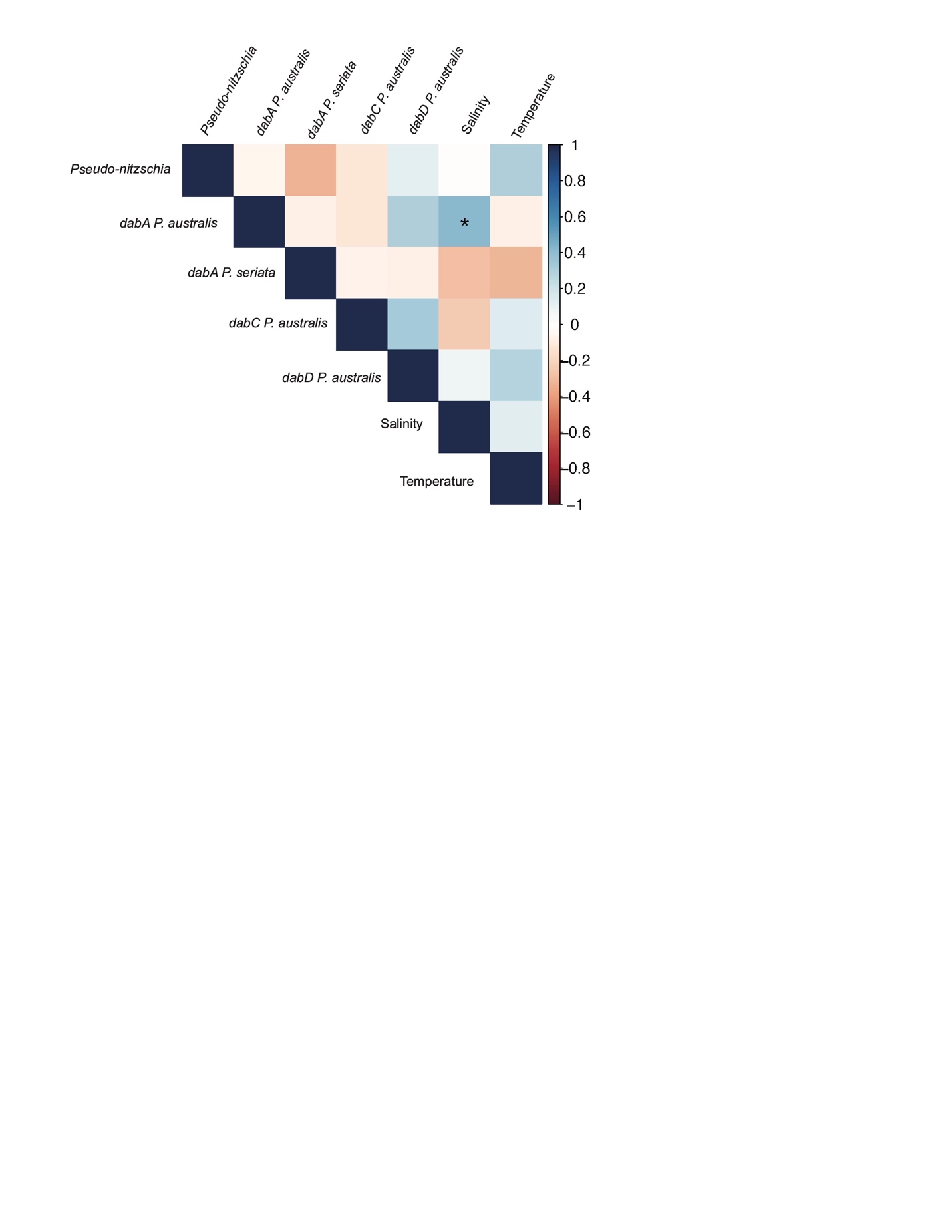

**Figure S5f.** Spearman rank correlations identified relationships between environmental parameters and *dab* gene expression from AUV samples, where values closer to 1 (dark blue) are positive correlations, and values closer to −1 (dark red) are negative correlations, and an asterisk (*) represents significant correlations. The data shows that expressions of *dabA, dabC,* and *dabD* are linked in the environment.

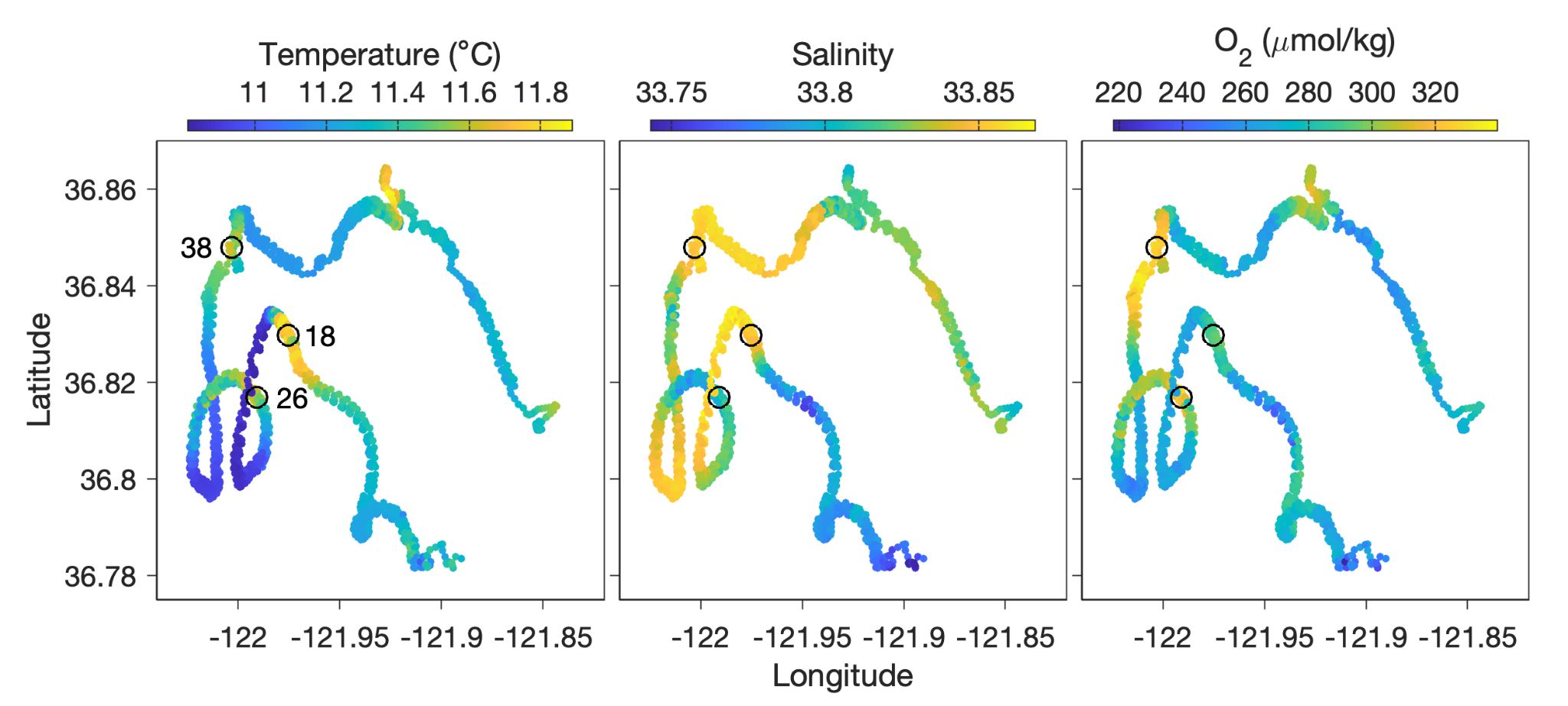

**Figure S5g:** Measurements of temperature, salinity and oxygen from the surface wave glider that followed the LRAUV. Select samples collected by the 3G-ESP are circled.

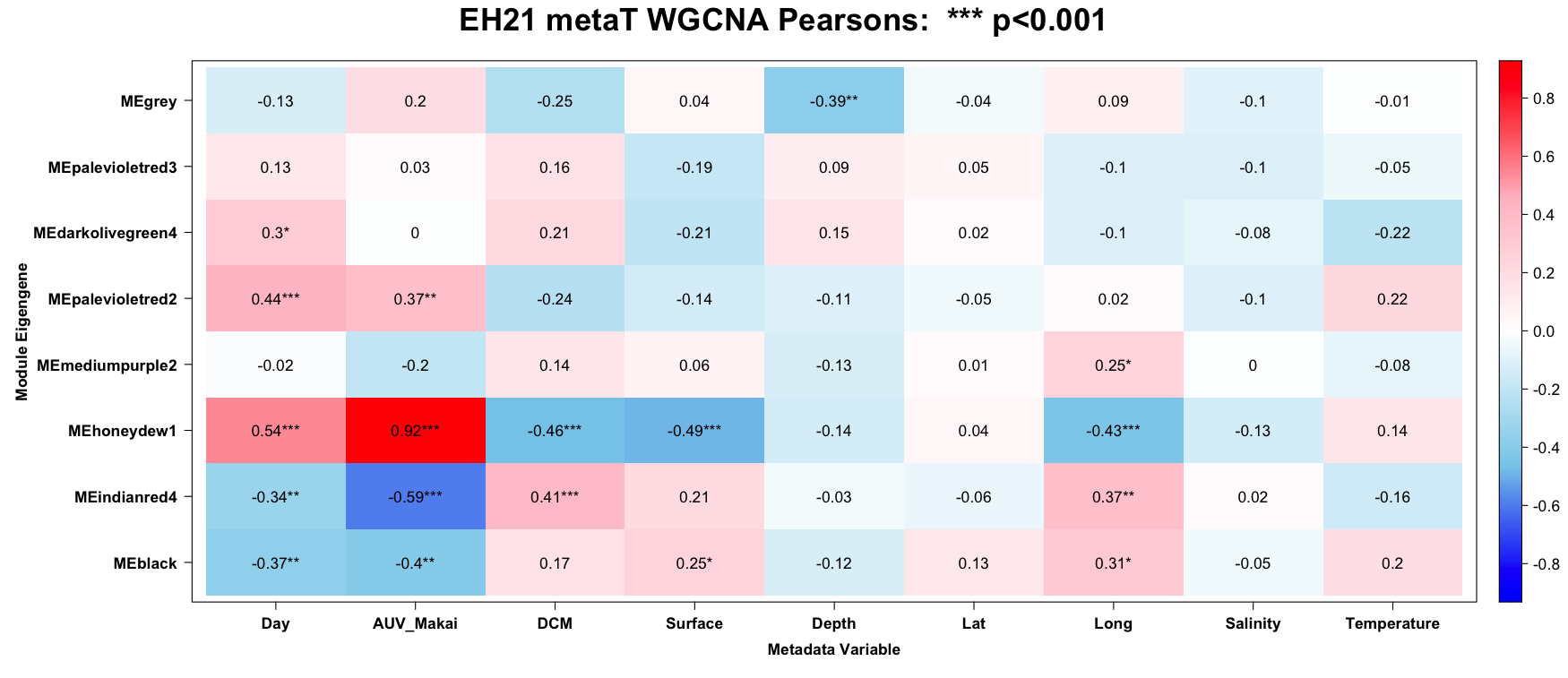

**Figure S5h:** Weighted Gene Correlation Network Analysis (WGCNA) was used to define seven modules of co-expressed ORFs. Associations between modules and metadata variables were analyzed using Pearson’s correlation. Highly significant associations indicated by *** (p<0.001) include a strong positive association between the MEhoneydew1 module and samples from the AUV. The MEhoneydew1 module also has a significant negative association with samples from the SCM and the Surface, and with increased days of the experiment.

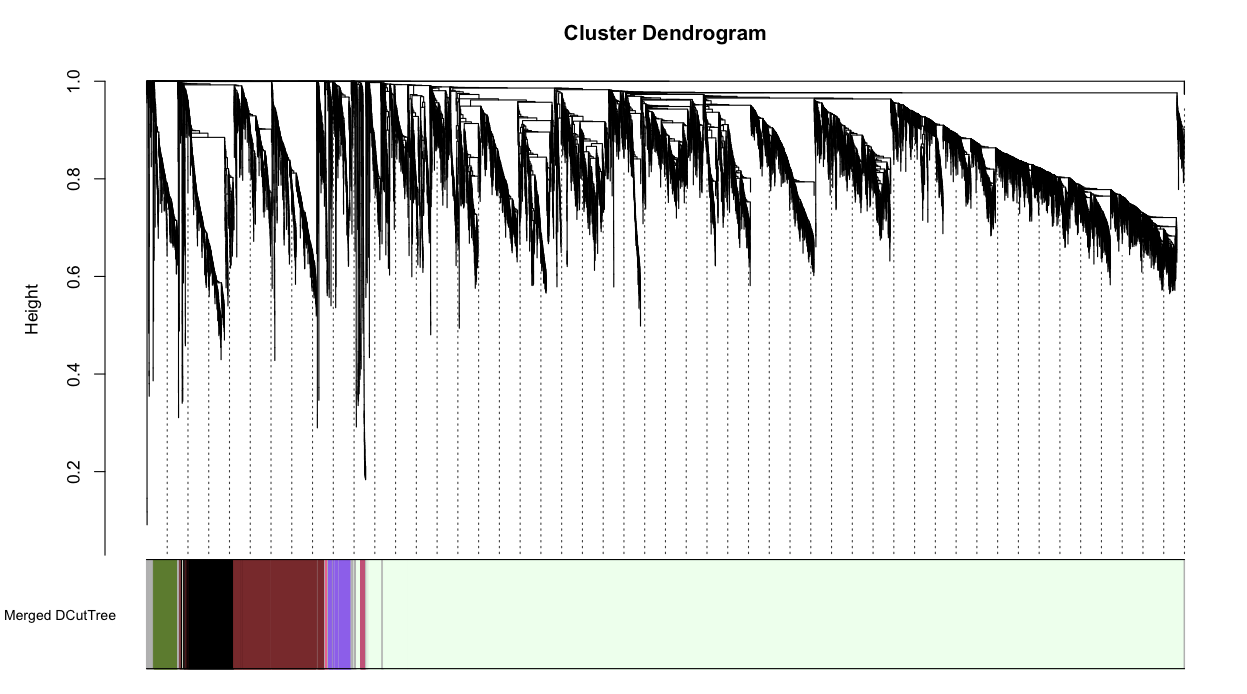
**Figure S5i:** Weighted Gene Correlation Network Analysis (WGCNA) was used to define seven modules of co-expressed ORFs. ORFs are plotted on a dendrogram showing similarity of expression.

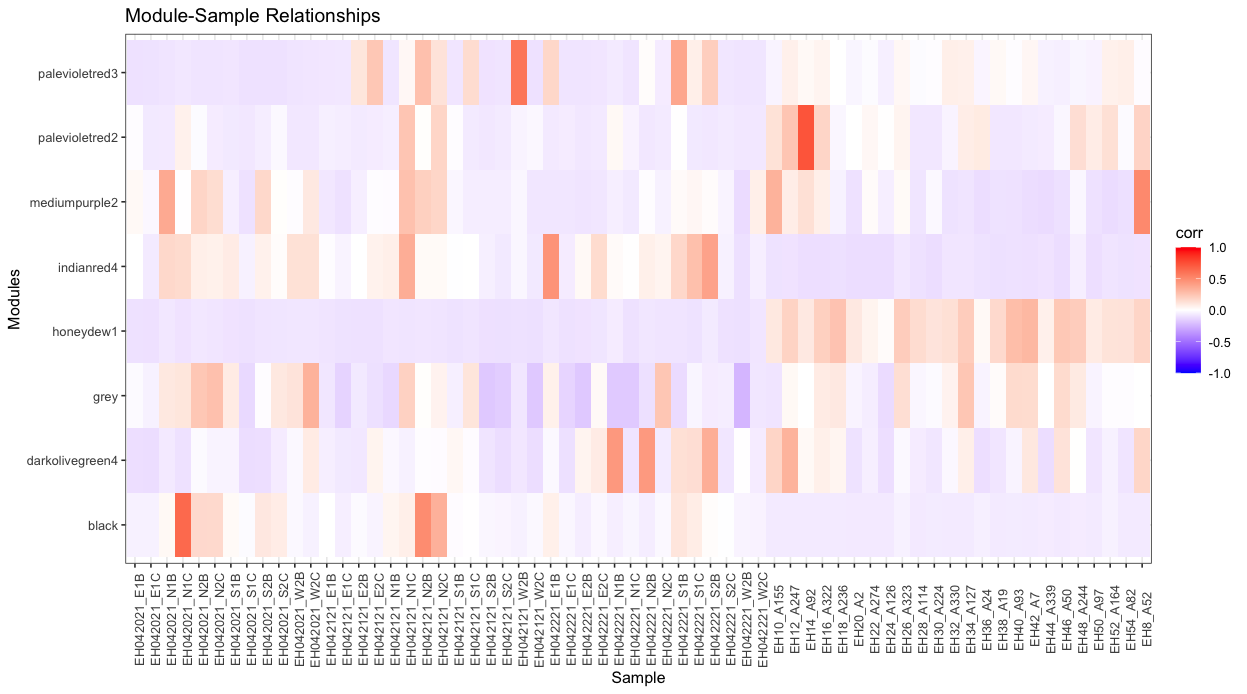

**Figure S5j:** Weighted Gene Correlation Network Analysis (WGCNA) was used to define seven modules of co-expressed ORFs. This heat map shows the strength of positive (red) and negative (white) correlations between samples and each of the seven modules.

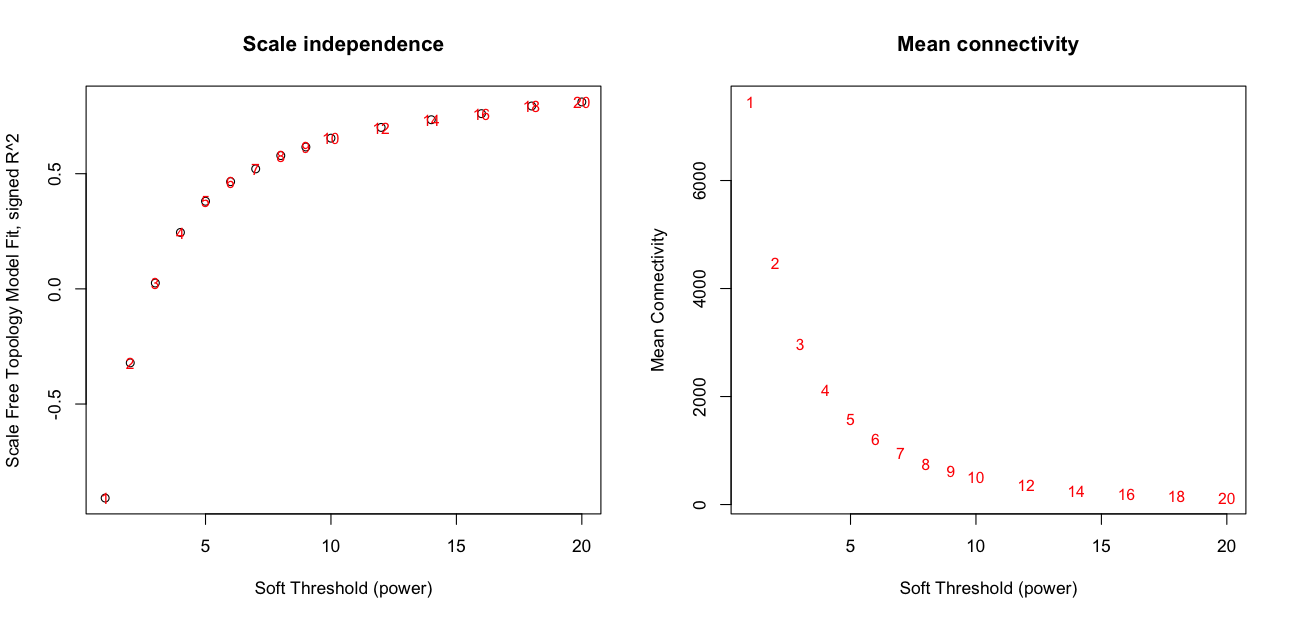

**Figure S5k:** Weighted Gene Correlation Network Analysis (WGCNA) was used to define seven modules of co-expressed ORFs. Scale independence and mean connectivity plots from WGCNA analysis.

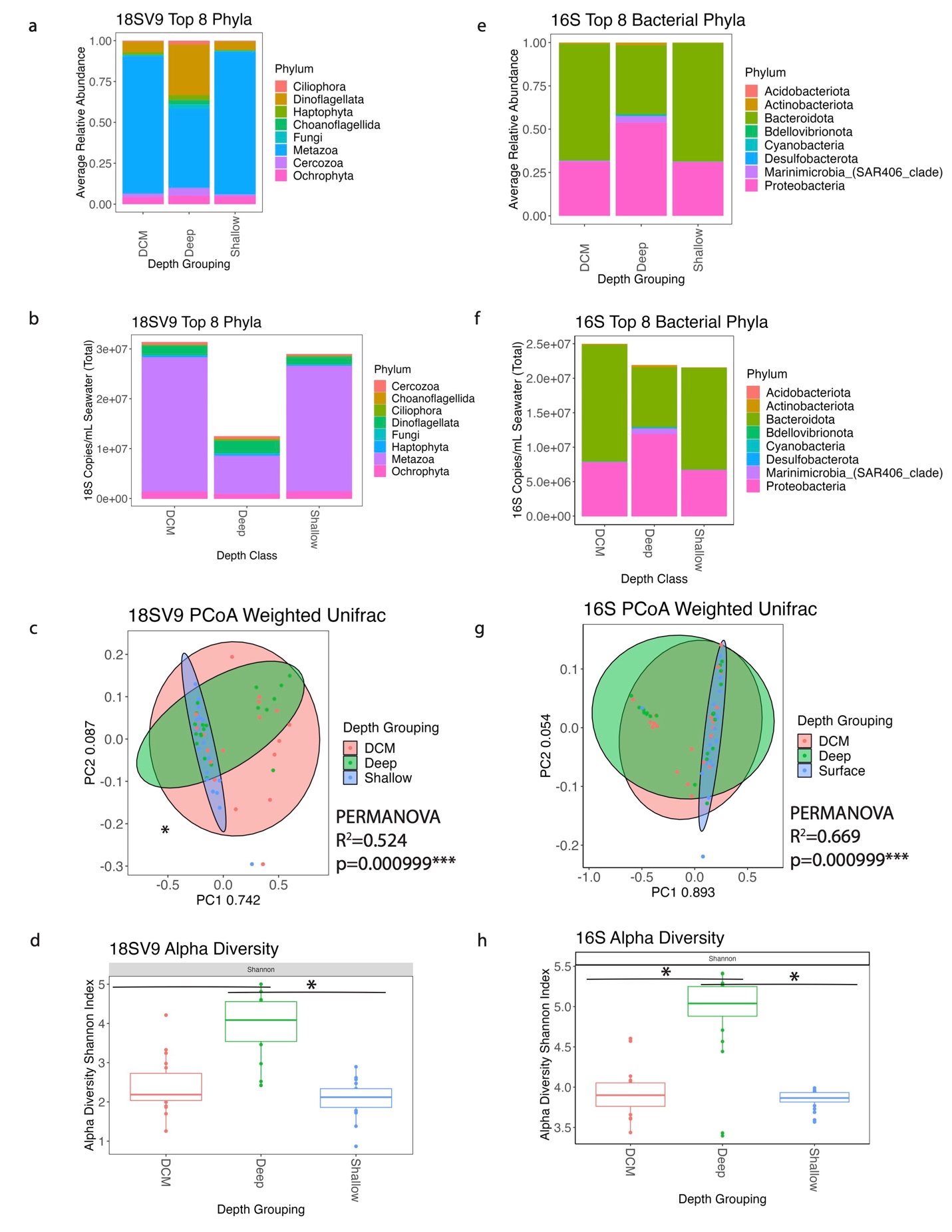

**Figure S6a-h.** Overview of 18SV9 and 16S Non-Plastid (“16S”) Metabarcoding Data from Ship Sampling. Depth classes are: “Shallow/Surface” (approximately 5 m from surface), “SCM” (variable), and “Deep” (approximately 5 m from seafloor, variable). 8 most abundant phyla in 18SV9 (a) and 16S bacteria (b). 18SV9 gene copies per mL sea water (c) and 16S gene copies per mL of seawater (d). Principle component analysis of 18SV9 (e) and 16S (f). Alpha diversity using the Shannon Index of 18SV9 (g) and 16S (h) with an asterisk (*) indicating significant difference by pairwise Wilcoxon Rank Sum Tests (p<0.05).

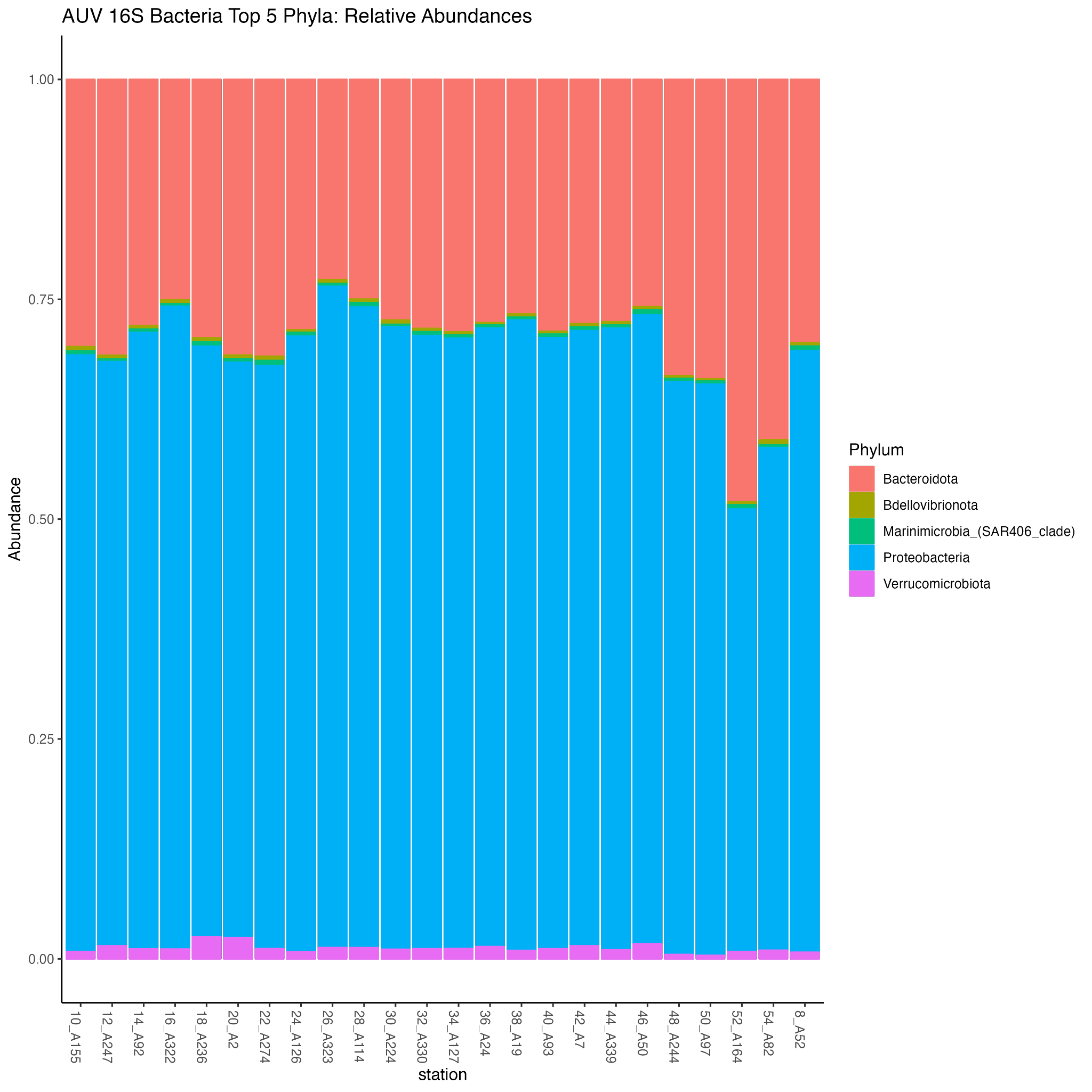

**Figure S7a:** 16S metabarcoding from samples collected by the 3G-ESP. Relative abundance of the five most abundant bacterial phyla reveals synteny in the bacterial community over time.

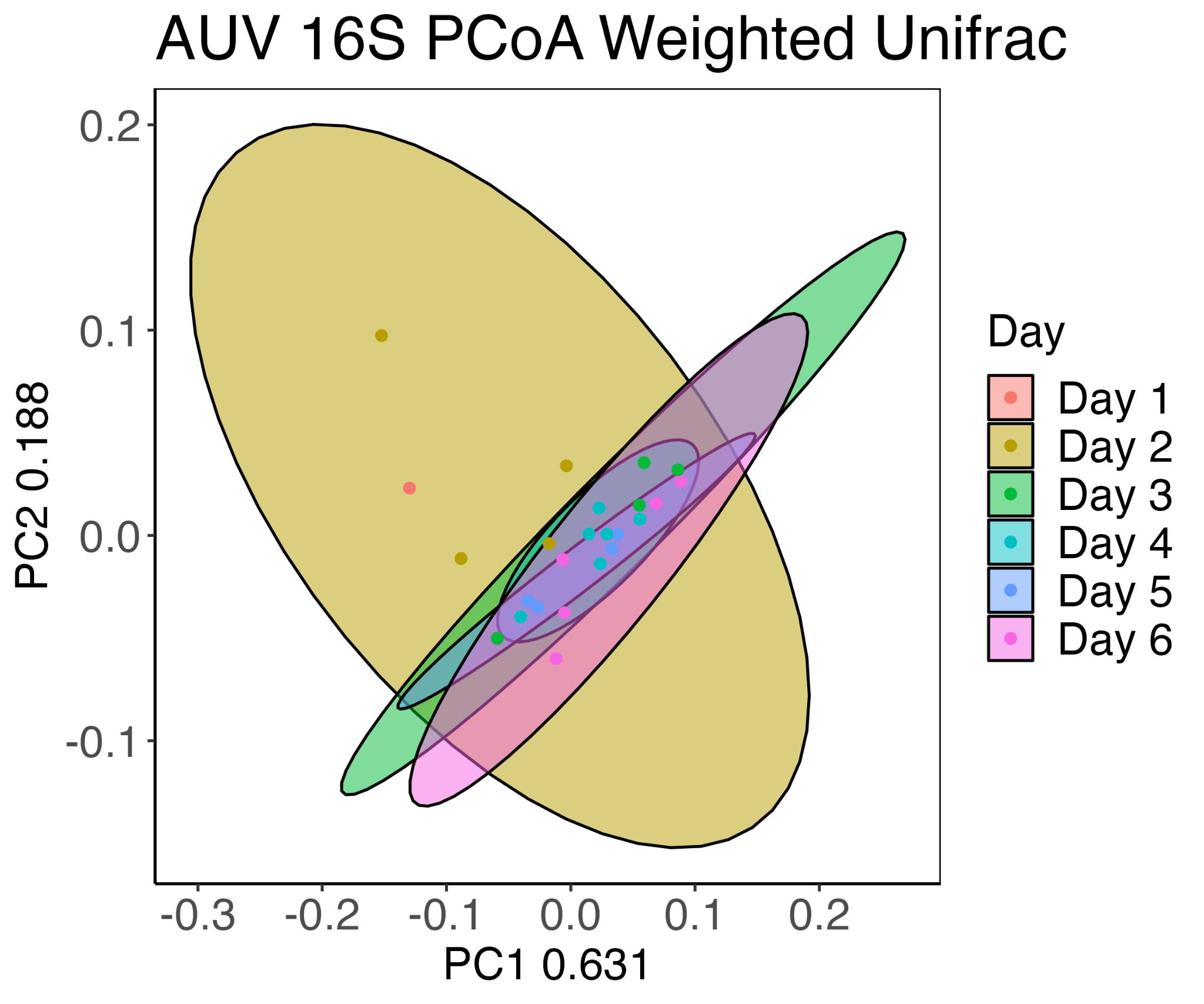

**Figure S7b:** 16S metabarcoding from samples collected by the 3G-ESP. Principal Component Analysis reveals synteny in the prokaryotic community over time.

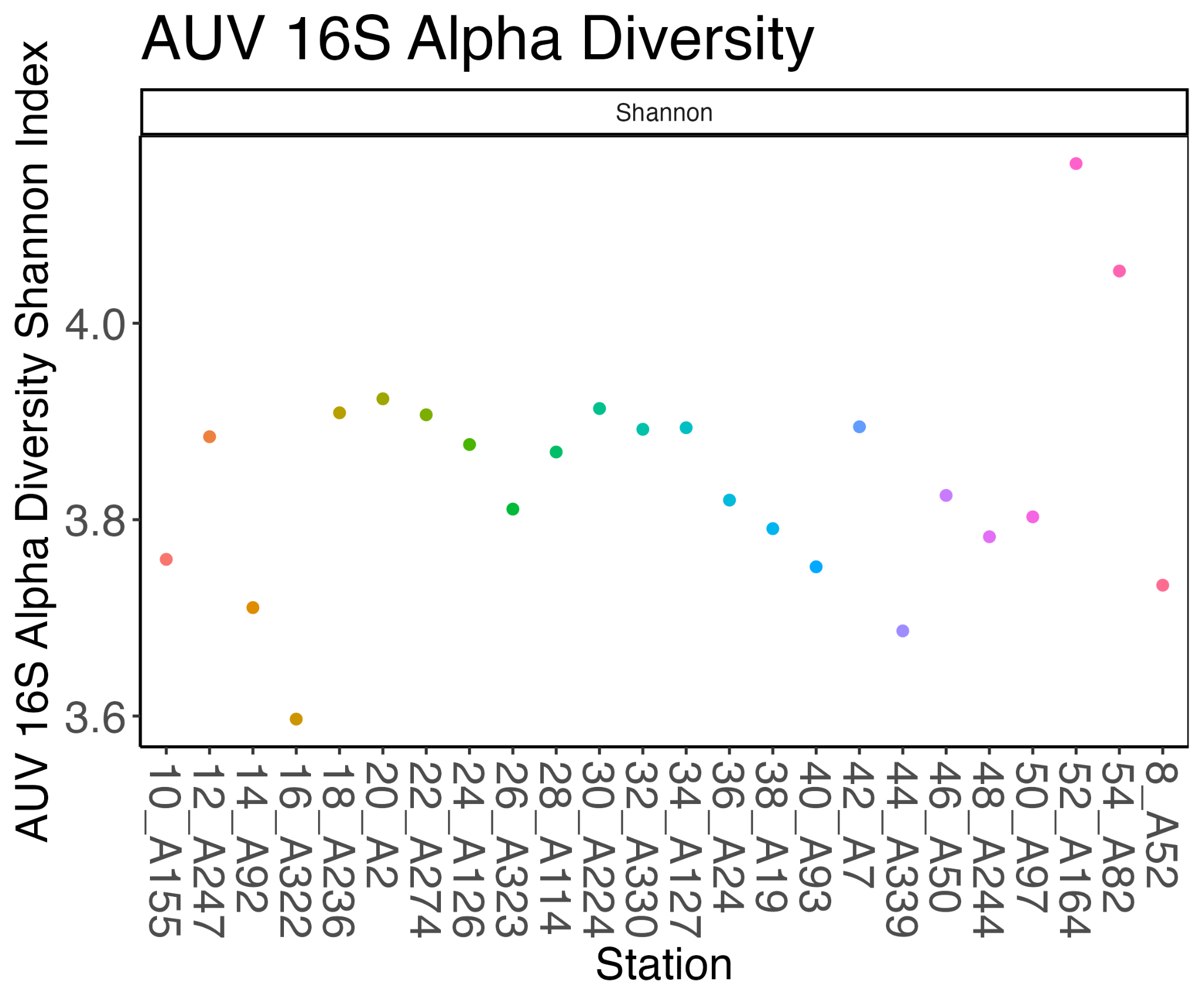

**Figure S7c:** 16S metabarcoding from samples collected by the 3G-ESP. Alpha diversity reveals synteny in the prokaryotic community over time.

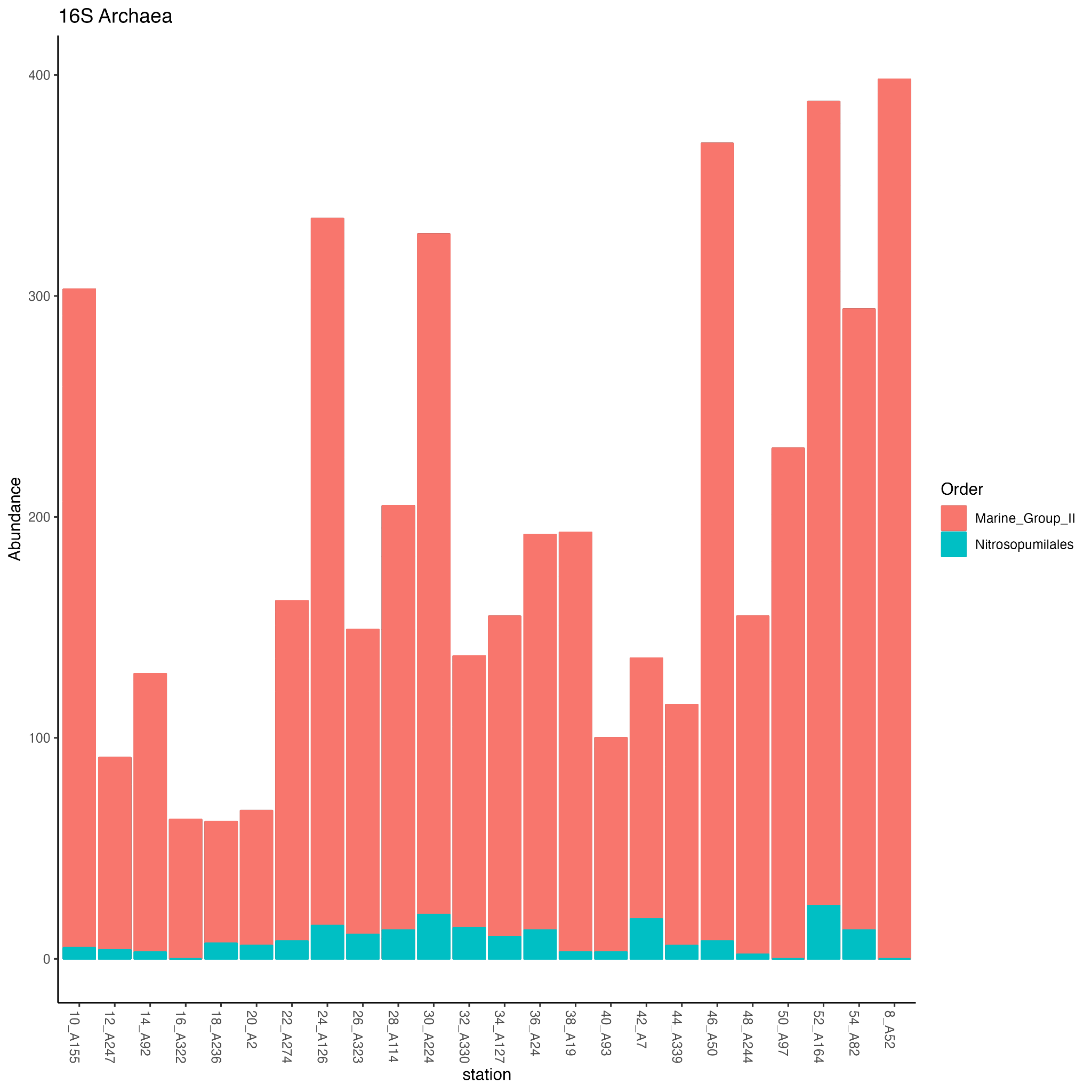

**Figure S7d:** 16S metabarcoding from samples collected by the 3G-ESP. Read count abundance of archaeal orders reveals synteny in the archaeal community over time.

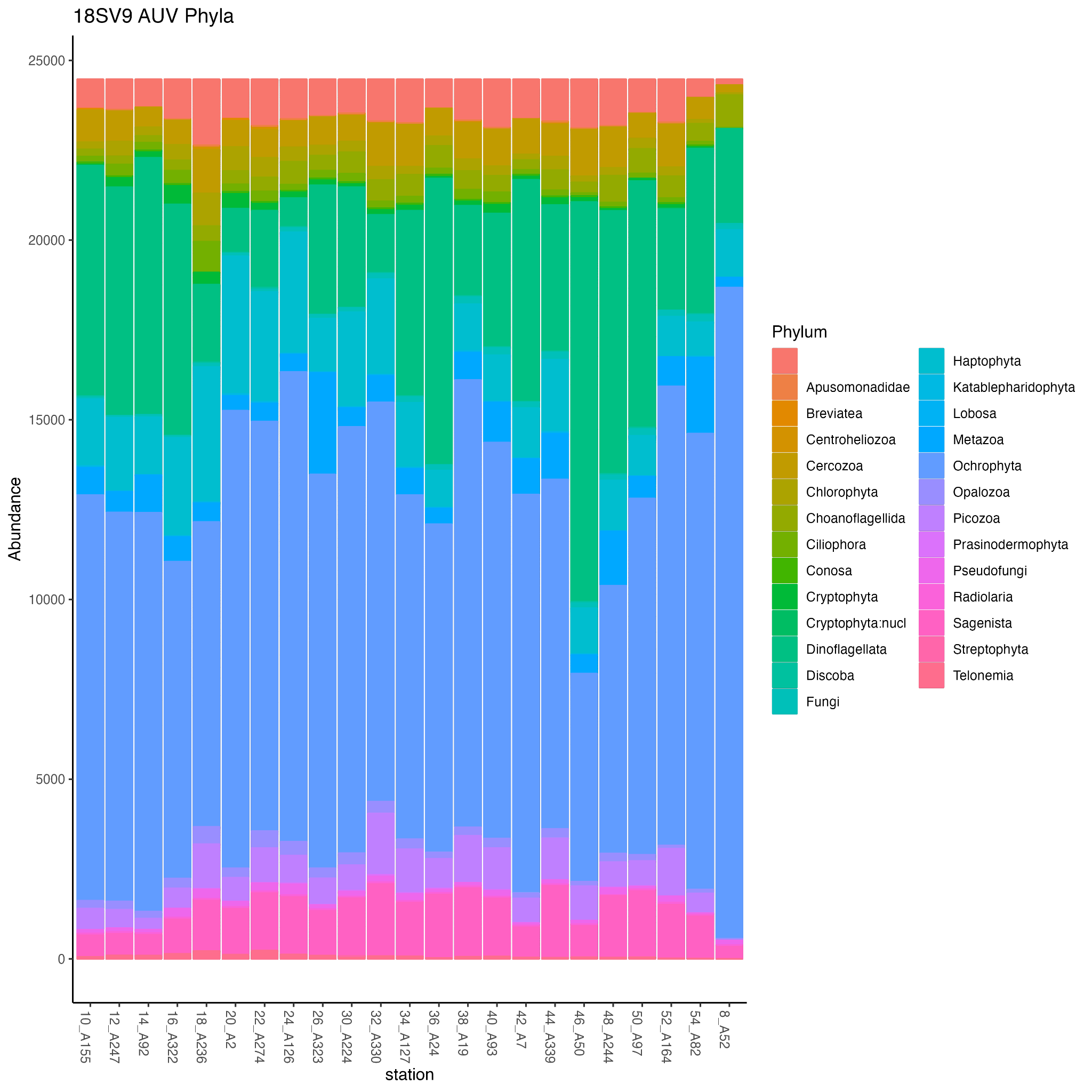

**Figure S7e:** 18SV9 metabarcoding from samples collected by the 3G-ESP. Relative abundance of phyla reveals synteny in the eukaryotic community over time and space.

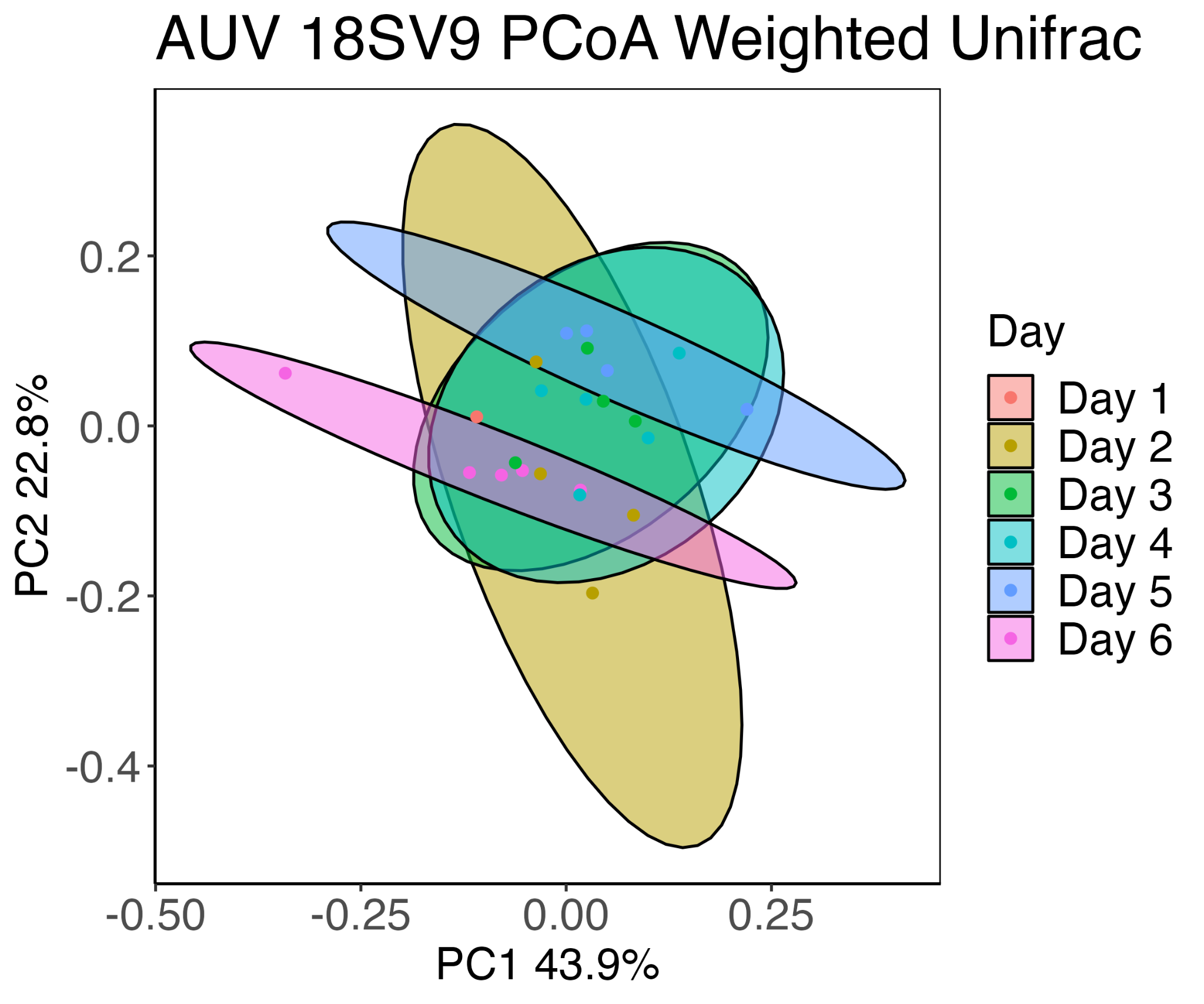

**Figure S7f:** 18SV9 metabarcoding from samples collected by the 3G-ESP. Principal Component Analysis reveals synteny in the eukaryotic community over time and space.

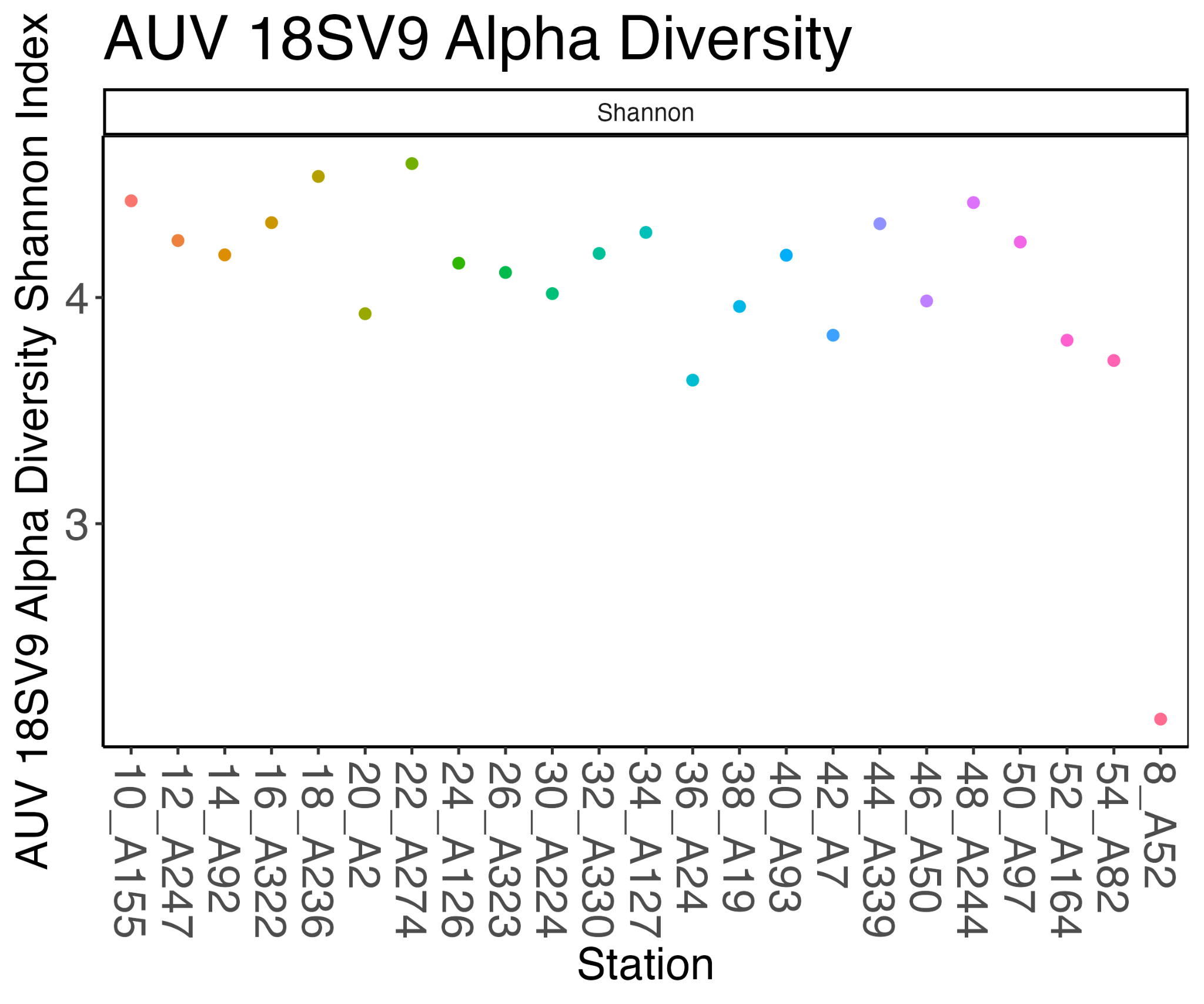

**Figure S7g:** 18SV9 metabarcoding from samples collected by the 3G-ESP. Alpha Diversity using the Shannon Index reveals synteny in the eukaryotic community diversity over time and space.

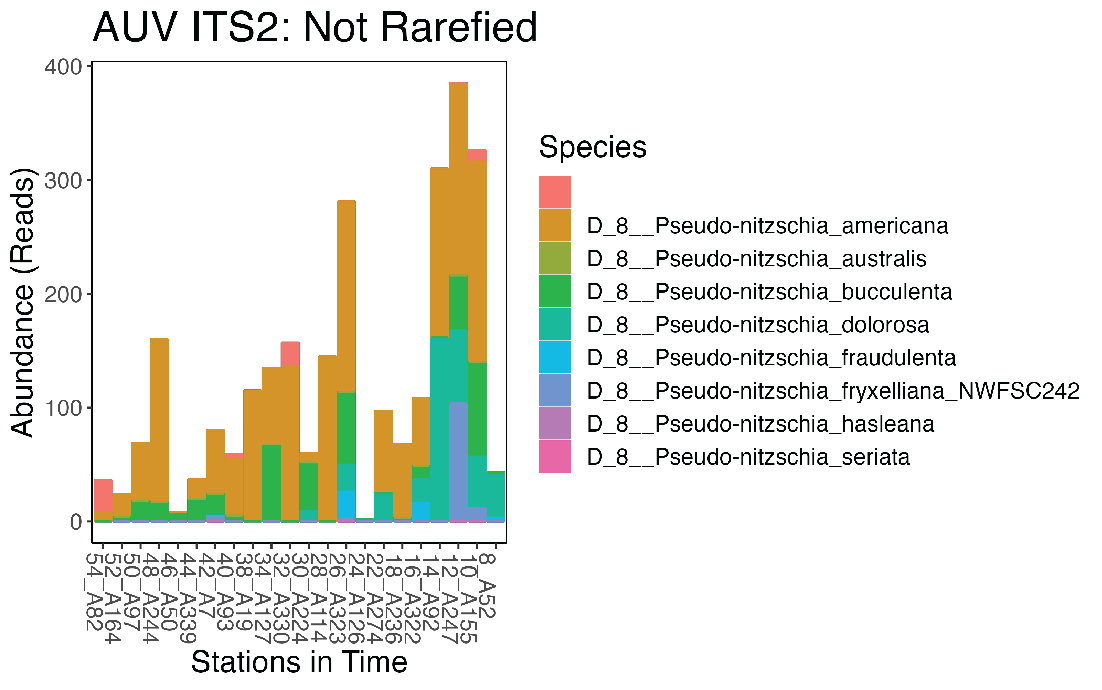

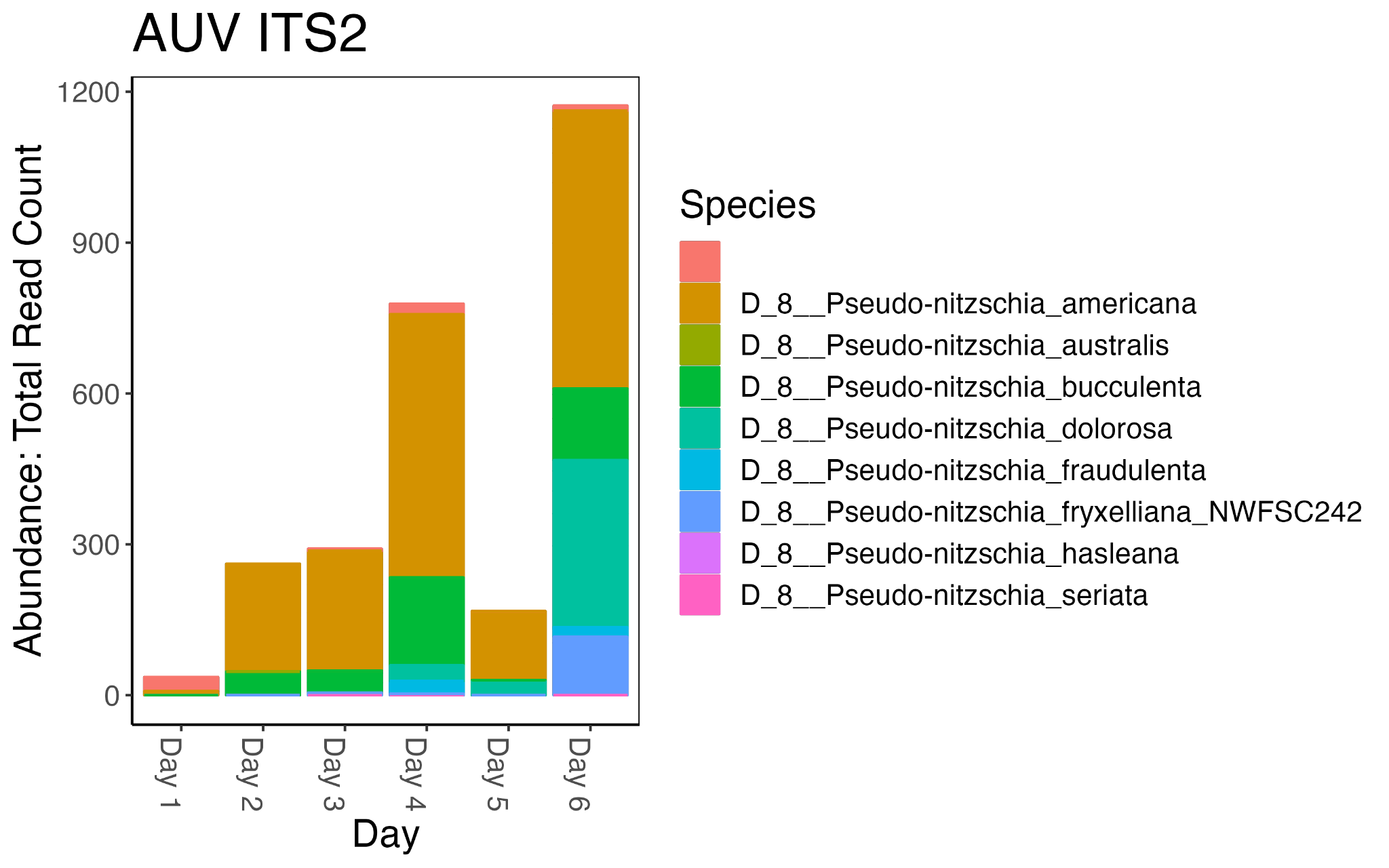

**Figure S7h:** ITS2 metabarcoding from samples collected by the 3G ESP. Read count over time and space (top) and binned by day (bottom) reveals 8 species of *Pseudo-nitzschia*. *Pseudo-nitzschia* abundance increases over time. Data was not rarefied due to low read count.

#### Table S1: Chlorophyll, Siex, Si:N

| *Depth_DepthClass_Station_Day* | *Chlorophyll_ug_L* | *Siex* | *Si:N Ratio* |
| --- | --- | --- | --- |
| *170_Deep_E1_Day1* | *0.10826296* | *10.928* | *29.2205128* |
| *10_SCM_E1_Day1* | *1.88035665* | *7.4296* | *2.52624063* |
| *5_Shallow_E1_Day1* | *1.64958561* | *10.3638* | *1.52010749* |
| *37_Deep_N1_Day1* | *2.36469094* | *4.9988* | *37.9558824* |
| *10_SCM_N1_Day1* | *2.36469094* | *8.2056* | *3.33021807* |
| *5_Shallow_N1_Day1* | *1.13676107* | *0.4708* | *1.22148202* |
| *50_Deep_N2_Day1* | *0.54701284* | *1.9326* | *1.26602214* |
| *50_Deep_N2_Day1* | *0.54701284* | *1.9326* | *1.26602214* |
| *8_SCM_N2_Day1* | *1.87750762* | *8.6158* | *4.4076694* |
| *5_Shallow_N2_Day1* | *1.81482907* | *11.9718* | *57.9383886* |
| *50_Deep_S1_Day1* | *0.53276772* | *0.7284* | *1.23495537* |
| *8_SCM_S1_Day1* | *1.78633882* | *11.2428* | *5.29424618* |
| *5_Shallow_S1_Day1* | *2.64959346* | *12.5662* | *20.2685888* |
| *5_Shallow_S1_Day1* | *2.64959346* | *12.5662* | *20.2685888* |
| *25_Deep_S2_Day1* | *1.66952878* | *0.4782* | *1.21785061* |
| *10_SCM_S2_Day1* | *1.78633882* | *6.8808* | *5.17504333* |
| *5_Shallow_S2_Day1* | *1.46724799* | *7.9696* | *7.69519152* |
| *25_Shallow_W2_Day1* |  | *10.0018* | *3.00995295* |
| *10_SCM_W2_Day1* | *1.95728033* | *8.2198* | *2.30614991* |
| *5_Deep_W2_Day1* | *0.16809249* | *9.0308* | *1.48456012* |
| *170_Deep_E1_Day2* | *2.2364848* | *11.0156* | *5.98790079* |
| *11_SCM_E1_Day2* | *2.73506422* | *7.9144* | *1.41046937* |
| *5_Shallow_E1_Day2* | *2.39318119* | *6.614* | *6.78600406* |
| *50_Deep_E2_Day2* | *0.65812483* | *1.164* | *2.10035751* |
| *12_SCM_E2_Day2* | *2.2364848* | *7.0516* | *1.2518601* |
| *5_Shallow_E2_Day2* | *2.82053497* | *8.7634* | *2.63709413* |
| *35_Deep_N1_Day2* | *2.19374942* | *-0.2632* | *2.33978583* |
| *11_SCM_N1_Day2* | *2.39318119* | *5.5348* | *1.18812864* |
| *5_Shallow_N1_Day2* | *2.30771043* | *6.237* | *3.40778761* |
| *40_Deep_N2_Day2* | *2.90600573* | *0.6558* | *2.78569624* |
| *8_SCM_N2_Day2* | *2.26782408* | *6.2746* | *7.75075377* |
| *5_Shallow_N2_Day2* | *8.09123164* | *6.518* | *1.22919078* |
| *55_Deep_S1_Day2* | *0.46154209* | *2.0322* | *1.72802713* |
| *17.5_SCM_S1_Day2* | *5.41314793* | *5.9176* | *1.80884908* |
| *5_Shallow_S1_Day2* | *2.25642798* | *4.5548* | *1.27553524* |
| *65_Deep_S2_Day2* | *0.27635545* | *2.3726* | *1.32401213* |
| *18_SCM_S2_Day2* | *1.97152546* | *4.8512* | *1.67264225* |
| *5_Shallow_S2_Day2* | *2.1681082* | *6.5572* | *2.19820368* |
| *65_Deep_W2_Day2* |  | *1.2188* | *1.47563805* |
| *15_SCM_W2_Day2* | *2.04844914* | *3.6828* | *2.72273355* |
| *5_Shallow_W2_Day2* | *2.39318119* | *6.6178* | *1.28209619* |
| *170_Deep_E1_Day3* | *0.3048457* | *1.5364* | *1.27971774* |
| *10_SCM_E1_Day3* | *2.01141181* | *6.9314* | *2.22000905* |
| *5_Shallow_E1_Day3* | *2.2364848* | *9.0128* | *2.09808241* |
| *85_Deep_E2_Day3* | *0.13105516* | *2.2828* | *1.89738503* |
| *12_SCM_E2_Day3* | *0.15384736* | *3.5042* | *1.41739562* |
| *5_Shallow_E2_Day3* | *2.07124134* | *7.734* | *1.30558254* |
| *35_Deep_N1_Day3* | *4.58693061* | *-0.9988* | *1.13494855* |
| *35_Deep_N1_Day3* | *4.58693061* | *-0.9988* | *1.13494855* |
| *12_SCM_N1_Day3* | *2.145316* | *4.3082* | *1.66304815* |
| *5_Shallow_N1_Day3* | *2.59261296* | *5.6774* | *6.0400682* |
| *62_Deep_N2_Day3* | *0.34758108* | *-25.1878* | *0.65761537* |
| *14_SCM_N2_Day3* | *1.6040012* | *7.8032* | *2.27201539* |
| *5_Shallow_N2_Day3* |  | *9.6672* | *6.21150855* |
| *55_Deep_S1_Day3* |  | *0.4906* | *1.47150765* |
| *10_SCM_S1_Day3* | *9.31631249* | *3.3352* | *1.87619942* |
| *5_Shallow_S1_Day3* | *1.98861961* | *6.4834* | *1.22705714* |
| *85_Deep_S2_Day3* | *0.26495935* | *2.5358* | *1.63409007* |
| *10_SCM_S2_Day3* | *1.82052712* | *4.7038* | *1.32282282* |
| *5_Shallow_S2_Day3* | *2.22223968* | *9.3774* | *7.21887035* |
| *75_Deep_W2_Day3* | *0.28490252* | *3.2272* | *1.31729301* |
| *28_SCM_W2_Day3* | *2.07978841* | *-1.365* | *1.14474803* |
| *5_Shallow_W2_Day3* | *1.87180957* | *7.08* | *2.16919918* |

#### Table S2: MZMine 3 Parameters

| Job name | EcoHAB 2021 |
| --- | --- |
| Mass Detection |  |
| MS level 1 |  |
| mass detector | centroid |
| noise level | 5.00E+04 |
| MS level 2 |  |
| mass detector | centroid |
| noise level | 1.00E+03 |
| Chromatogram Builder (ADAP) |  |
| MS Level 1 |  |
| min # data points for a peak | 3 |
| min feature height | 1.00E+05 |
| scan to scan m/z tolerance | 10 ppm |
| feature to feature m/z tolerance | 3 ppm |
| sample to sample m/z tolerance | 5 ppm |
| Max peaks in chromatogram | 15 |
| Min samples per aligned feature | 2 |
| Chromatogram Deconvolution |  |
| Algorithm |  |
| Chromatographic Threshold | 86.40% |
| Search minimum in RT range(min) | 0.05 |
| Minimum relative height | 0.00% |
| Minimum absolute height | 1.00E+05 |
| Min ratio of peak top/edge | 1.4 |
| Peak Duration range (min) | 0 |
| Isotope Peak Grouper |  |
| m/z tolerance | 0.0015 m/z or 3 ppm |
| Retention time tolerance | 0.05 min |
| Maximum Charge | 2 |
| Representative Istope | Most intense |
| Join Aligner |  |
| m/z tolerance | 0.0015 m/z or 5 ppm |
| Weight for m/z | 3 |
| Retention time tolerance | 0.1 |
| Weight for RT | 1 |
